# Distractor-specific control adaptation in multidimensional environments

**DOI:** 10.1101/2023.09.04.556248

**Authors:** Davide Gheza, Wouter Kool

## Abstract

Goal-directed behavior requires humans to constantly manage and switch between multiple, independent, and conflicting, sources of information. Conventional cognitive control tasks, however, only feature one task and one source of distraction. Therefore, it is unclear how control is allocated in multidimensional environments. To address this question, we developed a multi-dimensional task-set interference paradigm, in which people need to manage distraction from three independent dimensions. We use this task to test whether people adapt to prior conflict by enhancing task-relevant information or suppressing task-irrelevant information. Three experiments provided strong evidence for the latter hypothesis. Moreover, control adaptation was highly dimension-specific. Conflict from a given dimension only affected processing of that same dimension on subsequent trials, with no evidence for generalization. A new neural network model shows that our results can only be simulated when including multiple independent conflict-detector units. Our results call for an update to classic models of cognitive control, and their neurocomputational underpinnings.

## Introduction

From the ever-present lure of phone notifications to the cacophony of urban traffic, daily life requires us to manage many competing sources of information. This poses a challenge to our ability for *attentional control*. To achieve goals, we have to attend to sources that are relevant to our task, but we also have to determine how to process other sources. For example, at a talk, you should focus on the speaker, but complete suppression of the surroundings may lead you to miss the insightful observation your neighbor whispers in your ear. Even though we face a constant need for multidimensional attentional control, most empirical work has considered situations with a single source of distraction(Lindsay & Jacoby, 1994).

The notion of dimensionality is an important construct in theories of attentional control. Recent theoretical frameworks have drawn attention to the fact that attentional control needs to consider a wide variety of inputs(Shenhav et al., 2013), and outputs(Alexander & Brown, 2011). Moreover, it needs to configure control settings that are multivariate in nature(Ritz et al., 2022). It has also been suggested that the geometric nature of neural encoding of control settings can be best understood in terms of representational dimensionality(Badre et al., 2021; Fu et al., 2022). Yet, little is known about how attentional control is regulated in more complex environments with multiple sources of distraction(Pylyshyn & Storm, 1988). Here, we combine a new multidimensional attentional control task with computational modeling to address this question.

Psychologists study attentional control using response interference paradigms. In these tasks, one stimulus dimension signals the correct response, while a second one provokes a response tendency that is either congruent or incongruent with the former. For example, in the Stroop task(Stroop, 1935) participants name the color in which a word is printed. Responses on incongruent trials (e.g., “green” printed in red ink) are slower and less accurate, indicating a need for attentional control(Cohen et al., 1990; MacLeod, 1991). In other prominent tasks, conflict is achieved by presenting a target stimulus flanked by distractors(Eriksen & Eriksen, 1974) (Flanker task), or by manipulating whether the correct response and the irrelevant spatial location of a stimulus are on the same side(Simon & Rudell, 1967) (Simon task).

Data from these tasks demonstrate that humans configure attentional control based on prior demands. Specifically, people become less susceptible to distractors after experiencing conflict, a phenomenon known as the congruency sequence effect(Gratton et al., 1992). This effect is commonly interpreted to reflect an online adjustment of control strategies(Gratton et al., 1992), but its mechanisms are still debated. Some argue that prior conflict triggers increased processing of task-relevant information(Egner & Hirsch, 2005), but others have suggested that it results in suppression of distractor information(Danielmeier et al., 2011; Ritz & Shenhav, 2023). These two forms of control adjustments have been argued to produce similar behavior, because enhanced target processing and suppressed distractor processing should generally increase evidence in favor of the correct response(Botvinick et al., 2001).

Most models of the congruency sequence effect do not commit to a specific form of adaptation. According to the most influential account by Botvinick and colleagues(Botvinick et al., 2001), conflict yields both increased target processing and decreased attention to the distractor. Building on this model, Egner(Egner, 2008) proposed the existence of two independent conflict-control loops to explain adaptation during a task that combined Stroop and Simon interference. One of these loops detects stimulus-level conflict (i.e., between representations of color) and the other response-level conflict (i.e., between response options). One loop affects target processing while the other suppresses distraction, resulting in a mixture of both. Finally, a class of models proposes that conflict changes the connections between stimulus and task representations(Blais et al., 2007; Verguts & Notebaert, 2008). For example Verguts and Notebaert proposed a Hebbian-learning model of adaptation(Verguts & Notebaert, 2008), by which increased conflict strengthens associations between task and stimulus representations, also implementing both enhancement and suppression.

Unfortunately, there is a lack evidence to adjudicate between these theories. Classic interference paradigms are unable to address this question, because target and distractor processing are experimentally entangled. Neuroimaging research has also not provided a definitive answer. Early work suggested conflict results in enhanced processing of task-relevant information in posterior regions(Egner & Hirsch, 2005), but suppression of distractor activity has also been reported(Danielmeier et al., 2011; Padmala & Pessoa, 2011). This heterogeneity in findings may derive from the complex or imprecise relationships between neuroimaging techniques and cognitive mechanisms(Ritz & Shenhav, 2023). More recently, Ritz and Shenhav(Ritz & Shenhav, 2023) developed a behavioral task in which target and distractor information varied parametrically, and found that control adaptation affects distractor processing. However, they only used one distractor and one target dimension, and so it remains unclear how control adapts in multidimensional tasks. Finally, the multi-source interference task (Bush & Shin, 2006) introduced two sources of distraction by combining Flanker and Simon interference effects. However, these types of interference elicit qualitatively different conflict signals and engage separable conflict-control neural mechanisms of adaptation(Fu et al., 2022; Sheth et al., 2012), leaving unanswered how control is adjusted in response to multivariate conflict of the same type.

There are several distinct ways in which control adaptation can be implemented in multidimensional environments. If it affects distractors, there are two possibilities. Distractor adaptation could act at a global level, or it could be managed individually for each source of irrelevant information. In the case of global distractor adaptation, conflict from any source reduces all subsequent distractor processing. This algorithm, which is part of classic models of conflict monitoring(Botvinick et al., 2001; Egner, 2008), is efficient because it requires just one conflict monitor (or one for each conflict “type”(Egner, 2008)). In contrast, in the case of distractor-specific adaptation, conflict from a given source only reduces subsequent processing of the same distractor. This is less efficient but more optimal. For instance, it could suppress one (previously incongruent) distractor, while enhancing another (previously congruent) one. In addition, prior conflict may trigger target processing, simplifying the issue of how to allocate control.

Interestingly, some research suggests that control adaptation is conflict-type specific(Braem et al., 2014). When participants perform a task with two types of interference (e.g., Simon and Stroop), conflict from one source of distraction only reduces susceptibility to that distraction. These findings are revealing but limited, because these two sources yield types of conflict that require different forms of adaptation. For example, resolving interference stemming from the side on which the stimulus is presented (e.g., response conflict in a Simon task) requires qualitatively different changes in attentional control than adapting to conflict from word reading (stimulus conflict in a Stroop). It remains unknown how conflict is resolved when multiple distractors generate conflict of the same type (i.e., at the same stimulus or response level).

To address these questions, we developed a multi-dimensional task-set interference paradigm, based on a task by Liston and colleagues(Liston et al., 2006). On each trial, participants attend to one of four stimulus dimensions, with the remaining dimensions acting as distractors. Each of these three distractors is either compatible or incompatible with the correct response, resulting in a parametric manipulation of congruency. Like prior tasks, participants need to attend to a single cued dimension. However, the automaticity of processing non-cued dimensions is not driven by long-term training or exposure (as the case with word reading in the Stroop task), but because each dimension periodically signals the correct response(Hazeltine et al., 2011; Monsell et al., 2003). Critically, the interference generated from each dimension plays out at the same (stimulus) level.

The task allows us to measure not only the parametric effect of conflict on performance but also the individual contribution of each distractor dimension. Most importantly, we can use it to measure how control adapts in multidimensional environments. Because each dimension periodically acts as a distractor, the task allows us to determine how conflict from one dimension triggers attentional control adaptation towards itself and all other dimensions. This provides a particularly granular insight into how humans configure attentional control.

Across three task versions, we find striking evidence for dimension-specific control adaptation. Whenever a dimension generated conflict, it only affected subsequent processing of the same dimension. This is not compatible with a control mechanism that enhances task relevant information or with one that suppresses distraction globally. Rather, this suggests that conflict-driven control adaptation operates in parallel on task-irrelevant information pathways(Lindsay & Jacoby, 1994). Using computational modeling, we show that these results are incompatible with classic models, in which a single monitoring unit detects conflict at the response level (where evidence is aggregated across dimensions). Instead, we find that our results can be captured by including multiple independent conflict detectors for each dimension. This work improves our understanding of how people regulate attentional control in complex environments and provides experimental paradigms and a computational model for studying it.

## Results

We conducted three experiments, each with a different variant of the MULtidimensional Task-set Interference paradigm (MULTI; Figure 1A). In Experiment 1, each trial included two stimuli, presented side-by-side. These stimuli differed across four dimensions: their color (blue or orange), shape (oval or rectangle), edge (fully bounded or dashed), and motion of dots inside the object (upward or downward). A centrally located cue dictated which dimension was relevant on a given trial (C for color, S for shape, E for edge, or M for motion). For each of the four cues, participants were told to select the object with a particular target feature (left or right key press). For example, a participant could have been instructed to find the blue object if the task cue was C, the oval when the cue was S, the object with the fully bounded edge if the cue was E, and the object with the upwards moving dots if it was M. Target features were randomly assigned for each participant and remained stable throughout the entire session. The sequence of cues was pseudorandomly generated, cycling through each of the four tasks in random order with each cue repeating between 3 to 5 times (Figure 1C).

**Figure 1.**
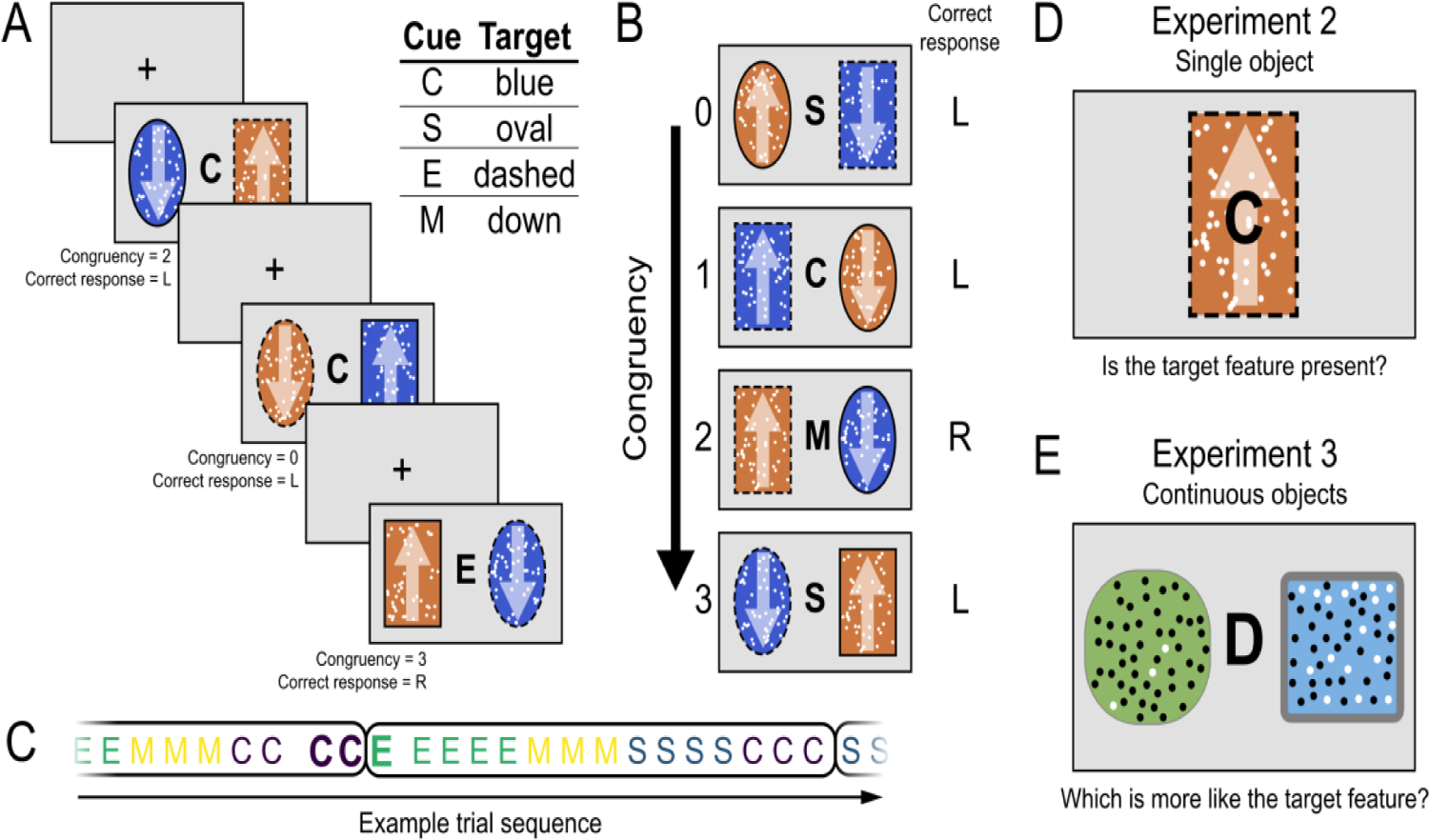
MULti-dimensional Task-set Interference task. **a**, Sequence of three example trials. On each trial, two objects are presented side-by-side that differ across four dimensions: the color (orange or blue), the shape (oval or rectangular), the edge (solid or dashed), and the direction of the dot motion (up or down; symbolized by white arrows, not shown in the experiment). Participants are instructed to pay attention to only one ‘task’ dimension cued by the letter between the stimuli. For each dimension, participants need to find a target feature, as instructed at the beginning of the experiment, and select the corresponding object (left or right response). In the example reported here, the target features are the blue color, the oval shape, the dashed edge, and the downward motion. All stimuli remain on screen until a response is given. Response deadline is set to 1.5s, and the inter trial interval to 1s, during which a fixation cross is presented. **b,** Each non-cued dimension is congruent when it primes the same side as where the cued dimension’s target feature is located. Therefore, congruency is parametrically manipulated from level 0 (all three non-cued dimensions prime the wrong side) to level 3 (all three non-cued dimensions prime the correct side). **c,** Example trial sequence in which the three example trials in panel **a**, represented by the cue letters C C E in bold, are embedded. Sequences are pseudo-randomly generated, with cued dimensions repeating 3 to 5 times, before a switch trial occurs. All dimensions are cued within 4 switches (encircled sequences), in random order. We refer to a sequence of repeat trials as ‘block’ (color coded sequences). **d, e,** Design of Experiments 2 and 3**. d,** In Experiment 2 a single central stimulus was presented at the center of the screen, consisting of a random sample of each of the four dimensions used in Experiment 1. The task cue was presented in the middle of the stimulus. At the beginning of the experiment, participants were given a target feature for each of the four dimensions. On each trial, they report whether the target feature of the cued dimension was present or absent. **e,** In Experiment 3, stimuli varied continuously on each dimension. Two objects are presented side-by-side, flanking a cue letter. Across trials, the two objects continuously differed in color (blue to green), shape (circle to square), edge (thick to thin), and the ratio of black to white dots (dots were static unlike Experiment 1 and 2). At the beginning of the experiment, participants were instructed to find a prototypical target feature, for each dimension (e.g., the more circular shaped). On each trial, participants reported which object was closer to the instructed feature.

Because features are randomly assigned to the stimuli, trials naturally involve different levels of congruency (Figure 1B). Any of the three non-cued dimensions could be congruent if it primed the correct response (i.e., the target features of the cued and a non-cued dimension are on the same side), or incongruent if it primed the opposite response. Therefore, the level of congruency varied from 0 (fully incongruent) to 3 (fully congruent). By independently varying the congruence of each non-cued dimension, the task allows us to comprehensively investigate congruency sequence effects both within and across non-cued dimensions. We provide access to all task code, data, analysis scripts, and simulation code, on the OSF (https://osf.io/j9fxc/).

### The number of congruent dimensions predicts task performance

The data from Experiment 1 revealed the MULTI imposes parametric control demands on participants. We found that the number of congruent dimensions predicted both response time and accuracy (Figure 2). Participants were fastest when the stimulus was fully congruent, and became slower with each decrease in the number of congruent dimensions (M_3_ = 637 ms, M_2_ = 655 ms, M_1_ = 674 ms, M_0_ = 694 ms). Analogously, participants showed highest accuracy for fully congruent stimuli, which decreased with each increase in the number of congruent dimensions (M_3_ = 98.6%, M_2_ = 97.5%, M_1_ = 95.3%, M_0_ = 91.4%). For both measures, a generalized Bayesian linear regression (see Methods) revealed a linear effect of congruency (ERs_congruency_ = Inf), which was corroborated by comparison of scores on adjacent congruence levels (range ERs: min = 13.5, max = Inf).

**Fig. 2.**
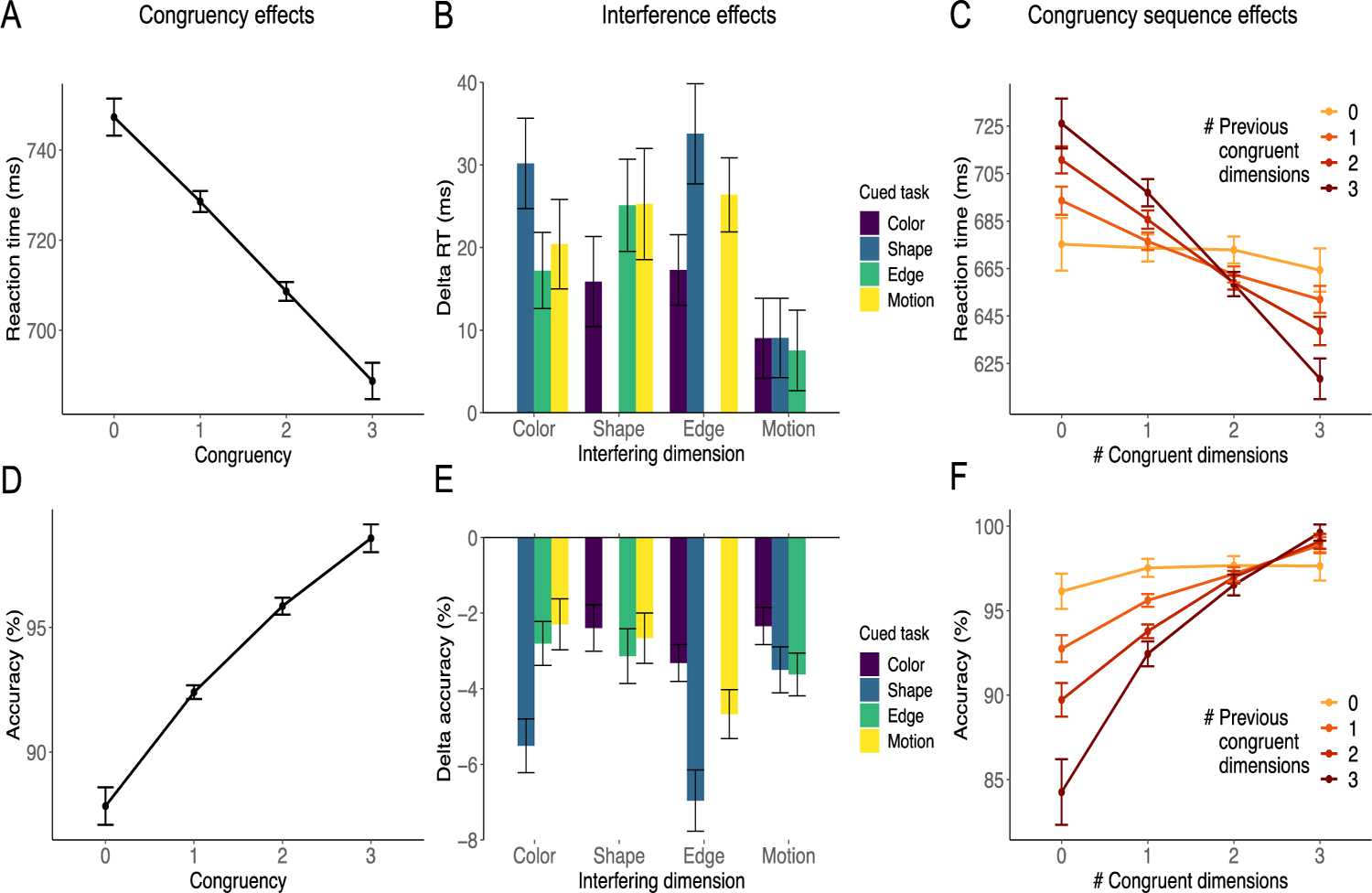
Experiment 1 (*N* = 104). Reaction time results in top row, accuracy results in bottom row. **a,d,** Parametric congruency effects. Participants responded faster and more accurately when the congruency level increased. **b,e,** Each non-cued dimension (x axis) generated a reliable interference effect (performance difference between trials in which the non-cued dimension is incongruent vs. congruent) on any cued task (color coded). **c,f,** Parametric congruency sequence effects: The effect of congruency on the current trial (x axis) is moderated by the congruency level of the preceding trial (color coded). Participants are less sensitive to the current congruency level after experiencing a low level of congruency in the previous trial, as indicated by flatter lines (lighter colors). Steeper lines (darker colors) reflect both stronger facilitation of the correct response by congruent non-cued dimensions, as well as stronger interference from incongruent non-cued dimensions; both facilitation and inhibition effects are more pronounced following highly congruent trials, reflecting a relaxed attentional control. Error bars represent within-subject 95% confidence intervals.

### Each stimulus dimension produces interference

A key benefit of MULTI is that it allows for the measurement of interference effects (the difference between congruent and incongruent trials) for each non-cued dimension (Figure 2BE). All non-cued dimensions produced interference for both response time (color: BF = 1.4×10^19^, shape: BF = 1.8×10^17^, edge: BF = 5.9×10^23^, motion: BF = 3.3×10^5^), and accuracy (color: BF = 9.6×10^21^, shape: BF = 4.4×10^20^, edge: BF = 1.0×10^28^, motion: BF = 1.2×10^20^). In fact, we observed reliable interference from any non-cued dimension on any cued task (Figure 2BE) for both RT (min BF = 16, max BF = 3.9×10^16^) and accuracy (min BF = 8.6×10^5^, max BF = 5.1×10^23^).

These results indicate that the task produces interference from multiple distractor dimensions. However, one alternative explanation is that each participant was susceptible to interference from only one randomly selected dimension and indifferent to the others. At the group level, this would yield reliable interference for each dimension. Our results are inconsistent with this hypothesis, because on average between 3 and 4 non-cued dimensions induced a positive interference effect (RT: M = 3.40, SD = 0.70; accuracy: M = 3.75, SD = 0.48). If only one dimension provided interference per participant, we would expect 2.5 positive effects on average (1 from the single interfering dimension, and 1.5 from 3 effects hovering around zero). Indeed, the average number of positive interference effects was higher than this value for both RT (BF = 1.8×10^44^) and accuracy (BF = 2.6×10^78^).

These findings suggest that, even though there is some heterogeneity in their potency, the distractors in the MULTI paradigm are processed in parallel. In order to formally test whether the distractors in our task act independently, we ran four hierarchical Bayesian models (one for each cued dimension) explaining response times as a function of the non-cued dimensions’ congruency levels and their interactions. We found very strong evidence against interaction effects between every pair of non-cued dimensions, and across all four models (range ERs: min = 7.7, max = 320.6; median = 158.5). This shows that reaction times (RT) only depend on the linear sum of each dimension’s congruency effects, which is consistent with the idea that stimulus dimensions are processed in parallel(Egner et al., 2007; Pylyshyn & Storm, 1988; Rumelhart & McClelland, 1987). We further corroborated this conclusion by analyzing the temporal profile of interference within a trial using distributional analyses(Van Den Wildenberg et al., 2010). Although the profile of interference varied across dimensions, the cumulative interference profile supported a linear combination of each dimension’s effects (see Supplementary Information for more details).

Unlike classic response interference tasks, distractors in the MULTI are not more automatically processed than targets. This is intentional, because each dimension can act both as a target and a distractor. Therefore, it is possible that a dimension only produces interference right after it was cued(Wylie & Allport, 2000). Contrary to this hypothesis, the Supplementary Information reports an analysis that demonstrates that interference effects last for several blocks of trials after the last switch (see Supplementary Figures 1 and 2) for all Experiments reported here.

### Conflict adaptation is modulated by the degree of prior congruency

Having validated our task, we then investigated how attentional control is regulated during multidimensional interference. Specifically, we measured susceptibility towards distraction as a function of the degree of congruence in the previous trial. Classic interference paradigms show a congruency sequence effect, where participants show reduced interference after an incongruent trial(Kerns et al., 2004). In the MULTI, however, congruence varies parametrically, which allows us to test whether the congruency sequence effect covaries with the prior congruency level.

A generalized Bayesian linear regression (see Methods) revealed very strong evidence for an interaction between current and previous congruency on RT (ER = Inf). The marginal means in Figure 2C show that the effect of current congruency was strongest after fully congruent trials, and then parametrically declined with prior congruency. Indeed, we observed only anecdotal evidence for an effect of congruency when the previous trial was fully incongruent (ER = 3.2, M_00_ = 665 ms, M_03_ = 662 ms), but there was very strong evidence for an effect of congruency after fully congruent trials (ER = Inf, M_30_ = 724 ms, M_33_ = 615 ms). Compared to trials following fully incongruent trials, trials following fully congruent trials showed strong interference when current congruency was low (leading to longer RTs; M_30-00_ = 59 ms) and strong facilitation when current congruency was high (leading to shorter RTs; M_33-03_ = −46 ms).

Analogous congruency sequence effects were observed for accuracy (Figure 2F). There was strong evidence for an interaction between current and previous congruency (ER = Inf), with larger congruency effects as previous congruency increases. Here, there was a strong effect of congruency even when the previous trial was fully incongruent (ER = Inf, M_00_ = 96.2%, M_03_ = 98.3%), but this effect was larger after fully congruent trials (ER = Inf, M_30_ = 83.8%, M_33_ = 98.8%). Mirroring RTs, we also found interference and facilitation effects on accuracy (M_30-00_ = −12.3%, M_33-03_ = 0.5%).

The Supplementary Information describes a series of analyses that demonstrate that these adaptation effects persist over multiple trials, but that they do not survive a task switch (e.g., switching from the color to the shape task). That is, after a task switch, the attentional state ‘resets’, with prior congruency not affecting behavior.

### Conflict adaptation is selectively driven within and not across dimensions

Even though these results establish parametric conflict adaptation, they do not address its mechanism. One possibility is that conflict is measured across dimensions, and then applied to all distractors or the target dimension. Alternatively, conflict may be estimated separately for each non-cued dimension, and then resolved selectively within them.

To address this question, we fully leveraged the MULTI, estimating trial-to-trial conflict adaptation within and across non-cued dimensions. For each participant, we computed average adaptation effects for each of the sixteen possible combinations of non-cued dimensions. Some of these combinations assess within-dimension adaptation (e.g., the effect of previous color congruency on current color sensitivity), while the others assess across-dimension adaptation (e.g., the effect of previous shape congruency on current edge sensitivity). For each combination of “current” and “previous” dimension, we computed the adaptation effect as the difference between the interference effects after congruent and incongruent values on the previous dimension. Larger adaptation effects indicate an increase in sensitivity to a non-cued dimension after a trial with congruence on (the same or a different) non-cued dimension.

Our results showed a highly specific pattern of congruency sequence effects (Figure 3), with strong adaptation effects for within-dimension sequences (on the diagonal of the heatmap on panels A and C) and no adaptation effects for across-dimension sequences (off-diagonal). This pattern was observed for RTs (Figure 3AB) and accuracy (Figure 3CD). To test this, we compared each within-dimension adaptation effect to an average across-dimension adaptation effect. For reaction times, we observed strong evidence for larger within-dimension compared to across-dimension adaptation effects (BF_color_ = 2.9×10^32^, BF_shape_ = 1.2×10^18^, BF_edge_ = 4.2×10^9^, BF_motion_ = 192). When considering across-dimension effects by themselves, we found some evidence favoring the null hypothesis, suggesting an absence of any adaptation effect (BF0_color_ = 9.2, BF0_shape_ = 0.7, BF0_edge_ = 8.3, BF0_motion_ = 8.9). Accuracy rates also showed larger within-compared to across-dimension adaptation effects (BF_color_ = 3.3×10^18^, BF_shape_ = 2.1×10^8^, BF_edge_ = 3.0×10^9^, BF_motion_ = 9.4×10^3^). Again, when considering the across-dimension effects by themselves, we found some evidence favoring the null hypothesis, suggesting an absence of adaptation effect for color and motion (BF0_color_ = 9.1, BF0_shape_ = 1.2, BF01_edge_ = 0.2, BF0_motion_ = 6.5).

**Fig. 3.**
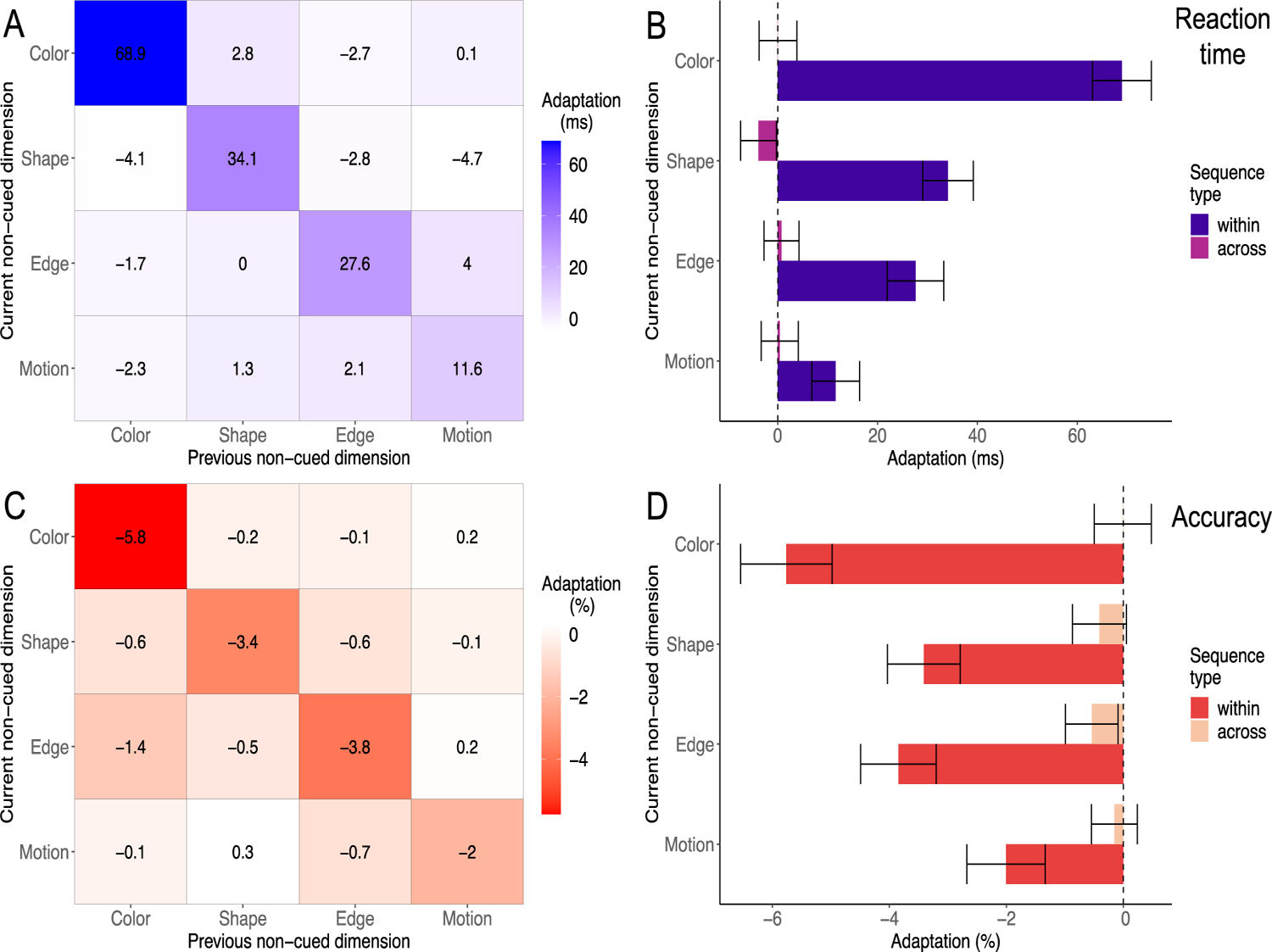
Distractor-specific conflict adaptation in Experiment 1. RT results in top row, accuracy results in bottom row. **a,c,** The heatmaps represent the trial-to-trial conflict adaptation effects measured in reaction time and accuracy, for each of the 16 possible sequences of previous and current non-cued dimensions. Diagonal cells correspond to within-dimension effects (e.g., the effect of previous color congruency on current color sensitivity), while off-diagonal cells correspond to across-dimension effects (e.g., the effect of previous color congruency on current shape sensitivity). The adaptation effects are defined as the difference between the current dimension’s interference effect after a congruent value on the previous dimension compared to after an incongruent value on the previous dimension. Adaptation effects were observed selectively within dimensions (diagonal), reflecting high specificity in control adaptation: Sensitivity to a current non-cued dimension only depends on its own congruency history. **b,d,** Average adaptation affects for sequences across dimensions (lighter colors, off-diagonal in heatmaps) compared to adaptation effects within dimensions (darker colors, on-diagonal cells in heatmaps). We found strong evidence for larger within-dimensions compared to across-dimensions effects. When considering across-dimensions adaptation effects by themselves, we found moderate evidence for absence of effects for color, edge, and motion dimensions. Error bars represent within-subject 95% confidence intervals.

This trial-average approach isolates adaptation effects for each combination of non-cued dimension, but it cannot account for cumulative effects of multiple non-cued dimensions on any given trial, as suggested by the parametric CSE results. In the Supplementary Materials, we report a series of Bayesian multilevel generalized linear models to explain within- and across-dimension adaptation effects at the single trial level. Their results were analogous to what is reported here.

Note that it is possible that each participant only applied within-dimension adaptation for one dimension, which then revealed itself as reliable within-dimension adaptation for all of them at the group level. To rule this out, we computed participants’ average number of positive within-dimension adaptation effects for both RT (M = 3.41 SD = 0.70) and accuracy (M = 3.39, SD = 0.67), and found strong evidence that these numbers were larger than the 2.5 positive adaptation effects one would expect under the alternative explanation (RT: BF = 2.2×10^44^, Accuracy: BF = 2.5×10^47^).

### Ruling out the role of eye movements

Because the MULTI presents two stimuli side-by-side, an alternative explanation is that our results are best explained by eye movements towards the stimulus with most target features, and not by attentional control. To rule out this explanation, participants in Experiment 2 (*N* = 104) completed a version of the MULTI in which each trial only presented a single stimulus, eliminating eye movement between objects (Figure 1D). The single central stimulus was constructed by randomly sampling one feature for each of the four dimensions, with the task cue presented in its middle. Participants reported whether the target feature of the cued dimension was present. We report the most important results here, but the Supplementary Information contains a more comprehensive description.

All results were replicated in this single-stimulus version of the MULTI (Figure 4). The number of congruent dimensions parametrically modulated task performance in terms of RT and accuracy. Non-cued dimensions produced interference both on the aggregate and when considered separately for each cued dimension. The dimensions’ individual interference effects did not depend on the congruency of the other non-cued dimensions, again showing that dimensions in the MULTI paradigm are processed in parallel.

**Fig. 4.**
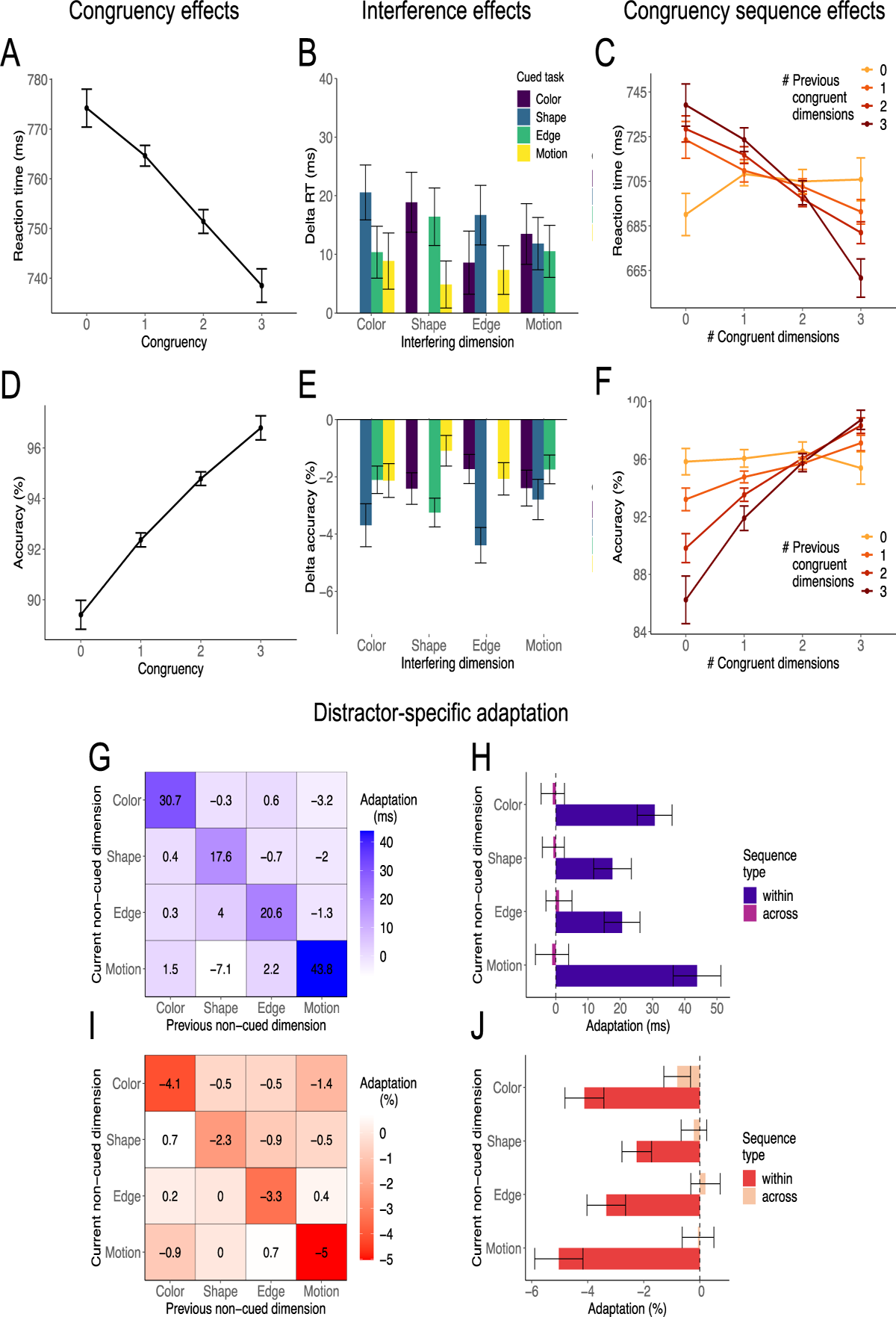
Experiment 2. (*N* = 104). RT results in top row, accuracy results in bottom row. Compare to Figure 2. **a,d,** Participants showed a parametric congruency effect with faster and more accurate responses when the congruency level was higher. **b,e,** Each non-cued dimension (x axis) generated a reliable interference effect on any cued task (color coded), except for a weak effect of the interfering dimension shape on the motion task (BF = 3.3). **c**,**f**, Parametric congruency sequence effects: Participants are less sensitive to the current congruency (x-axis) after experiencing a low level of congruency in the previous trial, as indicated by flatter lines (lighter colors). When previous congruency increases (darker colors), participants become sensitive to current congruency, with both facilitation (better performance for relatively congruent trials) and interference (worse performance for relatively incongruent trials), reflecting a relaxed attentional control. Error bars represent within-subject 95% confidence intervals. **g,i,** Heatmaps represent the trial-to-trial conflict adaptation effects on reaction times (**g**) and accuracy (**i**), for each of the 16 possible sequences of previous and current non-cued dimensions. Diagonal cells corresponds to within-dimension effects, while off-diagonal cells corresponds to across-dimension effects. Replicating the pattern found in Experiment 1, adaptation effects were observed selectively on diagonal cells, reflecting high specificity in control adaptation: Sensitivity to a current non-cued dimension only depends on its own congruency history. Notably, in this task with a single stimulus per trial, the dimension motion induced the most robust interference effect (Figure 5BE), mirrored by the strongest within-dimension congruency sequence effect. **h, i,** Average across-dimension sequence effects (lighter colors) compared to within-dimension adaptation effects (darker colors). For each dimension, we observed larger evidence for within compared to across dimension adaptation for both RT (BF_color_ = 1.5×10^13^, BF_shape_ = 1.6*10×^5^, BF_edge_ = 3.0×10^5^, BF_motion_ = 5.2×10^11^) and accuracy (BF_color_ = 3.5×10^8^, BF_shape_ = 9.8×10^5^, BF_edge_ = 2.5×10^9^, BF_motion_ = 8.0×10^12^). When we only considered across dimension adaptation effects, we observed some evidence favoring the null hypothesis, suggesting absence of adaptation for RT (BF0_color_ = 7.7, BF0_shape_ = 8.2, BF0_edge_ = 8.1, BF0_motion_ = 8.3) and accuracy (BF0_color_ = 0.01, BF0_shape_ = 4.9, BF0_edge_ = 6.3, BF0_motion_ = 8.9).

We again observed sequential conflict adaptation effects (Figure 4CF), with susceptibility to current congruency increasing when following more congruent trials (ER_RT_ = Inf, ER_accuracy_ = Inf). Most importantly, conflict was again resolved in a dimension-specific fashion, driven by conflict sequences within and not across dimensions (Figure 4G-J). Bayesian hierarchical regressions provided consistent results at the single-trial level (see Supplementary Information).

### A neural network model of dimension-specific adaptation

Classic computational models of cognitive control suggest that conflict is monitored after evidence from all dimensions have been merged into stimulus or response representations. However, such a mechanism is incompatible with our results, because conflict adaptation is sensitive to the source of conflict. This suggests that conflict monitoring occurs at a lower level in the information processing stream.

To demonstrate this, we developed a novel neural network model of our task based on prior models of classic interference paradigms(Cohen et al., 1990; Cohen & Huston, 1994; Ritz & Shenhav, 2023). An outline of the model is shown in Figure 5. The model processes stimulus information through four feed-forward pathways, corresponding to the four dimensions of the MULTI paradigm. Each pathway starts with two units that represent the location of the target feature of the corresponding dimension on the current trial. Their activation propagates to intermediate units, which also receive activation from four ‘task’ units that represent attention allocated to each dimension. Each task unit is connected to the two intermediate units in the pathway of the corresponding dimension and is activated when the corresponding dimension is cued. Their activation sensitizes processing in the intermediate units by offsetting a constant negative bias(Cohen et al., 1990). Each intermediate unit then propagates its activity to two response units. In order to simulate RTs and accuracy, each trial’s evidence (the difference in activation between the two response units) is sampled through a diffusion process(Ratcliff et al., 2016). We hand-tuned the parameters of the model, aiming to reduce complexity by equating them where possible. In the Supplementary Information, we show that the model effectively reproduces the essential behavioral patterns reported below across a broad parameter range.

**Fig. 5.**
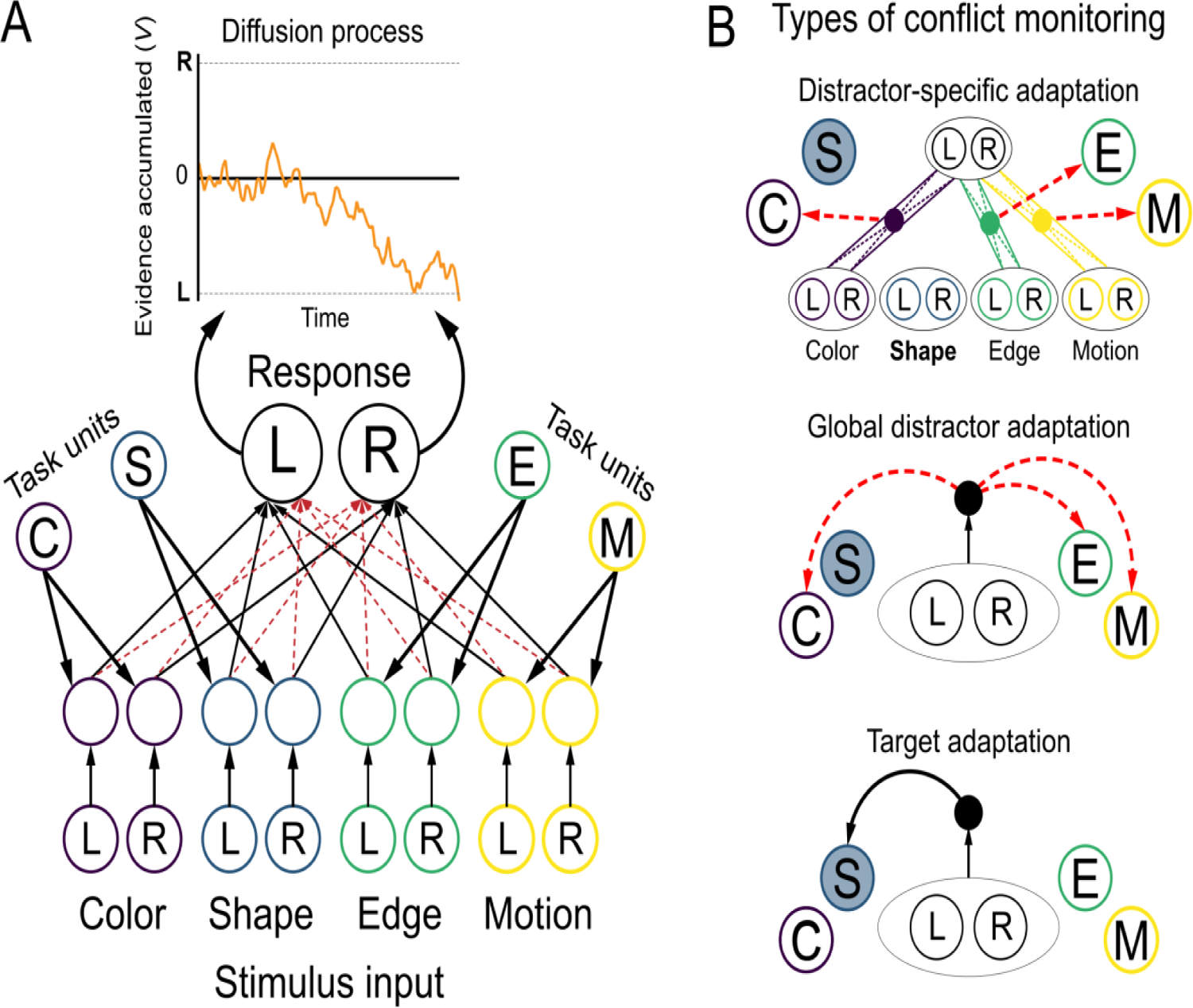
Computational model. **a**, Schematic description of the neural network model developed for the MULTI paradigm. (Bottom) The model processes stimulus information through four feed-forward pathways (color coded), corresponding to the four dimensions of the MULTI paradigm. Each pathway projects to a set of intermediate neurons, where stimulus input activity converges with activity from task units and a constant negative bias. The intermediate layer projects to two response units using balanced excitatory (black lines) and inhibitory connections (red dashed lines). (Top) In order to simulate RTs and accuracy, each trial’s evidence (the difference in activation between the two response units) is sampled through a diffusion process. **b**, Three variants of conflict-control loops. In all, conflict (solid circles) is transformed into a measure of control, which is used to update the activity of the task-demand units on the next trial (red dashed arrows: subtraction, black solid arrows: addition). (Top) Dimension-specific adaptation assumes that multiple conflict measures are computed within each processing pathway. Three conflict-monitoring units compute conflict as Hopfield energy within each dimension-specific pathway. Each dimension’s conflict is then transformed into a control value which is used to change activity in the corresponding task unit on the next trial. Because each dimension has a separate conflict-control loop, this mechanism allows for distractor-specific adaptation. (Middle) To implement global distractor adaptation, conflict is detected at the response level, where the processing pathways have fully converged. Specifically, the single conflict value is transformed into a control value and then subtracted from the input to all non-cued dimensions’ task demand units. (Bottom). For target adaptation, conflict is detected at the response level. This single conflict value is transformed into a control value, which is then added to the input to the cued dimension’s task demand unit.

The model captured single-trial-level performance well. Average RT decreased and accuracy increased when the number of congruent non-cued dimensions increased (Figure 6AB). This happens because on trials with more congruent dimensions, evidence in the response layer is more favorable towards the correct response, leading to a faster and more accurate resolution of the diffusion process.

Next, we aimed to simulate the distractor-specific conflict adaptation effects we observed in Experiments 1 and 2. To do this, we considered three variants of models that implement some form of conflict monitoring. In all, “monitoring” units measure the degree of conflict, or simultaneous activation of incompatible pieces of information (e.g., responses). This measure of conflict is then transformed into a measure of *control*, which is used to configure the activity of the task units on the next trial. Attentional control (target enhancement or distractor suppression) is increased after trials with a high degree of conflict(Botvinick et al., 2001; Egner, 2017; Gratton et al., 1992).

We simulated three models to demonstrate that distractor-specific congruency sequence effects require individual conflict monitors for each distractor. The key difference between these models is how they monitor conflict. The first detects conflict separately for each input dimension, whereas the last two models detect conflict at the response level.

Our first model assumed multiple conflict-monitoring units at the intermediate layer of the network (Figure 5B top). Here, we define conflict as the simultaneous activation of incompatible representations between a given processing pathway (i.e., at the stimulus representation level) and the response layer. Specifically, the model measures Hopfield energy for each dimension’s pathway as the coactivation of dimension-specific intermediate units and response units, weighted by their connections(Botvinick et al., 2001). The absolute value of this measure is largest when all units are maximally active, becomes zero if all units are inactive, and its sign depends on the configuration of connections’ weights. In terms of our model, energy within a pathway is high when the intermediate representation is not compatible with the response activations. The Hopfield energy for each pathway then determines the control value applied to the corresponding task unit on the next trial, in such a way that increased conflict from a given dimension leads to suppression of attention towards that dimension.

This model reproduced the parametric congruency sequence effects observed across all three experiments (see Figure 6, panels A & B). It also reproduced adaptation effects over longer timescales (see Supplementary Information) without the need for parameter changes (Jiang et al., 2014). More importantly, this model successfully captured dimension-specific congruency sequence effects. As can be seen in Figure 6C, adaptation effects were determined exclusively by congruency sequences within the same dimension, and not across dimensions – in full agreement with the empirical evidence.

**Fig. 6.**
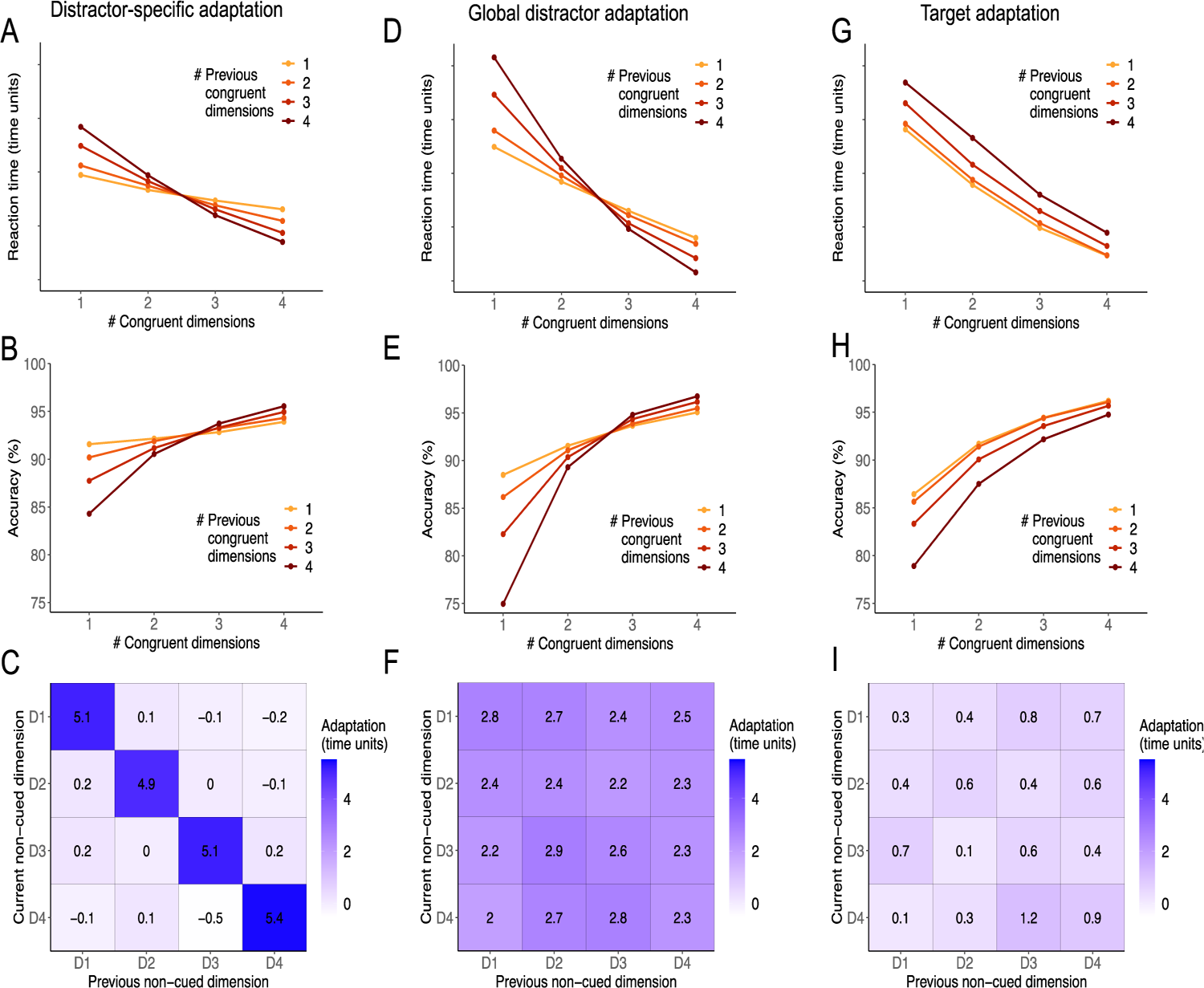
Simulation results of the three conflict adaptation models. **a,b,c** Distractor-specific adaptation reproduces the parametric congruency sequence effects observed across all three experiments, and yields dimension-specific congruency sequence effects: Adaptation effects are determined exclusively by congruency sequences within the same dimension (on diagonal cells). **d,e,f,** Global distractor adaptation reproduces the parametric congruency sequence effects, but fails to capture dimension-specific congruency sequence effects: Adaptation effects are determined both within- and across-dimensions. **g,h,I**, Target adaptation fails at capturing the parametric congruency sequence effects observed across the three experiments, and also the within-dimension specificity of this adaptation. However, it does yield generally increased performance after high-conflict trials (lower RT and higher accuracy), but these effects are orthogonal to the current congruency level. For accuracy, it may appear we observe a parametric sequence effect, since the slope of the curves change with prior congruency, but this reflects a linear shift in accuracy combined with a ceiling effect at the higher congruency levels.

To demonstrate that the existence of multiple conflict monitors is crucial for capturing our results, the last two models implement a single conflict detector at the level of the response nodes(Botvinick et al., 2001; Verguts & Notebaert, 2008). Specifically, we quantify response conflict as Hopfield energy in the response layer, or the product of response units’ activations, by assuming mutual inhibition between them. When the activation of one response unit is larger than the other, evidence is large and conflict is low. When the activity of the two units is similar, evidence is low and conflict is high. This single conflict signal is then applied to the task demand units following the logic described above.

For the second model, we moved conflict detection to the response level(Botvinick et al., 2001), but still used this to adapt distractor representations (Figure 5B middle). After conflict detection, the control value was subtracted from the input to all non-cued task units. This way, attention to all distractors reduces (increases) after particularly incongruent (congruent) trials, yielding global distractor adaptation. This model produced parametric congruency effects. For both accuracy and reaction time, the effect of congruency on a given trial was dependent on the congruency of the previous trial (Figure 6, panels D & E). However, the model did not produce dimension-specific adaptation. As shown in Figure 6F, adaptation effects were observed both for sequences within and across dimensions. This pattern arises because the system is agnostic to the source of the conflict. At the response layer, conflict reflects the cumulative contributions of all non-cued dimensions, making dimension-specific contributions unidentifiable.

The third model implemented conflict adaptation by enhancing the target representation (Figure 5 bottom). Specifically, after the conflict value is transformed into a control value, it is added to the input to the cued dimension’s task unit. This model failed to capture parametric congruency sequence effects (Figure 6GH). High-conflict trials induced increases in performance (lower RT and higher accuracy) on the next trial, but they did not change susceptibility to distractor information. Naturally, this model was also unable to capture the distractor-specific adaptation we observed in our experiments (Figure 6I).

This last set of simulations demonstrates that, at least in the gold-standard neural network model of attentional control(Cohen et al., 1990), congruency sequence effects cannot be achieved by modulating attention to the cued task. Instead, the first two simulations suggest that they only emerge when attention to the distractors changes from trial to trial. Importantly, target adaptation leaves unaffected the processing of the distractors^1^. Therefore, no changes in facilitation nor interference from non-cued dimensions are to be expected. Rather, target adaptation primarily changes the availability of evidence related to the cued dimension, leading to similar changes in accuracy and RT (i.e., in the signal-to-noise ratio of the target feature) independently of the current level of congruency.

In short, only our novel network was able to capture our behavioral results, due to a control mechanism that independently monitors conflict for each non-cued dimension.

### Ruling out associative priming and target adaptation

Our neural network simulations revealed that adaptation of attention towards target and distractor information have a qualitatively different profile. Distractor adaptation changes sensitivity towards the non-cued dimensions, but target adaptation simply increases performance on the cued task. Experiments 1 and 2 did not reveal such evidence for conflict-driven enhancement of target information in the form of faster responses after particularly incongruent trials. On the other hand, we observed a modest post-conflict increase in accuracy (which generalized across dimensions; see Supplementary Information). Taken alone, this effect is compatible with target adaptation, in combination with increased domain-general response caution(Ratcliff, 1978; Ratcliff et al., 2016) in response to conflict. Thus, it is possible that target adaptation was present but modest, and perhaps masked by an increase in response caution.

Therefore, we ran a third experiment (*N* = 103) in which we varied the evidence strength of each dimension across trials. We reasoned that, in the case of target adaptation, we should observe a change in sensitivity to the evidence in the cued task. Specifically, on each trial of this new MULTI paradigm, all features on each dimension were sampled from continuous ranges (Figure 1E). Objects continuously differed in color (blue to green), shape (circle to square), edge (thick to thin), and the ratio of black to white dots. Participants were asked to report which object was closer to the instructed feature (e.g., which is object is more circular?).

This experiment also allowed us to address a set of alternative explanations for our previous findings. It is often suggested that congruency sequence effects do not measure dynamics in attentional control, but instead reflect priming of associations between stimulus features or between stimulus and response(Hommel et al., 2004; Mayr et al., 2003; Schmidt & De Houwer, 2011). The MULTI mitigates some of these concerns because responses are not tied to a given feature: each target feature can appear left or right on any trial. Moreover, stimuli are unlikely to repeat from trial-to-trial because there are sixteen possible combinations of features. Contingency learning is a more serious confound affecting both experiments: among the sixteen potential stimuli, fully congruent and fully incongruent ones occur less frequently. Therefore, stimuli with more extreme congruency levels could be more strongly linked to certain responses. The design of Experiment 3 mitigates these concerns. Because each trial included two unique new features for each dimension, there was no possibility for associative priming and contingency learning to affect behavior. The Supplementary Information reports the results in full detail.

Experiment 3 replicated the previous two experiments. On each trial, the level of congruency affected performance for both RT and accuracy (Figure 7). Individual dimensions produced reliable interference effects for both performance measures, although the effect of the new dot proportion dimension was markedly reduced for RT. As before, individual interference effects did not depend on the congruency of the other non-cued dimensions, suggesting that dimensions were processed in parallel. We also again found strong evidence for conflict adaptation. On the aggregate, susceptibility to current congruency was increased following more congruent trials (ER_RT_ = Inf, ER_accuracy_ = Inf).

**Fig. 7.**
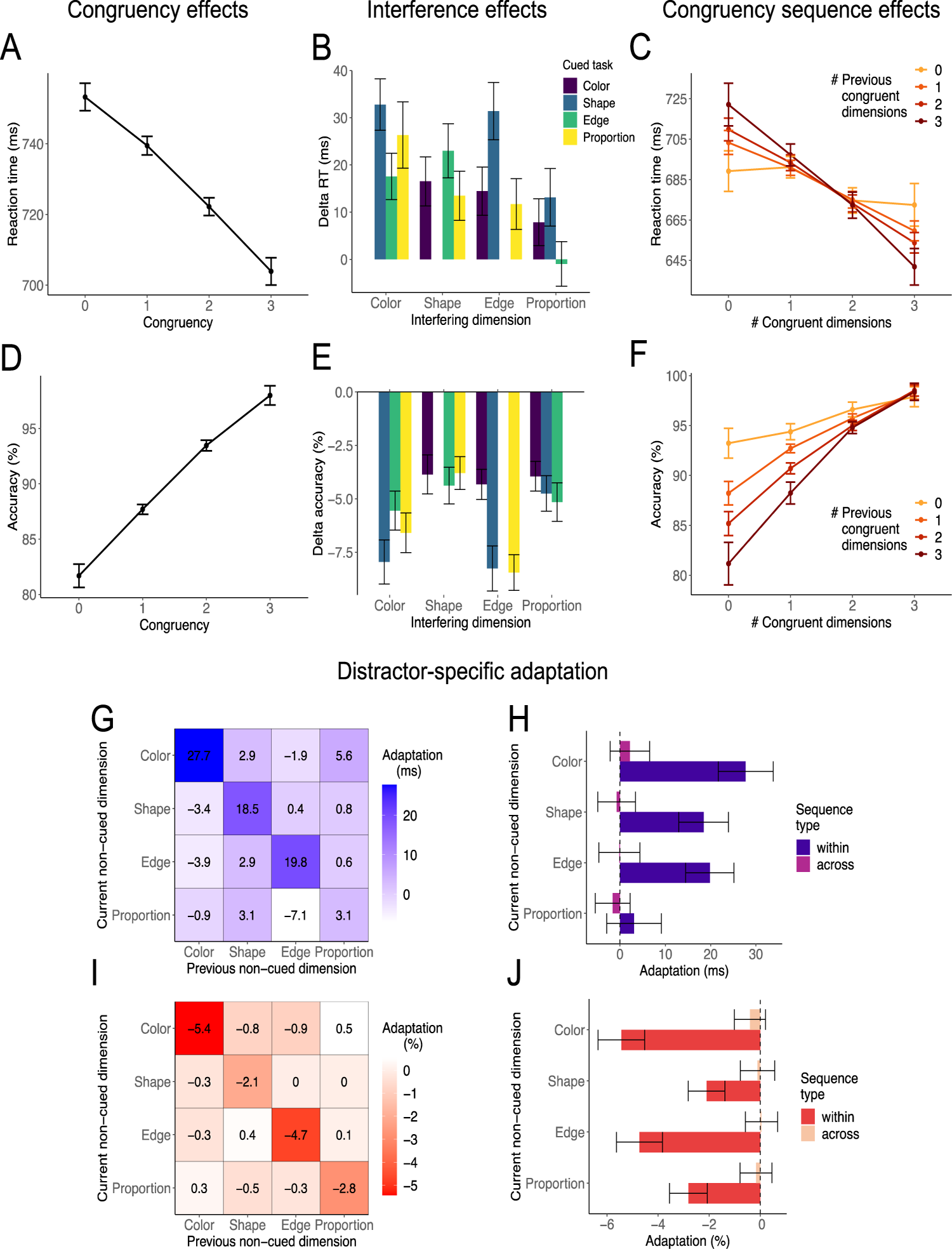
Experiment 3 (*N* = 103). RT results in top row, accuracy results in bottom row. Compare to Figures 2 and 3. **a,d,** Participants showed a parametric congruency effect with faster and more accurate responses when the congruency level was higher. **b,e,** Each non-cued dimension (x axis) generated a reliable interference effect on any cued task (color coded), except for the interfering dimension dot proportion on the edge task (BF = 0.08). **c**,**f**, Parametric congruency sequence effects: Participants are less sensitive to the current congruency (x-axis) after experiencing a low level of congruency in the previous trial, as indicated by flatter lines (lighter colors). Error bars represent within-subject 95% confidence intervals. **g,i,** Heatmaps represent the trial-to-trial conflict adaptation effects on reaction times (**g**) and accuracy (**i**), for each of the 16 possible sequences of previous and current non-cued dimensions. All effects replicated except for a weak within-dimension congruency sequence effect for the new dot proportion dimension, on RT selectively. **h,j,** For RT, we observed larger evidence for within compared to across dimension adaptation (BF_color_ = 6.7×10^7^, BF_shape_ = 1.6×10^5^, BF_edge_ = 8.5×10^4^), except for the new dot proportion dimension (BF_proportion_ = 0.4). For accuracy, there was strong evidence for larger within than across adaptation for all dimensions (BF_color_ = 3.8×10^11^, BF_shape_ = 605, BF_edge_ = 6.1×10^9^, BF_proportion_ = 2.4×10^4^). Note that this asymmetry in adaptation for dot proportion between measures is consistent with the weak interference effect on RT but not accuracy for this dimension (cf. panel B). We found moderate evidence for a lack of across-dimension adaptation for all dimensions for RT (BF0_color_ = 5.0, BF0_shape_ = 8.5, BF0_edge_ = 9.1, BF0_proportion_ = 6.7) and accuracy (BF0_color_ = 3.8, BF0_shape_ = 8.5, BF0_edge_ = 9.0, BF0_proportion_ = 7.8). Error bars represent within-subject 95% confidence intervals.

As before, adaptation effects were resolved selectively within each dimension (Figure 7, panels G-J) for accuracy. For RT, we observed larger evidence for within-compared to across-dimension adaptation except for the new dot proportion dimension. For accuracy, we observed larger evidence for within-compared to across-dimension adaptation for all dimensions. Note that the asymmetry in adaptation for dot proportion between measures is consistent with the weak interference effect on RT but not accuracy for this dimension. Moreover, we found moderate evidence for a lack of across-dimension adaptation for all dimensions for both RT and accuracy.

Next, we leveraged the fact that all dimensions varied in discriminability to investigate more subtle effects of target adaptation. We reanalyzed the data from a signal-detection perspective, defining each dimension’s signal as the absolute difference in intensity between left and right features (Figure 8). Thus, each non-cued dimension could now vary both in congruency and signal intensity. The novel prediction afforded by this approach is that conflict-driven changes in target processing would affect the target sensitivity. More precisely, model simulations of this continuous-object version of the task indicated that conflict-driven target adaptation (Figure 5B), but not distractor adaptation, leads to faster RT and higher accuracy, especially for high-target-signal trials (Figure 8F). We found that response speed was affected by previous signal congruency but not in the direction predicted by target adaptation. Responses were only slightly slower following more incongruent trials, in line with post-conflict slowing (RT: ER_prev_con<0_ = 40.6. Figure 8E). The target-adaptation simulations predicted an interaction between previous signal congruency and current target signal, but we found evidence against this effect (ER_prev_con:target=0_ = 101.5). On the other hand, we did observe higher accuracy following low signal congruency (ER_prev_con<0_ = Inf.; Figure 8G), as predicted by our simulations (Figure 8H). This pattern of increased accuracy and substantially invariant RTs suggests that, to some degree, people enhanced target processing following more incongruent trials, but became also more cautious. Specifically, it is plausible that faster evidence accumulation, resulting from increased target processing, was offset by heightened response thresholds (in line with the results of Experiments 1 & 2). Most importantly, the strongest signal-level adaptation effects confirmed distractor adaptation (Figure 8CD), which recapitulated the parametric congruency effects reported above (Figure 7). Specifically, signal congruency effects (ER_cur_con<0_ = Inf.) were moderated by previous signal congruency (ER_cur_con:prev_con<0_ = Inf.). These effects were orthogonal to target signal. In fact, adaptation effects on RTs were comparable along target signal intensity levels (ER_target:prev_con:cur_con=0_ = 53), and so were congruency effects (ER_target:cur_con=0_ = 256).

**Fig. 8.**
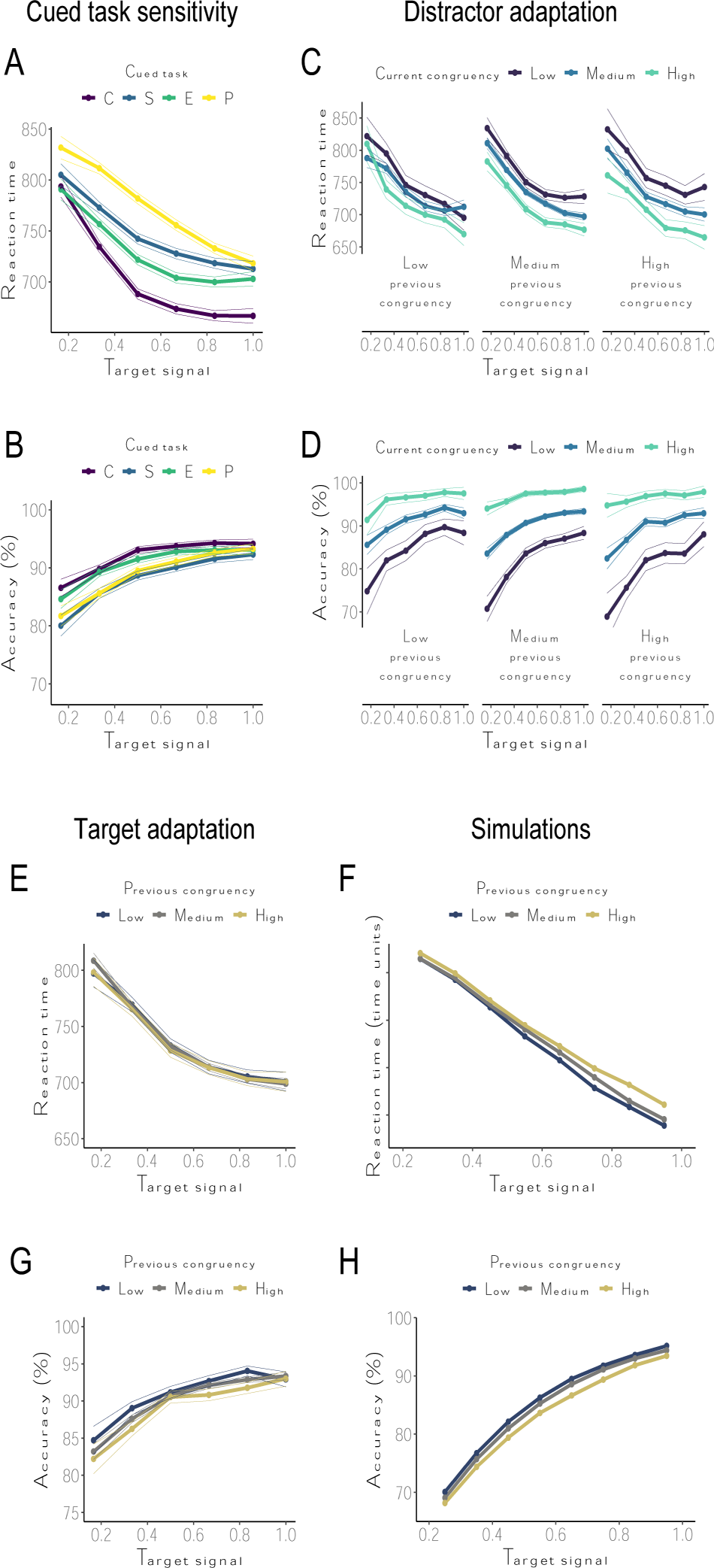
Experiment 3 (*N* = 103). Signal detection analyses on RT (top rows) and accuracy (bottom rows). Congruency is computed as the average *signed signal* across the non-cued dimensions. **a,** Participants responded faster to trials with larger target signal. **b,** Mean accuracy differed across cued tasks, but their psychometric curves were comparable. **c,d,** Signal congruency effects on RTs (c) and accuracy (d) were moderated by previous signal congruency, but were orthogonal to target signal. These signal-level adaptation effects confirmed a distractor-based adaptation, and recapitulated the parametric congruency sequence effects (Figure 7CF), as increased interference (facilitation) from incongruent (congruent) trials following high congruency in the preceding trial. **e-h,** Signal detection analyses highlighted some evidence for target adaptation. **e,f,** Reaction times were substantially unaffected by previous congruency, at odds with simulation results (f) of target adaptation in this continuous-object task. **g,h,** Nevertheless, low congruency was associated with higher accuracy in the following trial, as predicted by simulations (h). Overall, these results imply simultaneous adaptation effects on both target processing and response caution. Note that the stimulus features’ signals were randomly sampled from a uniform and common feature space, and that their absolute difference ranged from 0.2 to 1. This ensured above-chance target detection and distractor salience at the lowest level of discriminability.

Interestingly, we also observed target-signal sequential effects (Supplementary Figure 43), characterized by slower RTs (ER_prev_target<0_ = Inf.; ER_prev_target:target<0_ = 3.9×10^3^), and higher accuracy (ER_prev_target<0_ = 36.9) following trials with low target signal. These effects underscore that low discriminability of the cued dimension can elicit response caution independently from the congruency of non-cued dimensions.

This last experiment replicated the findings from the first two, indicating that the task induces multi-dimensional attentional control demands and that these are resolved in a dimension-specific way. They also illustrate that multiple mechanisms of control adaptation are simultaneously at play, with attentional gain primarily driven by distractor interference and response caution modulated by global response difficulty. Finally, the results suggested that adaptation effects in the MULTI are not driven by associative priming or other stimulus identity confounds.

## Discussion

In everyday life, we constantly need to switch attention between multiple competing sources of information. However, attentional control is typically investigated assuming a clear dichotomy between one task-relevant and one task-irrelevant dimension. Here, we designed a multidimensional task-set interference paradigm that allowed us to observe attentional control dynamics in multidimensional environments. We show that attentional control selectively adjusts the processing of each task-irrelevant dimension based on its own congruency history. Computational modeling demonstrated that, in order to capture this pattern of behavior, conflict must be monitored for each distractor separately, and detected at the level of stimulus representation.

This finding invites important revisions to established models of conflict adaptation. As mentioned before, classic models assume a single conflict detector at the response level, where information about individual stimulus features is no longer accessible. Our simulations show that this produces adaptation effects that transfer between distractor dimensions, unlike our findings. Interestingly, the domain-specific model proposed by Egner(Egner, 2008) assumes separate monitors for different conflict types along the information processing stream (e.g., semantic vs. response conflict), but at levels where individual dimension information is unavailable. This model explains conflict adaptation in tasks where distractors elicit different conflict types (e.g., a combined Stroop-Simon task(Braem et al., 2014)). In our task, however, all conflict happens exclusively at the stimulus level(Kornblum et al., 1990), yet we see no transfer of adaptation between dimensions. This flattens any proposed hierarchy of conflict monitors, allowing them to operate independently at the level of feature-based attention(Algom & Chajut, 2019). Finally, another class of models applies control to the connections between task representations and stimulus features(Blais et al., 2007; Verguts & Notebaert, 2008). For example, Verguts and Notebaert(Verguts & Notebaert, 2008) proposed the ‘adaptation by binding’ account by which conflict changes the connections between all task representations (cued and non-cued) and stimulus features, depending on the activity level of the connected units. Therefore, these models implement both target and distractor adaption. Because they measure conflict at the response level, they are agnostic to the original decomposition of the conflict. This means that they are unable to adapt individual stimulus dimensions based on dimension-specific information. Neither of these assumptions are met in our data. However, these models can simulate how people learn to attach attentional control settings to specific stimulus features(Bugg & Crump, 2012; Bugg & Hutchison, 2013; Jacoby et al., 2003). Our model does not account for such behavior. Future work may attempt to unify these perspectives, for example by assuming that Hebbian learning signals do not spread towards more local representations of stimulus features.

Unlike classic response interference tasks, the potency of the distracting dimensions in the MULTI does not derive from a natural asymmetry in the strength of tasks (such as reading vs. color naming in Stroop). Instead, the non-cued dimensions influence information processing because they are periodically task-relevant. This introduces a temporal dynamic in which the attentional gain of each dimension is boosted whenever it becomes task-relevant, and then slowly trails off (see Supplementary Figures 1 and 2). The recently proposed Learned Attention for Control model(Alexander, 2022) proposes that such dynamics can be captured by a reinforcement-learning algorithm that deploys attention to dimensions that appear to be useful for responding correctly. Similar to our account, this model assumes independent learning processes for each dimension. However, the model has no access to task cues, so it is completely reliant on trial-and-error learning. Therefore, it would likely struggle on tasks where relevant dimensions alternate often and unpredictably (such as the MULTI). Interestingly, Ritz and Shenhav(Ritz & Shenhav, 2023) have used attractor networks to model an analogous dynamic that resolves within a trial of an attentional control task. There, attention to distractor information is high at the onset of a trial and reduces to zero over time. Taking inspiration from this approach, attentional gains to the four dimensions of the MULTI could be independently regulated by replacing task units with corresponding attractor dynamics. This way, both increased salience of recently cued dimensions, and conflict-driven adaptations, could be modeled as perturbations of a dimension-specific attractor network across trials^2^.

A somewhat different modeling approach has successfully used action-outcome prediction errors(Alexander & Brown, 2011) and error likelihood(Brown, 2009; Brown & Braver, 2005) to explain brain signals traditionally associated with conflict detection. This line of work focuses on how control states track variables that can be explicitly learned, e.g., by associating incongruent stimuli with higher error probability or actions with their probable outcome. Moreover, independent probabilities can be simultaneously estimated for multiple response options, allowing these models to account for brain signals elicited in tasks where joint response activations are not necessarily incompatible(Brown, 2009). However, trial-and-error learning is generally slow. Thus, it is unlikely these models would effectively capture behavior in rapidly changing tasks such as the MULTI paradigm. Yet, these bottom-up solutions (including the Learned Attention for Control model), relying on simple associative learning, may complement the costly mechanisms of rule-based executive control(Kool et al., 2010). Our results may inform future models with the crucial notion that the learning signal cannot just be stimulus-general (response time, error likelihood, action-outcome), but must also be tied to the specific source of interference.

Interestingly, most models of attentional control resolve conflict by enhancing target information (but see Ritz & Shenhav, 2023), in clear disagreement with our data. This leads to a simple question: Why did our participants not engage in target enhancement? One possibility is that target enhancement is not a feasible physiological mechanism. A growing body of work suggests that a relative amplification of task-relevant information relies on inhibition of regions processing distractors(Jensen, 2023; Jensen & Mazaheri, 2010). Alternatively, it may be more effective to suppress distractors in our task, especially considering the high number of them. Moreover, more granular control adaptation leads to more optimal control settings, because each dimension can be individually tuned to its response history. For example, it allows for facilitation of a congruent dimension, even when the majority of dimensions is incongruent. This predicts that target enhancement is more likely to happen when distractors are mostly incongruent(Bugg & Crump, 2012; Jacoby et al., 2003) since their potential utility would be smaller. The MULTI is perfectly suited to test this prediction. It should be noted that our simulations were unable to capture congruency sequence effects when implementing target enhancement, underscoring it may not be a viable algorithmic solution. Perhaps other computational frameworks can demonstrate adaptation through target enhancement. However, our model closely followed the benchmark neural network developed by Cohen and colleagues(Cohen et al., 1990), suggesting that multiple innovations would be necessary for this to become feasible. At any rate, such adaptation would produce generalized transfer of control, which is inconsistent with what we observe, and often counterproductive. Perhaps, the pervasive need to tailor attention to individual dimensions in our environments inflates these implicit computational costs of increasing target sensitivity, which may be offset only when incentives are introduced(Ritz & Shenhav, 2023).

We should note that our results seem at odds with a small body of literature that finds general control adaptation in two-dimensional task designs. Transfer between distractor dimensions has been reported between two spatially-related dimensions(Kunde & Wühr, 2006), and cross-task transfer has been reported when both tasks shared identical response requirements(Notebaert & Verguts, 2008). These findings suggest that transfer of control may be possible if dimensions share some representational similarity(Braem et al., 2014). Such similarity could arise from perceptual factors(Kunde & Wühr, 2006), statistical associations between co-occurring features over time(Schapiro et al., 2012), or alignment of behavioral outcomes(Ileri-Tayar et al., 2022, 2024). In future studies, we will use the general framework of the MULTI to test this hypothesis.

Our results reveal that control adaptation in multidimensional environments involves post-error adjustments. Across our three experiments, we have replicated the typical observation that response times following incorrect trials are slower, suggesting increased response caution(Laming, 1979; Ritz et al., 2022). Moreover, our multidimensional paradigm allowed us to infer a similar adjustment in response caution proportional to the degree of incongruency on correct trials. This shift in behavior can be modeled through changes in the response threshold parameter of evidence accumulation models(Danielmeier et al., 2011; Fischer et al., 2018; Ratcliff et al., 2016; Ritz et al., 2022). However, our current focus was to demonstrate that dimension-specific adaptation only requires selective adjustments in attention to non-cued dimensions, which we implemented through reconfigurations in corresponding task units’ inputs. Nevertheless, the existence of post-conflict and post-error slowing suggest a parallel, dimension-agnostic, form of adaptation, that can be modeled as conflict-driven adjustments of the response threshold in the diffusion process. This process could draw on a distinct representation of general conflict (e.g., in the response layer), but it could also estimate the average dimension-specific level of conflict used for the reconfiguration of attentional control.

The distractor-specificity of conflict adaptation has implications for neuroscientific models of cognitive control. These models posit the anterior cingulate cortex (ACC) as the central hub for conflict detection(Botvinick et al., 2001; Holroyd & McClure, 2015). Neuronal populations in this region integrate current and previous interference, in line with behavioral CSE (Fu et al., 2022; Sheth et al., 2012), and are thought to transmit need-for-control signals to lateral prefrontal regions(Cavanagh & Frank, 2014; Shenhav et al., 2013; Verguts, 2017). Our findings challenge the notion that the ACC listens to compressed estimates of conflict. Instead, they suggest that the brain is equipped to monitor a wide variety of stimulus dimensions, and at least three of them in parallel. However, it seems implausible that the brain is set up with anatomically separated conflict monitors for each possible distractor one may encounter. Rather, we believe that our results point to a need to rethink the nature of neural representations of conflict.

One intriguing possibility is that the ACC relies on mixed selectivity in neuronal populations(Fusi et al., 2016; Rigotti et al., 2013) to encode multivariate conflict representations, rather than multiple univariate conflict signals. In the macaque prefrontal cortex (PFC), the presence of mixed selectivity neurons, which code for mixed representations of task variables, allows local integration of task rules and stimulus features, supporting context-dependent perceptual discrimination(Mante et al., 2013). The presence of mixed selectivity within the ACC has been confirmed recently by two different groups(Ebitz et al., 2020; Fu et al., 2022) who found that neurons coding for different types of conflict are intermixed. Thus, akin to the PFC, distributed neuronal populations of the ACC could simultaneously encode and integrate conflict arising from multiple novel dimensions(Heilbronner & Hayden, 2016). For example, the ACC could combine low dimensional subspaces, known as manifolds(Flesch et al., 2022), to flexibly allow for monitoring of incongruency from multiple sensory regions of the cortex. In such a framework, the role of the ACC, integrating task-relevant information from across the brain, would be consistent with Expected Value of Control theory(Shenhav et al., 2013). Conflict could then be resolved by suppressing interfering information(Danielmeier et al., 2011), represented within distal neural populations, possibly through oscillatory neural dynamics(Buschman & Miller, 2022; Jensen, 2023; Verguts, 2017).

Alternatively, the ACC could use these manifolds to encode stimulus information in local and task-specific representations(Flesch et al., 2022). Conflict would then arise from the readout of overlapping, collinear neural representations of incongruent dimensions. A computational advantage of these population-level representations is the possibility of overcoming conflict “locally” on a trial-by-trial basis. In fact, Ebitz and colleagues(Ebitz et al., 2020) found no abstract conflict coding axes in ACC populations. Instead, they suggested that conflict signals from ACC reflect the amplification of task-relevant sensorimotor information. In our multidimensional task, this predicts generalization of adaptation across dimensions, at odds with our data. However, complementary mechanisms may operate through suppression of single interfering stimulus features, their orthogonalization(Flesch et al., 2022; Ritz & Shenhav, 2024), or through an increase in dimension separability(Badre et al., 2021). In conclusion, hypotheses based on ‘local representations’ alleviate the need for a high degree of flexibility in connectivity between ACC and posterior regions, but it comes at the expense of an (inefficient) duplication of information from more sensory areas to medial frontal cortex. Our task and computational model may inspire a new research program to contrast these hypotheses.

## Conclusion

By expanding the dimensionality of traditional interference paradigms, we have gained valuable insights for our understanding of cognitive control. Our findings suggest that attentional control selectively adapts task-irrelevant stimulus dimensions based on their congruency history. Control adaptation occurs at the level of feature-based attention and involves multiple independent conflict-control loops. These results challenge previous models of control adaptation and highlight the need for alternative neurocomputational models of conflict representation. We hope that these findings can serve as inspiration for future investigations, leveraging increased complexity while maintaining experimental rigor.

## Method

### Participants

For Experiment 1, we recruited 151 participants from the student population of Washington University in St Louis. The experiment was administered online. To take part in the study participants were screened for color blindness, and their participation was compensated with curriculum credits. The experimental procedure was approved by the ethical committee (Institutional Review Board of Washington University in St Louis). Digital informed consent was obtained before the experiment, and participants were debriefed after the experiment. Inclusion criteria were set to 80% accuracy on each experimental task. Accordingly, data from 47 participants were excluded from further analysis, leading to a final sample size of N=104. Age: mean = 19.1, standard deviation = 1.3. Gender: male = 30, female = 73, non-binary = 1.

For Experiment 2, we recruited 172 participants from the same student population. None of them participated in Experiment 1. The procedure and exclusion criteria were the same as in Experiment 1. Accordingly, data from 68 participants were excluded from further analysis, leading to a final sample size of N=104. Age: mean = 19.5, standard deviation = 1.2. Gender: male = 40, female = 63, non-binary = 1.

For Experiment 3, we recruited 115 participants from the same student population. None of them participated in Experiment 1 or 2. The procedure was the same as in Experiment 1 and 2. The exclusion criteria was lowered from 80% to 70% accuracy on each experimental task, to account for the higher difficulty that emerged in this design. Accordingly, data from 12 participants were excluded from further analysis, leading to a final sample size of N=103. Age: mean = 19.9, standard deviation = 1.4. Gender: male = 26, female = 77, non-binary = 0.

### Experiment 1 - Multidimensional Task-set Interference Paradigm

In the MULTI, participants attend to a letter cue and choose between two stimuli that differ in four dimensions (color, shape, edge, dot motion). Specifically, for any dimension, participants had to find a target feature, and to select the corresponding side (pressing the ‘f’ or ‘j’ key for the left or right stimulus, respectively). At the beginning of the experiment, for each of the four dimensions, participants were instructed which feature was the target. For instance, a participant could receive instructions to ‘find the blue shape’ when the letter cue between the stimuli was C, ‘find the oval’ if it was S, ‘find the dashed-edged object’ if it was B, or ‘find the upward-moving dots’ if it was M. The target features were randomly assigned to each participant at the start and would remain the same for the entire experiment. Participants were trained on five blocks of practice trials to consolidate the dimension-target feature association, first on each cued dimension separately (until criterion of 5 correct trials in a row), and then on the four dimensions interleaved (10 correct trials in a row).

Participants were instructed to respond as quickly and as accurately as possible, and a response deadline of 1.5 s was imposed. An incorrect or late response was signaled by a red X at the center of the screen lasting 0.75 s. The inter trial interval (ITI) was set to 1 s, during which a fixation cross was shown at the center of the screen. In the practice phase, the fixation cross would turn green for 0.75 s after a correct response.

Cued dimensions could repeat or switch across trials. Repeat trials were preceded by one to four trials of the same dimension, while switch trials were preceded by trials of a different dimension(Liston et al., 2006). During the experiment, the sequence of cued dimensions was pseudo-randomized, so that the same dimension was cued a minimum of three and a maximum of five times in a row. All four dimensions were cued within four switch trials. Therefore, participants constantly cycled through the four tasks, organized in series of three to five repeat trials. Participants completed a total of 1600 trials split into two blocks. The average experiment duration was ∼60 min, including one self-paced break in the middle.

From this design, it follows that task-set interference is parametrically manipulated on a discrete scale ranging 0-3. On each trial, any of the three non-cued dimensions could be congruent if it primed the response side where the cued dimension’s target feature was located (i.e., the target features of the cued and non-cued dimensions were on the same side), or incongruent if it primed the opposite response side. Henceforth, congruency is defined as the number of non-cued dimensions that primed the correct response.

All versions of the task were developed with the “jsPsych” library(de Leeuw, 2015), using a customized version of the random-dot-kinematogram plugin(Rajananda et al., 2018). Task code is freely available at https://osf.io/j9fxc/.

### Experiment 2 – single stimulus task

In this experiment, participants completed a version of the MULTI used in Experiment 1, but with a few important modifications. First, a single stimulus was presented at the center of the screen. Across trials, the stimuli differed along the same four dimensions (color, shape, edge, and dot motion). The cue still indicated which dimension was relevant but was now presented at the center of the single stimulus object. In other words, only four features (one per dimension) were presented on any given trial. Second, for any cued dimension, participants were now instructed to report whether the target feature was present (by pressing the ‘f’ key) or absent (by pressing the ‘j’ key).

### Experiment 3 – continuous stimuli task

For this experiment, we again presented two stimuli flanking the task cue. Importantly, on each trial and for each dimension, the two objects varied in their appearance along a continuum. For the color dimension, the objects varied from more blue to more green (drawn from a 130° section of a circle in L*a*b color space, L= 70, center at [a,b] = [0, 0], radius = 55, and from 111° to 241°). For the shape dimension, the objects varied from more square-shaped to more circle-shaped, with the degree of ‘circleness’ being implemented as the size of the radius of rounded corners (0px to 50% of the square’s width). We ensured that both objects were drawn with the same surface size. For the edge dimension, the objects varied in the degree of thickness of their borders (1px to 10px). Finally, to accommodate a more perceptible continuous change on the fourth dimension, we manipulated the relative proportion of static black and white dots (100 dots per object; range 0% to 100% black dots). To ensure that within each dimension differences between objects were perceptible, we drew random features with a minimum distance of 20% of the entire range.

### Data analysis and inferential statistics

Switch trials were excluded from the current analyses unless specified. Additionally, we rejected outlier trials with responses faster than 200ms or slower than 5 standard deviations of individual participants’ reaction time distribution. To avoid contamination of conflict adaptation effects with different sources of control adaptation (e.g., post-error slowing), we discarded trials following incorrect responses and outlier trials, unless specified. Incorrect trials were further excluded for RT analyses.

Accuracy and reaction times were modeled with a Bayesian multilevel generalized linear approach implemented in R(R Core Team, 2017) with the “brms” package(Bürkner, 2017), which interfaces with the Stan language(Carpenter et al., 2017). The analysis pipeline included defining a probability model, computing the posterior distributions for each model’s parameter, and testing hypothesis based on the posterior probability distribution of the parameters and/or marginal means of interest. For each model, the estimation of the posterior distribution was obtained running four Markov Chain Monte Carlo (MCMC) simulations (i.e., chains). Each chain included at least 5000 iterations, of which 2000 were used for warmup. The frequency distribution of the resulting 12000 post-warmup samples were assumed as posterior probability of the model parameters. All models predicting reaction times assumed a shifted lognormal response distribution, with identity and log link functions for the distributional parameters. All models predicting accuracy assumed a Bernoulli distribution with a logit link function. Median and 95% credible intervals of the posterior parameters’ distributions are reported in the log scale (reaction times) or logit scale (accuracy), while estimated marginal means are reported in the response scales (with units ms and percentage, respectively). All models assumed weakly informative priors centered on zero for the parameters of interest, and on the raw mean for the intercept. For hypothesis testing we report evidence ratios (ER). For directional hypotheses, ER corresponded to the proportion of posterior probability samples supporting a hypothesis divided by the proportion supporting its alternative. For point hypothesis (two-sided), the ER was the Bayes Factor obtained with Savage-Dickey density ratio method, i.e. the posterior density at the point of interest divided by the prior density at that point.

### Neural network

We simulated behavior on our task using three different neural networks, modeled after influential neural networks of response interference(Botvinick et al., 2001; Cohen et al., 1990). Full code for these models will be provided on the Open Science Framework (https://osf.io/j9fxc/). All these networks shared the same basic structure.

### Basic structure

This structure consists of four independent pathways, one per dimension, each starting with a pair of input units that project to a set of hidden units in a hidden layer. Stimulus input reflected the location of the corresponding dimension’s target feature (left or right). Each input unit only projects to a single corresponding hidden unit with the weight of this connection set to 1.

Therefore, the representation of each individual dimension is maintained in the hidden layer, with eight units reflecting the location of each dimension’s target feature. Moreover, each pair of hidden units receives input from one corresponding ‘task’ unit (four units in total). Critically, the weight of each task unit’s connection to the corresponding pathway was set to 3, which was meant to partially offset a constant negative bias term of −4 for each hidden unit (for distractor-specific adaptation, best model performance was found setting these weights to 4 and −5, respectively). In the basic model, task units are activated when the corresponding dimension is cued. Total net input to each hidden unit was transformed to an activity value between 0 and 1 using the logistic activation function.

As described in detail by Cohen and colleagues(Cohen et al., 1990), activation of a task unit facilitates processing in the hidden units of the corresponding dimension. In short, differences in stimulus input activity will be larger for the cued dimension than for the non-cued dimensions, because the bias reduces sensitivity to input. Only for the cued dimension, the task units provide input that relieves this bias, allowing those units to reflect stimulus evidence with more sensitivity.

The hidden layer units then project to two response units (corresponding to a left and a right response respectively). Each hidden unit is connected to both response units, but with opposite signs. Specifically, hidden units that represent the presence of target features on the left project to the left response with a weight of 6, and to the right response with a weight of −6. In contrast, hidden units that represent the presence of target features on the right project to the right response with a weight of 6, and to the left response with a weight of −6. These values were chosen to ensure high variance in response units’ activations, which is necessary to estimate conflict values sensitive to congruency sequence effects. As before, net input for each response unit was transformed to an activity value between 0 and 1 using the logistic activation function.

### Diffusion process

In order to simulate response times and accuracy, we fed the difference in activity between the left and the right response unit (the ‘evidence’) to a diffusion process(Ratcliff et al., 2016). The model updates a decision variable *V* (starting at *V*_0_ = 0) by accumulating noisy information about the evidence over time. New evidence is accumulated until it exceeds either a negative threshold (*A* = −1) representing a left response, or a positive threshold (*A* = 1) representing a right response. Specifically, the model updates the decision variable as.

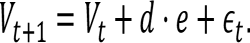

Here, *V*_*t*_ is the decision variable at time *t*, *d* is a drift rate parameter indicating the amount of evidence *e* that is sampled at each point in time, and ∈_*t*_ the amount of noise added to the decision variable. Critically, noise is distributed normally as ∈_*t*_∼*N*(0, σ), with σ the degree of noise. In all our simulations, we set *d* = 0.02 and σ = 0.1.

### Conflict monitoring

In order to simulate the parametric congruency sequence effects in our data, we took the core structure of this model, and implemented three different conflict monitoring mechanisms. Each of these mechanisms detects conflict (either between conflict response tendencies, or between feature representations), transforms this conflict into a measure of control, and then uses this to change the input to the task units on the following trial. Because congruency sequence effects were not present on switch trials, we did not implement this mechanism on those trials for any of the models.

Our first distractor-specific conflict monitoring model detects the degree of conflict for each non-cued dimension separately. Specifically, we define conflict as Hopfield energy within each dimension-specific pathway. Pathway energy is equal to the negative sum of the products of neuron states and their associated synaptic weights. Following Botvinick and colleagues(Botvinick et al., 2001), Hopfield energy(Hopfield, 1982) is defined as

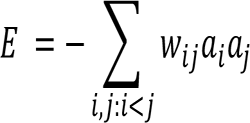

where *a* indicates unit activity and both subscripts are indexed over all units in the pathway.

Because of the excitatory and inhibitory configuration of the connections between intermediate and response units, both the sign and the magnitude of dimension-specific conflict are merely a function of the pathway state. It follows that dimension-specific conflict is positive if a non-cued dimension provides evidence inconsistent with the correct response, and negative if it provides evidence consistent with it.

Then, for each dimension, conflict is transformed into a measure of control using the following formula, adapted from Botvinick and colleagues(Botvinick et al., 2001):

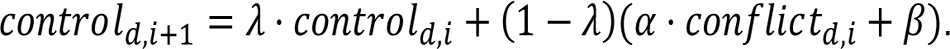

The parameters *d* and *i* index the dimension and trial, respectively. The parameter λ allows control to reflect a time-discounted measure of prior conflict. However, for simplicity and parsimony we set λ = 0 (but qualitatively similar effects were observed for non-zero values of this parameter). The parameters α and β are scaling parameters. Here, we set these parameters to α = 3.5 and β = 0. Unlike in the simulations described below, where β was used to set the sign of the control update, it was not necessary to assign β a non-zero value here, because the sign of each distractor-specific conflict value perfectly corresponded to whether the corresponding dimension was consistent or inconsistent with the correct response.

At the start of the next trial, each non-cued dimension’s control value was then subtracted from the (zero) input of that dimension. As described in the main text, this algorithm predicts distractor-specific adaptation effects, with the model becoming more susceptible to a non-cued dimension after a congruent trial on that dimension, and less susceptible after an incongruent trial.

The remaining two models measured a single form of conflict at the response level. Following Botvinick and colleagues(Botvinick et al., 2001), we assumed that the response units are interconnected by inhibitory weights, and computed Hopfield energy in the response layer. Here, more positive values indicate smaller evidence, less inhibition, and therefore increased conflict. For both, this measure of conflict is then transformed into control as

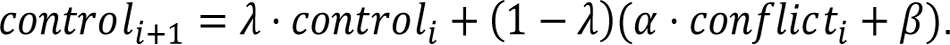

Not that this formula is almost identical to the one above, except that we do not index conflict and control by dimension, since this information is unavailable at the response level. For the simulations of the two remaining models, we set α = 3.5 and β = −0.56. Note that this value of β was not hand-tuned. Instead, β was set so that it would offset the conflict measure, yielding positive values for trials with more conflict, and negative values with less conflict. To do this, we first estimated the average amount of conflict in a model without adaptation (≈ −0.16), and we then multiplied this by α.

For the second model, implementing global distractor adaptation, the control measure was applied negatively to the task units of all non-cued dimension on the next trial. As described in the main text, this results in a parametric congruency sequence effect, but this adaptation is not distractor-specific. Instead, conflict arising from any distractor leads to an adaptation of susceptibility to all non-cued dimensions on the subsequent trial.

For the third model, implementing target adaptation, the control measure was applied to the task unit of the cued dimension. As described in the main text, this does not result in congruency sequence effects. Instead, the model becomes faster and more accurate if previous trials were more incongruent, but sensitivity to distractor information remains identical.

### Simulations

We simulated 1000 synthetic participants on each model. Each of these ran through the same number of trials as in our experiments (1600 trials). The process that generated the sequences of cued dimensions was identical to the experiments as well.

## Data availability

We provide access to all data and analysis scripts on the OSF (https://osf.io/j9fxc/).

## Code availability

We provide access to both task and simulation code on the OSF (https://osf.io/j9fxc/).

## Acknowledgements

We would like to thank Michael Freund, Roselyne Chauvin, Todd Braver, and Julie Bugg for helpful conversations. We also would like to thank the members of the Control and Decision Making Lab, and of the Connections with Control group for their advice and assistance, including Eric Gruber and Jack Dolgin for their contributions to early versions of the task. This work was supported by a Multi-University Research Initiative grant (ONR/DoD N00014-23-1-2792) to WK.

## Footnotes

1 Note that in the original model by Botvinick and colleagues(Botvinick et al., 2001), attention deployed to targets and distractors was bound together. Their model implemented control adaptation by first allocating attention to the target, and then setting complementary values to distractors. Here we decoupled these two mechanisms by allowing independent contributions to the response units from cued and non-cued dimensions’ pathways. Therefore, here a reduced processing of cued dimensions does not necessarily entail an increased processing on non-cued ones, and vice versa.

2 A similar approach can also successfully model the within-trial dynamics of interference that we observe in our experiments (see Supplementary Figures 35-40).

