## Supplementary Information for "Distractor-specific control adaptation in multidimensional environments"

### Supplementary results

### Experiment 1

Here, we report the results of Experiment 1 in more detail. Most importantly, we described Bayesian multilevel generalized linear models to explain within- and across-dimension adaptation effects at the single-trial level. For ease of reading, some of these results are also presented in the main text.

#### The number of congruent dimensions predicts task performance

We analyzed reaction times (RTs) and accuracy as a function of how many non-cued dimensions were congruent with the cued one using Bayesian linear regression. For both measures, the following linear model was used:

$$\sim 1 + \text{congruency} + (1 + \text{congruency} \mid \text{subject}) \qquad (1)$$

where *1* refers to the intercept, *congruency* is a numeric predictor, and the terms in parenthesis specify their random (subject level) effects.

We found that the number of congruent dimensions predicted both RT and accuracy. In particular, we observed a negative linear effect of congruency on RTs (Mdn = -0.04, 95% CI = [-0.04, -0.03], $ER_{congruency<0}$ = Inf), and a positive linear effect of congruency on accuracy (Mdn = 0.65, 95% CI = [0.61, 0.70], $ER_{congruency>0}$ = Inf). Participants were fastest when the stimulus was fully congruent, and became slower with each decrease in the number of congruent dimensions ($M_0$ = 694 ms, CI = [675, 713]; $M_1$ = 674 ms, CI = [656, 691]; $M_2$ = 655 ms, CI = [638, 671]; $M_3$ = 637 ms, CI = [621, 653]; $ER_{0>1}$ = 13.5, $ER_{1>2}$ = 14.8, $ER_{2>3}$ = 15.2).

Analogously, participants showed highest accuracy for fully congruent stimuli, which decreased with each increase in the number of congruent dimensions ($M_0$ = 91.4%, CI = [90, 92]; $M_1$ = 95.3%, CI = [95, 96]; $M_2$ = 97.5%, CI = [97, 98]; $M_3$ = 98.6%, CI = [98, 99]; $ER_{0>1}$ = Inf, $ER_{1>2}$ = Inf, $ER_{2>3}$ = Inf).

Further, we verified whether individual differences in the strength of the congruency effect on both measures were associated with each other. A negative Pearson's correlation between the random effects of congruency on either measure ($r$ = -0.25, $p$ = 0.01, 95% confidence intervals = [-0.42 - 0.06]) confirmed a subject-level association. This indicates that people who showed stronger susceptibility to congruency on RTs also showed stronger susceptibility on accuracy.

#### Interference effects persist over time

Congruency effects induced by a given non-cued dimension were the largest following a sequence of trials in which that dimension was the cued task, but they persisted over the following task switches. For this analysis, we grouped trials within blocks of task repetitions that followed either one of the four cued tasks. Because the trial list was composed of small lists in which each task was presented once, there could be one to six blocks following a task switch before a given task was cued again. To test the persistence of interference, we computed mean interference effects for each non-cued dimension as a function of the number of blocks that passed since it was the cued dimension (Supplementary Figure 1 & 2). Interference effects were largest on the first block after the switch, but remained reliable until block five (i.e., ~20 trials) for both RT (range BF: min = $8.8 \times 10^{1}$, max = $4.9 \times 10^{30}$) and accuracy (range BF: min = $1.0 \times 10^{4}$, max = $1.8 \times 10^{33}$).

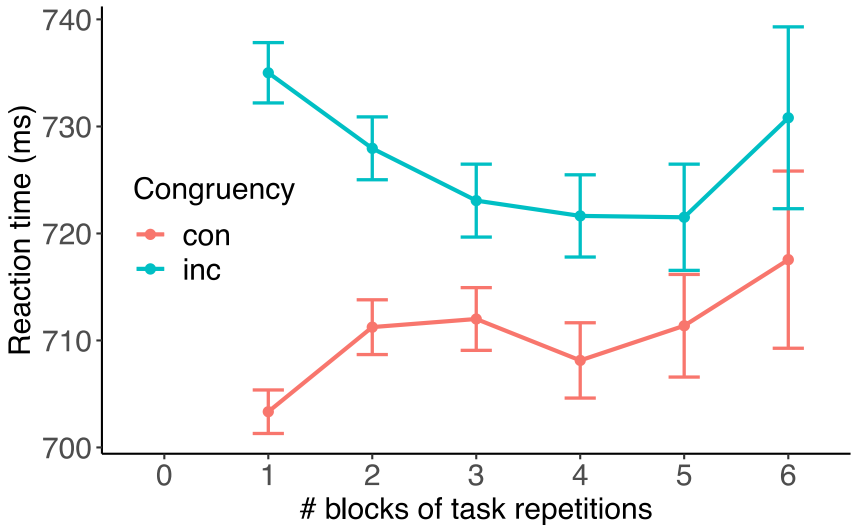

**Supplementary Figure 1.** In Experiment 1, the interference effect, measured in accuracy, of a non-cued dimension persists for 5 blocks after it was the cued dimension.

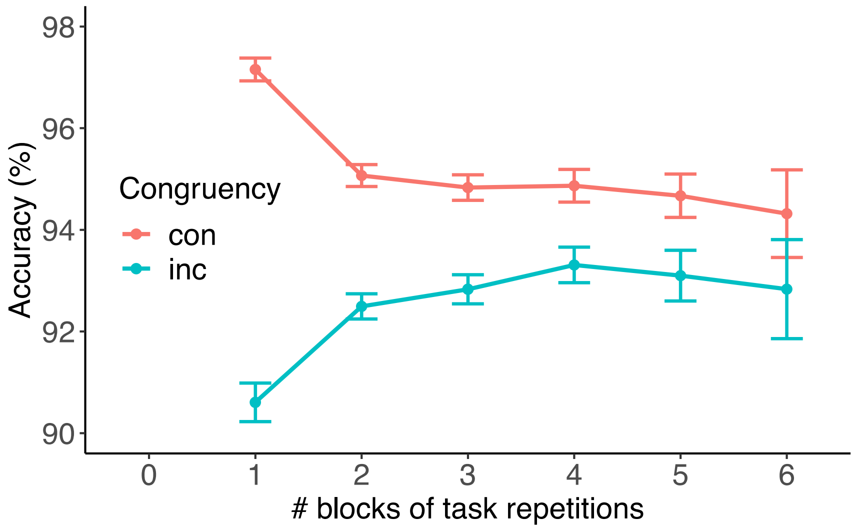

**Supplementary Figure 2.** In Experiment 1, the interference effect, measured in accuracy, of a non-cued dimension persists for 5 blocks after it was the cued dimension.

**Conflict adaptation is modulated by the degree of prior congruency**

Next, we measured susceptibility towards distraction as a function of congruence in the previous trial. Given that our multidimensional paradigm generates parametric congruency levels, we asked if the congruency sequence effect covaried with prior congruence. Therefore, we tested the interactive effects of the number of congruent non-cued dimensions across two subsequent trials. We predicted RT and accuracy with the following model:

~ congruency * previous_congruency + (1 + congruency * previous_congruency | subject) (2)

where *previous_congruency* is a numeric predictor specifying the congruency level in the preceding trial.

For RTs, we observed only anecdotal evidence in favor of a main effect of congruency (Mdn = -0.001, 95% CI = [-0.01, 0.00], $ER_{cong.<0}$ = 3.2), accounting for a very modest decrease of RTs with increasing congruency, conditional on previous congruency set to 0 ($M_{00}$ = 665 ms, CI = [649, 681]; $M_{03}$ = 662 ms, CI = [646, 679]). We observed strong evidence for a main effect of previous congruency (Mdn = 0.03, 95% CI = [0.03, 0.04], $ER_{prev\_cong.>0}$ = Inf), accounting for increasing RTs with increasing previous congruency, conditional on current congruency set to 0 ($M_{00}$ = 665 ms, CI = [649, 681]; $M_{30}$ = 724 ms, CI = [705, 743]). Importantly, we observed strong evidence in favor of an interaction effect between current and previous congruency (Mdn = -0.02, 95% CI = [-0.02, -0.02], $ER_{cong.:prev\_cong.<0}$ = Inf), accounting for larger effects of current congruency on RTs with increasing number of previous congruent dimensions.

As can be seen from the marginal means for the 16 levels of this interaction (Supplementary Figure 3), the effect of current congruency was strongest after fully congruent trials, and then parametrically declined as prior congruency decreased. Indeed, we observed only anecdotal evidence for an effect of congruency when the previous trial was fully incongruent (ER = 3.2, $M_{00}$ = 665 ms, $M_{03}$ = 662 ms). In contrast, there was very strong evidence for an effect of congruency after fully congruent trials (ER = Inf, $M_{30}$ = 724 ms, $M_{33}$ = 615 ms). In other words, compared to trials following fully incongruent trials, trials following fully congruent trials showed strong interference when current congruency was low (leading to longer RTs; $M_{30\text{-}00}$ = 59 ms, CI = [33, 83]) and strong facilitation when current congruency was high (leading to shorter RTs; $M_{33\text{-}03}$ = -46 ms, CI = [-69, -25]).

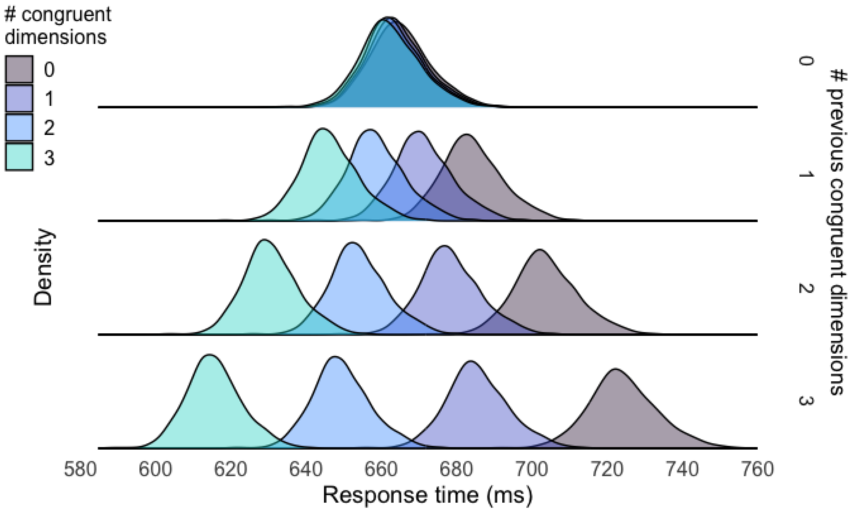

**Supplementary Figure 3.** Posterior distributions of the marginal means from a Bayesian linear model explaining response time as a function of both current (colors) and previous (rows) congruency in Experiment 1.

Analogous congruency sequence effects were observed for accuracy (Supplementary Figure 4). In contrast to the RT results, a strong main effect of congruency (Mdn = 0.28, 95% CI = [0.21, 0.35], $ER_{cong.>0}$ = Inf) indicated that accuracy increased with congruency even when holding previous congruency was 0 ($M_{00}$ = 96.2%, CI = [96, 97]; $M_{03}$ = 98.3%, CI = [98, 99]). In line with the RT results, we observed strong evidence for a main effect of previous congruency (Mdn = -0.53, 95% CI = [-0.58, -0.49], $ER_{prev\_cong.<0}$ = Inf), accounting for decreasing accuracy with increasing previous congruency, conditional on current congruency set to 0 ($M_{00}$ = 96.2%, CI = [0.96, 0.97]; $M_{30}$ = 83.8%, CI = [0.82, 0.86]). Importantly, there was strong evidence for an interaction between current and previous congruency (Mdn = 0.22, 95% CI = [0.18, 0.26], $ER_{cong.:prev\_cong.>0}$ = Inf), indicated by a larger effect of congruency with increasing numbers of previous congruent dimensions. Here, there was a strong effect of congruency even when the previous trial was fully incongruent (ER = Inf, $M_{00}$ = 96.2%, $M_{03}$ = 98.3%), but this effect was larger after fully congruent trials (ER = Inf, mean $M_{30}$ = 83.8%, $M_{33}$ = 98.8%). Similarly to RTs, we found interference and facilitation effects on accuracy ($M_{30-00}$ = -12.3%, CI = [-14, -11]; $M_{33-03}$ = 0.5%, CI = [0, 1]).

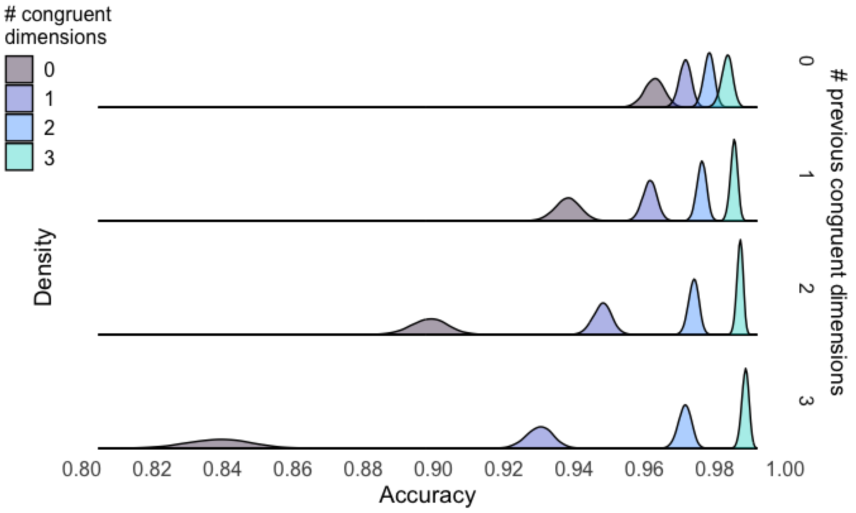

**Supplementary Figure 4.** Posterior distributions of the marginal means from a Bayesian linear model explaining accuracy as a function of both current (colors) and previous (rows) congruency in Experiment 1.

**Conflict adaptation weakens over time and resets after a task switch**

Thus far, we have only considered how multidimensional conflict is resolved within the same task and as a function of congruency in the previous trial (by excluding switch trials). However, the MULTI includes both task repeats and switches, which provide a natural way to study these temporal dynamics.

When we analyzed adaptation effects on switch trials alone (Supplementary Figure 5AB), using the model in Equation 2, we observed strong evidence against an interaction between current and previous congruency for both RT (Mdn = $-1.02\times10^{-3}$, 95% CI = [-0.00, 0.00], $ER_{cong.:prev\_cong.=0}$ = 1483) and accuracy (Mdn = $-5.81\times10^{-3}$, 95% CI = [-0.04, 0.06], $ER_{cong.:prev\_cong.=0}$ = 116). This suggests that control history effects do not survive a task switch, with control settings ‘resetting’ upon encountering new task demands.

The lack of conflict adaptation on switch trials is surprising. One possible reason is that the stronger interference on switch trials is likely driven by congruency of the non-cued dimension that was previously cued (note also the ‘recency’ effect in Supplementary Figures 1 and 2); however, because on any given trial the cued dimension is not assigned a congruency level (a cued dimension is always congruent with itself), its interference is not accounted in the analysis of switch-trial CSE (i.e., lack of previous congruency information). A second explanation for the lack of adaptation of switch-trial calls into account the presence of strong switch costs, which may overshadow the weaker CSE effects (e.g., RTs are overall higher on switch trials, introducing noise in the CSE estimation). Relatedly, the cognitive load induced by switching rule representations may tax the cognitive system and prevent utilizing control values from the previous trial. Last, we can speculate that control settings are both dimension-specific and task-specific. In other words, the cognitive system may track attentional gain values for each non-cued dimension, and each cued task (leading to 12 separate control values). By updating each value independently, model simulations replicate the observed reset of task settings on switch trials. To understand this, consider that on switch trials the sets of control values specific for the newly cued task are retrieved, and the resulting change in values (compared to the pre-switch trials) is unrelated to the change in congruencies.

Last, we tested whether congruency sequence effects persist over multiple trials. We modeled RTs and accuracy adapting Equation 2, where *previous_congruency* now specifies the congruency level in the $2^{nd}$-to-last or $3^{rd}$-to-last trial. First, we tested whether the number of congruent non-cued dimensions experienced two trials ago modulated the congruency effect on the current trials (Supplementary Figure 5CD). We observed strong evidence for such adaptation both in RT (Mdn = $-5.69\times10^{-3}$, 95% CI = [-0.01, 0.00], $ER_{cong.:prev\_2\_cong.<0}$ = Inf) and accuracy (Mdn = 0.08, 95% CI = [0.04, 0.12], $ER_{cong.:prev\_2\_cong.>0}$ = Inf). Next, we tested the CSE on the third-next trial (Supplementary Figure 5EF). Congruency experienced three trials ago reliably modulated congruency effects on the current trial for accuracy (Mdn = 0.06, 95% CI = [-0.02, 0.10], $ER_{cong.:prev\_3\_cong.>0}$ = 332), with only moderate evidence for RT (Mdn = $-1.33\times10^{-3}$, 95% CI = [0.00, 0.00], $ER_{cong.:prev\_3\_cong.<0}$ = 4.8).

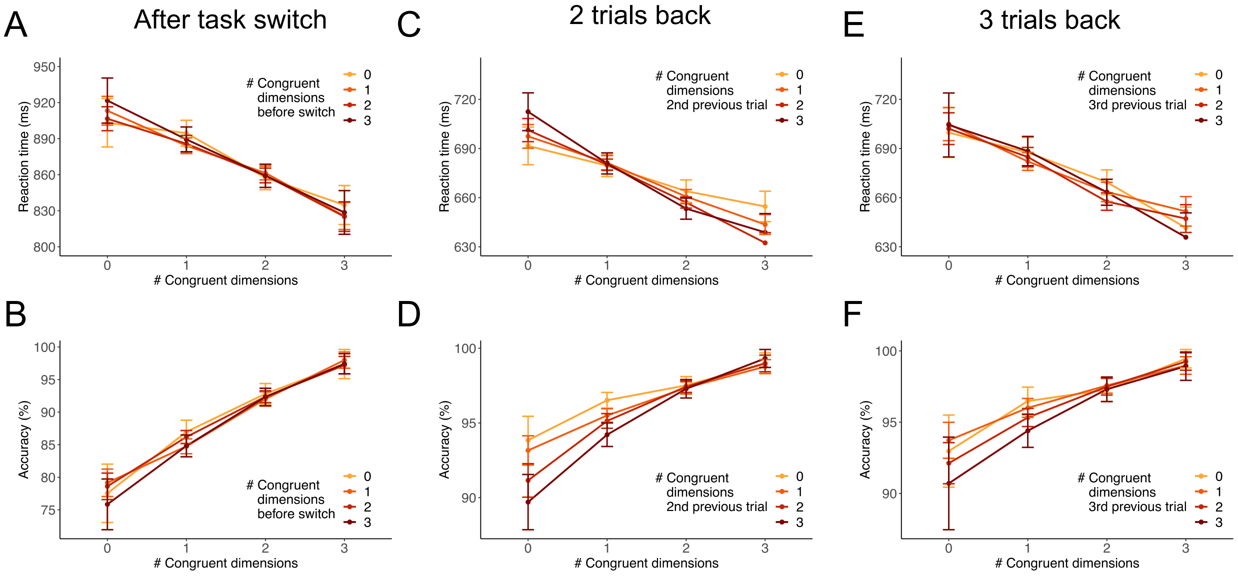

**Supplementary Figure 5.** Congruency sequence effects over time and after task switches in Experiment 1. We found that there was no conflict adaptation after task switches for (A) RT or (B) accuracy. However, we found that conflict from (CD) 2 trials back reliably induced adaptation for both measures. For conflict from 3 trials back we found moderate evidence of adaptation for (E) RT and strong evidence for accuracy.

Our model can capture these temporal dynamics in adaptation only when equipped with the distractor-specific adaptation mechanism. In particular, it can capture conflict adaptation from 2 or 3 trials back without any parameter changes. This appears to be an emergent property of sequential adaptations informed by Hopfield-based conflict, even when the mixing parameter $\lambda$ is set to 0 (i.e., the chain of dimension-specific conflict-control adaptations creates dependencies across multiple trials).

**Conflict adaptation is selectively driven within and not across dimensions.**

Here, we report analyses estimating trial-to-trial conflict adaptation within and across non-cued dimensions.

In the first analysis, reported in the main text, we computed *average* adaptation effects for each of the sixteen possible combinations of non-cued dimensions. These included congruent-congruent, congruent-incongruent, incongruent-congruent, and incongruent-incongruent (CC, CI, IC, II) sequences, for every combination of current and previous non-cued dimensions. Adaptation effects (for each participant) were obtained from the double difference of mean RTs [ (CI-CC) - (II-IC) ]. There, we showed that these adaptation effects were driven by sequences within dimension (on-diagonal, Figure 3AC), and not across dimension (off-diagonal).

Notably, the trial-average approach in the main text allowed us to isolate the adaptation effect for a given non-cued dimension, but it could not account for the cumulative effects of multiple non-cued dimensions acting in concert in any given trial, as suggested by the parametric CSE results. To confirm our results assuming this cumulative effect, we modeled adaptation effects at the single trial level. We first defined each trial's congruency sequence "fingerprint" by factorizing the presence of each sequence type (i.e., II, IC, CI, CC) for each combination of non-cued dimensions, and distinguishing between within-dimensions and across-dimensions. Next, we counted the instances of each type. This procedure resulted in the definition of 8 numeric predictors accounting for the presence and number of each sequence type, segregating within and across sequences. For any trial and sequence type, the number of "within" sequences could range from 0 to 3, while the number of "across" sequences could range from 0 to 6. Next, we predicted RTs with the following linear models:

$$rt \sim 1 + II\_within + IC\_within + CI\_within + (1 \mid subject) \quad (3)$$
$$rt \sim 1 + II\_across + IC\_across + CI\_across + (1 \mid subject) \quad (4)$$

where the intercept accounted for RTs assuming all the numeric predictors set to 0, and CC_within==3 for the within model or CC_across==6 for the across model.

In other words, the intercept value accounted for a trial in which all previous and all current dimensions were congruent. (Please note that 'all previous' and 'all current dimensions' refer to the combinations considered within each model). Therefore, the intercept value can be thought of as the easiest condition corresponding to the maximum number of CC sequences, and no instance of any other type. For example, including one instance of CI reduces the number of CC instances to 2 (for within models) or 5 (for across models). This is because there can be a maximum of 3 sequences within and 6 sequences across, and we use dummy-coded numeric predictors. Given this fixed number of possible sequences, to avoid multicollinearity between predictors, we implemented separate models to independently explain within and across CSE effects. To illustrate the cumulative effect of each numeric predictor, below we report their estimates when introducing a single instance of their sequence type.

The estimated marginal means from these two models are reported in Supplementary Figure 6. When considering currently incongruent non-cued dimensions (red distributions), we observed slower RTs in trials following a congruent vs. incongruent non-cued dimension of the *same* type (CI==1 vs. II==1, both implying CC==2; within effect: $M_{CI-II}$ = 18.6 ms, 95% CI = [15.3, 21.7], $ER_{CI-II>0}$ = Inf.), while there was no difference when comparing across dimensions sequences (across effect: $M_{CI-II}$ = 0.1 ms, 95% CI = [-4.52, 4.31], $ER_{CI-II>0}$ = 1.1). We also observed slower RTs for currently congruent dimensions (blue distributions) following the same dimension being incongruent in the previous trial (IC==1 with CC==2), as compared to the intercept value (CC==3; within effect: $M_{IC-Int}$ = 15.7 ms, 95% CI = [11.9, 19.5], $ER_{IC-Int>0}$ = Inf.). No evidence for an effect was observed when considering across dimensions sequences (across effect: $M_{IC-Int}$ = -0.7 ms, 95% CI = [-5.5, 3.9], $ER_{IC-Int<0}$ = 0.6), suggesting that a facilitation effect of CC was only observed in within-dimension sequences. In other words, we observed adaptation effects (including facilitation and interference) selectively when considering within-dimension sequences.

| Contrast | Evidence ratio | Mean difference | HDI low | HDI high |
|---|---|---|---|---|
| CI-II>0 - within | Inf | 18.60 | 15.34 | 21.75 |
| CI-II>0 - across | 1.06 | 0.08 | -4.52 | 4.31 |
| IC-Int>0 - within | Inf | 15.74 | 11.93 | 19.51 |
| IC-Int>0 - across | 0.61 | -0.73 | -5.52 | 3.93 |

**Supplementary Table 1.** RT contrasts within and across dimensions.

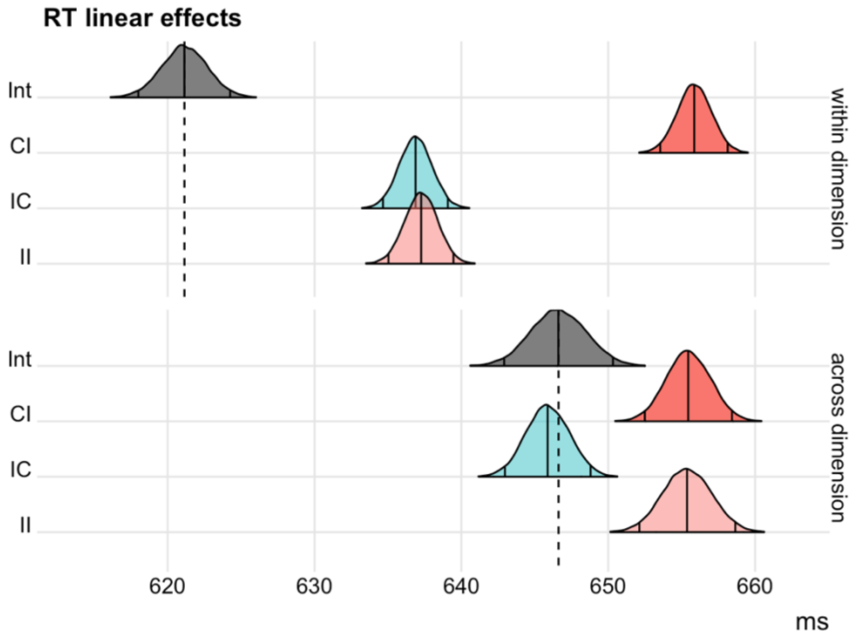

**Supplementary Figure 6.** Posterior distributions of the marginal means from two Bayesian linear models explaining RT adaptation effects in Experiment 1 as a function of either within-dimension conflict sequences (top row), or across-dimension conflict sequences (bottom row). The vertical solid lines represent the median and equal-tailed 95% credible intervals of the posterior samples. The intercept distributions (grey) correspond to the maximum number of CC sequences (within:

3; across: 6), and no instance of any other type. The other distributions represent the marginal means when introducing a single instance of the corresponding sequence type.

Analogous results were observed when predicting accuracy with the same approach (Supplementary Figure 7). Intriguingly, when turning our attention to across-dimensions sequences, we observed a reliable main effect of previous congruency, with lower accuracy after trials including additional congruent dimensions ($M_{CI-II}$ = -0.33 %, 95% CI = [-0.77, 0.08], $ER_{CI-II<0}$ = 14.9; $M_{Int-IC}$ = -0.49 %, 95% CI = [-0.84, -0.14], $ER_{Int-IC<0}$ = 386). In other words, when considering across-dimension sequences, the main effect of previous congruency suggests that accuracy increased if there were more incongruent dimensions in the previous trial.

| Contrast | Evidence ratio | Mean difference | HDI low | HDI high |
|---|---|---|---|---|
| CI-II<0 within | Inf | -1.26 | -1.51 | -1.01 |
| IC-Int<0 within | 1713.2 | -0.27 | -0.43 | -0.11 |
| CI-II<0 across | 14.8 | -0.33 | -0.77 | 0.08 |
| Int-IC<0 across | 386.1 | -0.49 | -0.84 | -0.14 |

**Supplementary Table 2.** Accuracy contrasts within and across dimensions.

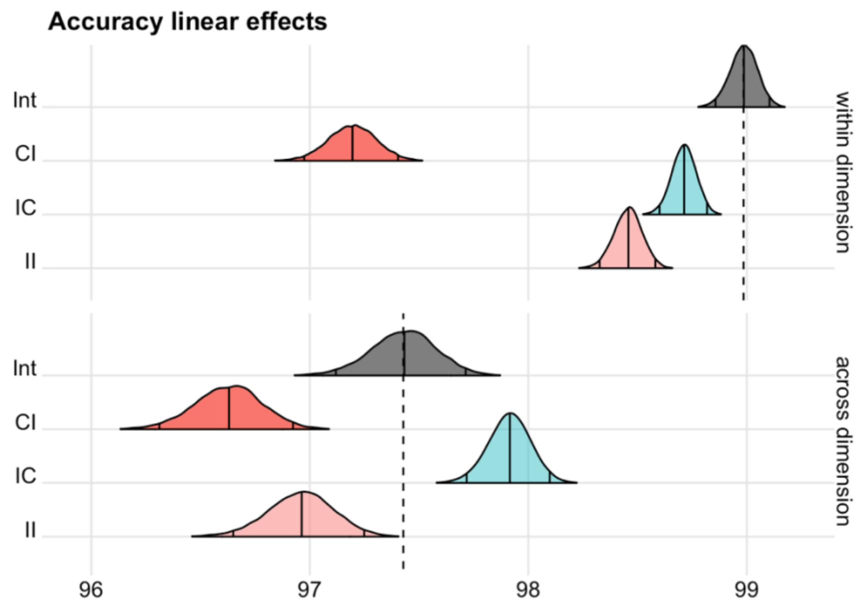

**Supplementary Figure 7.** Posterior distributions of the marginal means from two Bayesian linear models explaining accuracy adaptation effects in Experiment 1 as a function of either within-dimension conflict sequences (top row), or across-dimension conflict sequences (bottom row). The vertical solid lines represent the median and equal-tailed 95% credible intervals of the posterior samples. The intercept distributions (grey) correspond to the maximum number of CC sequences (within: 3; across: 6), and no instance of any other type. The other distributions represent the marginal means when introducing a single instance of the corresponding sequence type.

#### Dimensions' processing pathways are independent

To demonstrate that the stimulus processing pathways of the MULTI are independent, we sought to rule out the existence of interactive effects between non-cued dimensions' interference effects. First, we subset the data based on which dimension was cued. For each subset, we run a linear model predicting RTs including the main effects of congruency for each of the three non-cued dimensions, and the interactions between them:

RT(cued_dim) ~ 1+ congruency_1 * congruency_2 * congruency_3 + (1 + … | subject)

where 1 refers to the intercept, and congruency_i is a categorical predictor of congruency for a non-cued dimension. All main effects and interactions were also added as random effects.

Since the main effects of congruency of each non-cued dimension have been already reported, here we only tested their interaction effects against the point-null hypothesis. We found very strong evidence against interaction effects between every pair of non-cued dimensions, and across all of the four models run for each cued task (range $ER_{cong\_x:cong\_y=0}$ : min = 7.7, max = 320.6; median = 158.5). This result confirms that dimensions' interference effects do not interact. Rather, they show that RTs depend on the linear sum of each dimension's congruency effects.

#### Distributional analyses

The MULTI paradigm derives its strength from inducing parallel processing of multiple stimulus dimensions. However, each dimension exhibited a different interference profile (Figures 2BE, 4BE, 7BE). To investigate the temporal dynamics of dimensions' interference, here we report distributional analyses on accuracy and RT. By parsing congruence effects over different RT quantiles (van den Wildenberg et al., 2010), these analyses are informative about how conflict changes within a trial. These results of these analyses reveal intriguing heterogeneity in the way distractors' interfering effects develop within a trial. Moreover, they suggest that the time course of cumulative interference (and suppression thereof) results from the linear combination of the single dimensions' dynamics. Therefore, these results are compatible with the independent interference effects demonstrated above, and with the notion of independent and parallel dimensions' processing pathways.

#### Accuracy

We carried out distributional analyses of accuracy rates as a function of RT quantiles. First, we quantified the conditional accuracy function stratifying data by interfering (non-cued) dimension.

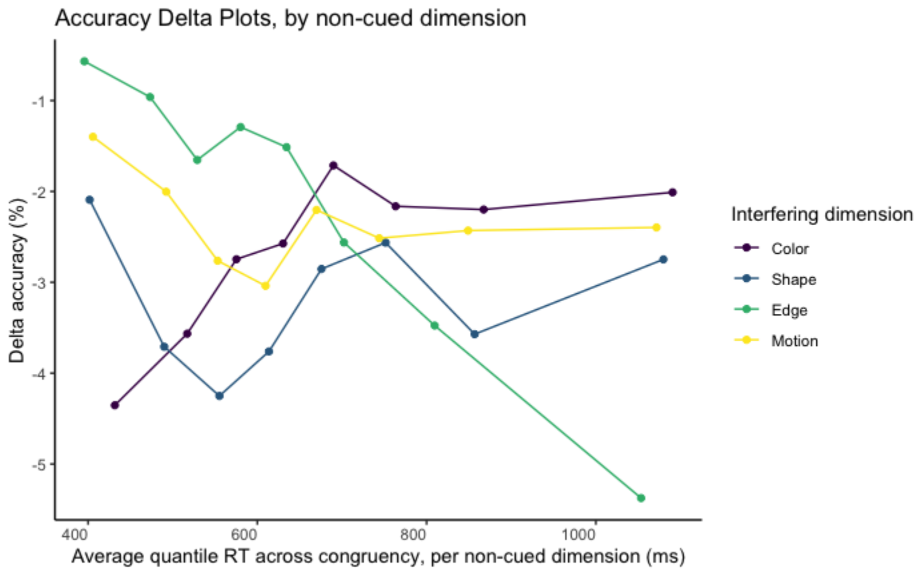

**Supplementary Figure 8.** Accuracy delta plots stratified by interfering dimension, obtained from contrasting incongruent vs. congruent trials, at each RT quantile. The RT quantile distributions were computed for each congruency level and interfering dimension. Data is plotted over the weighted RT mean across congruency levels, but separately for each interfering dimension. To note, each non-cued dimension exhibits a unique temporal profile. For example, the interfering dimension Color produced a strong initial attentional capture (high rate of fast errors for incongruent vs. congruent color), followed by a decline of such interference. The interference of the other dimensions exhibited different profiles leading to an increase or stabilization of interference.

The idiosyncratic nature of the temporal dynamic of dimension interference is reminiscent of the different magnitudes of their congruency effects. Next, we explore how these independent dynamics are combined across dimensions to produce the cumulative interference time course. To note, each dimension was cued (or non-cued) for an equal number of trials.

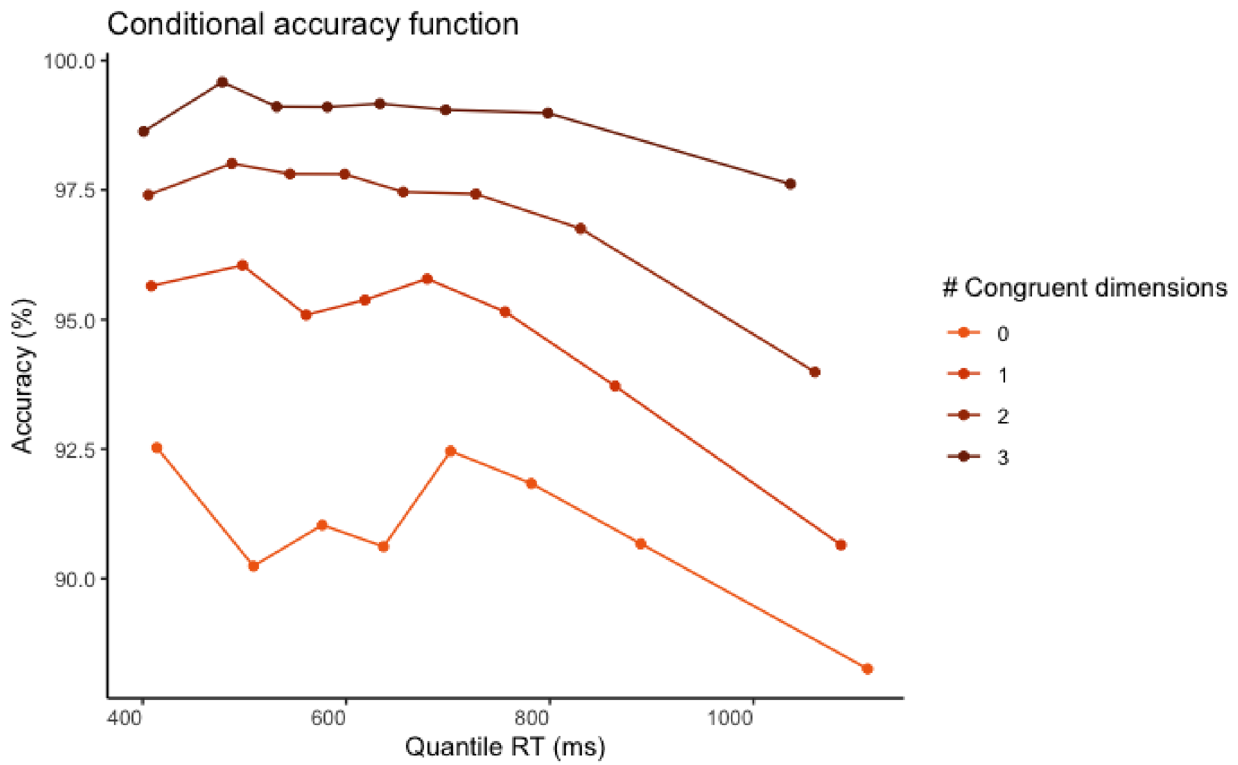

**Supplementary Figure 9.** Conditional accuracy function. Data is stratified by parametric congruency. The accuracy rate is computed separately for each RT quantile and congruency level.

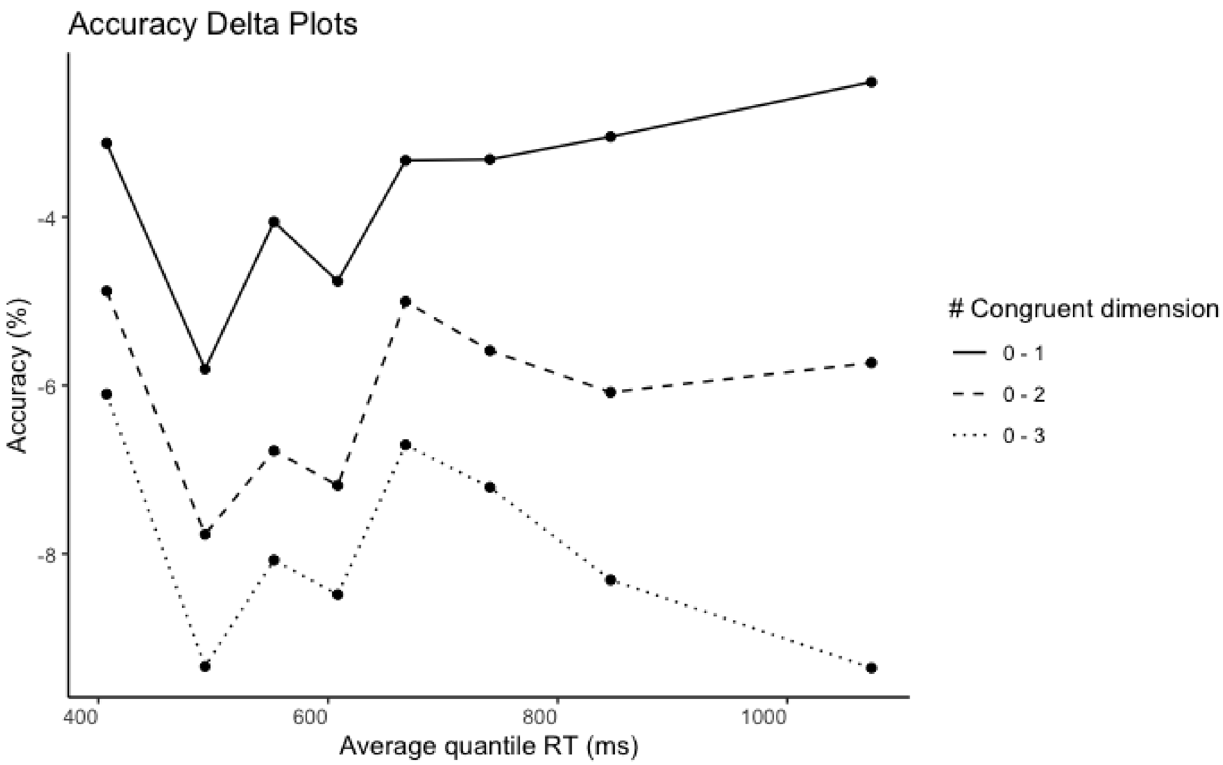

**Supplementary Figure 10.** Accuracy delta plots obtained contrasting congruency levels (zero vs. one, two, or three congruent dimensions) at each RT quantile.

The conditional accuracy function (the accuracy rate as a function of RT) suggests that most errors were either fast or slow responses. Importantly, only the intercept of the conditional accuracy function exhibited a linear increase as congruency levels rose. This is confirmed by overall flat accuracy delta plots when we contrasted congruency levels between each other. This pattern is compatible with a cumulative effect of interference (or facilitation) from independent non-cued dimensions.

**Reaction time**

Next, we carried out the same distributional analyses on RT.

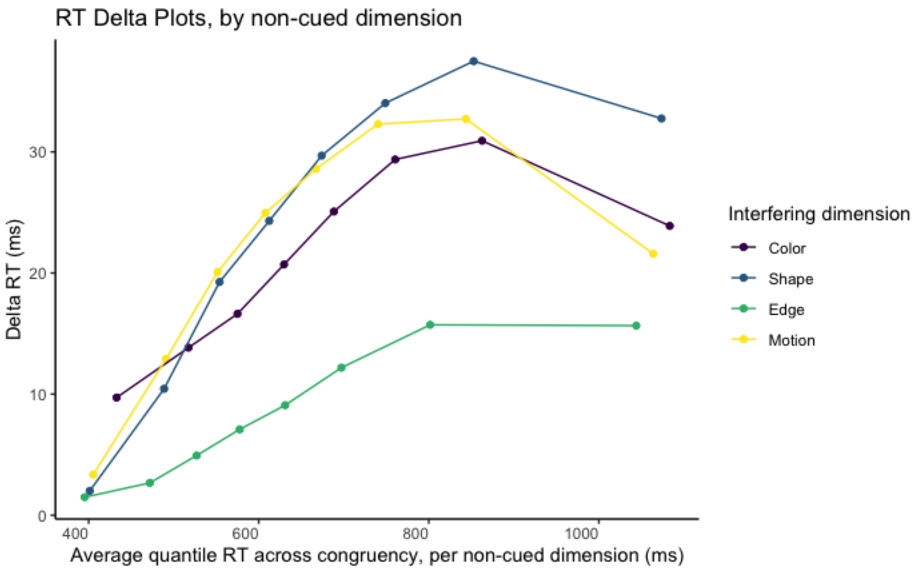

**Supplementary Figure 11.** Response time delta plots stratified by interfering dimension, obtained contrasting incongruent vs. congruent trials, at each RT quantile. RT quantile distribution were computed for each congruency level and interfering dimension. Data is plotted over the weighted RT mean across congruency level, but separately for each interfering dimension.

Similar to accuracy delta plots, each non-cued dimension exhibited a unique temporal profile. The slopes during early responses suggested different strengths and timing of interference. Next, we explore how these independent dynamics are combined across dimensions to produce the cumulative interference time course.

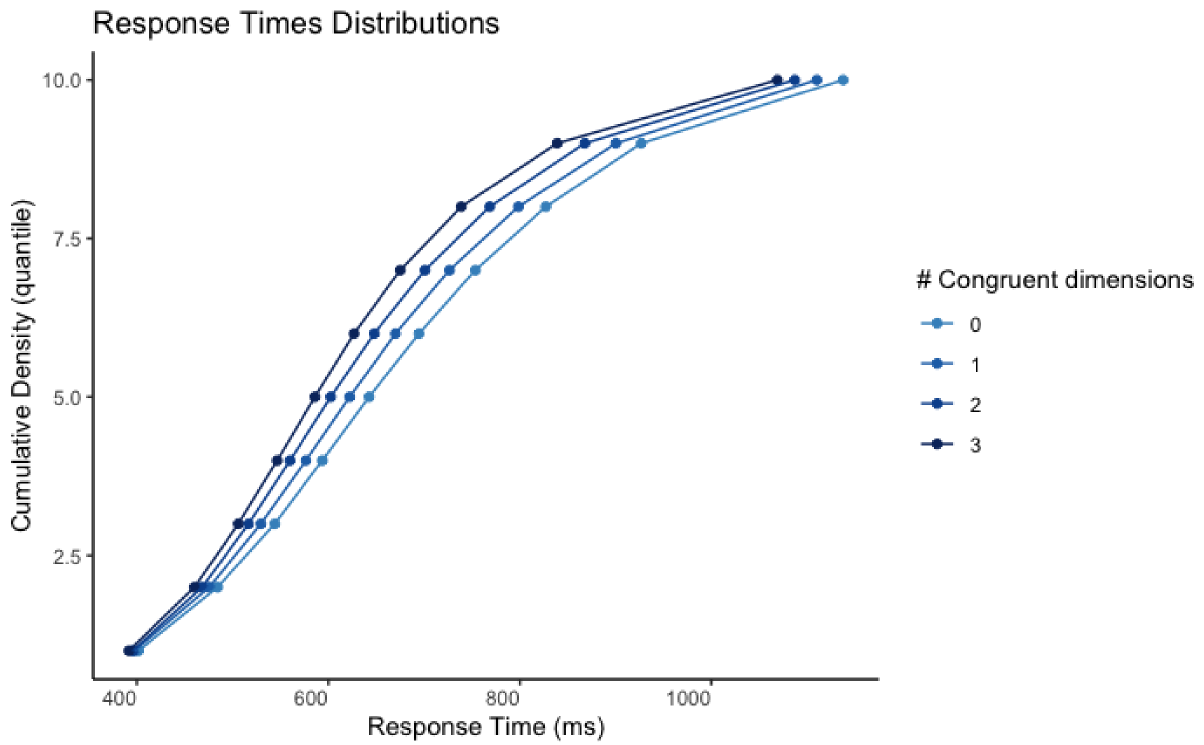

**Supplementary Figure 12.** Cumulative density distribution of response time. Data is stratified by parametric congruency.

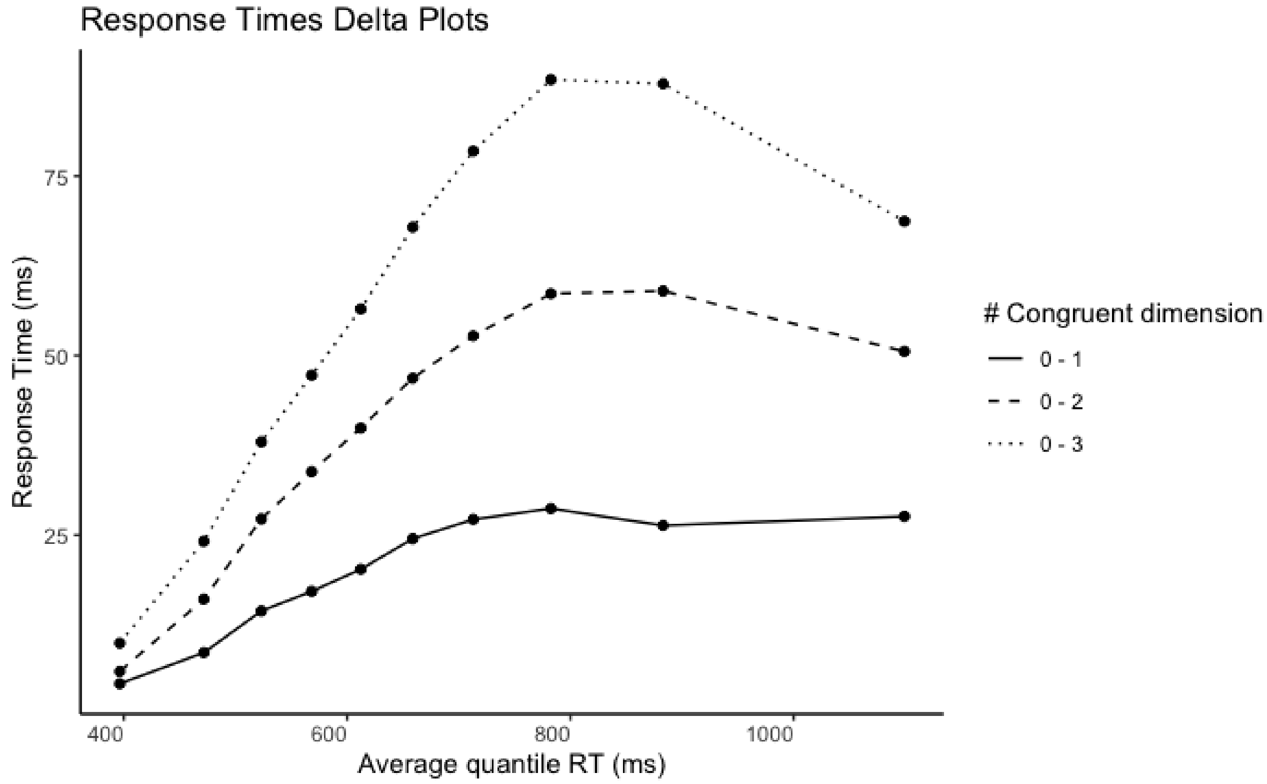

**Supplementary Figure 13.** Delta plots of interference effects obtained as RT difference between congruency levels (zero vs. one, two, or three congruent dimensions). Effects are shown over RT

quantiles, corresponding to the average RT of trials within each quantile and across congruency conditions, weighted by the corresponding trial count (note that the number of trials with congruency levels 1 and 2 is larger than with levels 0 and 3).

Congruency effects were the smallest for both early and late responses. The congruency effects increased and decreased within the RT range, with a peak time of around 800ms. Importantly, this peak increased linearly across congruency levels, again suggestive of a cumulative effect of interference (or facilitation) from independent non-cued dimensions. The temporal dynamic of the congruency effects suggests a distractor-based activation function (Mittelstadt et al., 2023), because the timing of non-cued dimensions' processing is temporally constrained. While in classic interference tasks (e.g., Eriksen flanker task, Simon task) Delta plots usually capture only a section of this function, our data show both the initial increase and the latter fading of interference (or facilitation) driven by non-cued dimensions. More generally, the temporal profile of the congruency effect in the first portion of the delta plots (earlier quantiles) is compatible with the results of a recent study (Mittelstadt et al., 2023) which manipulated the relevance of distracting information in Simon and Flanker tasks. The authors reported steeper and increasing slopes of delta plots when distractors were more relevant, reflecting a decreased and slowed suppression of potentially relevant stimulus dimensions. A similar mechanism can apply here, where an active suppression of non-cued dimensions may be slowed down and less efficient with more incongruent non-cued dimensions (or faster and stronger processing of non-cued dimensions with more congruent non-cued dimensions).

**Responses are slower and more accurate after an error.**

We excluded the trials following incorrect responses from all analyses to avoid contamination of conflict adaptation with other sources of adaptation, such as post-error slowing. Here we included these trials to highlight post-error slowing. We also observed increases in accuracy following errors. Together, these results suggest a post-error increase in response caution.

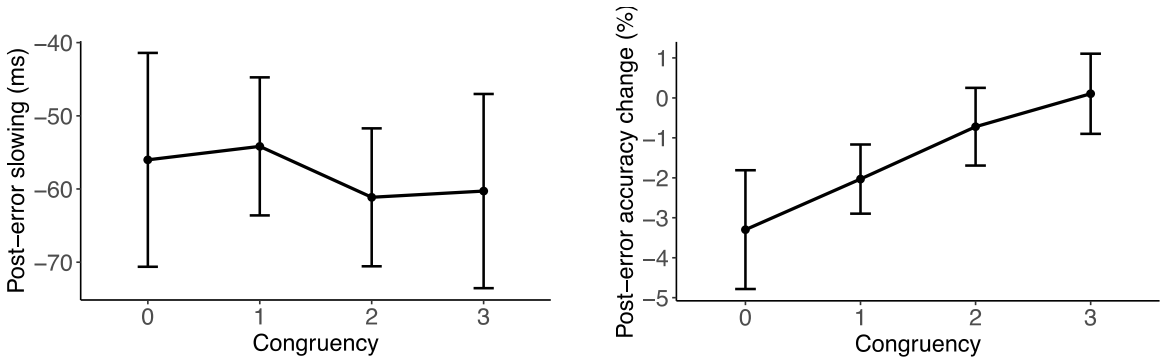

**Supplementary Figure 14.** RT (left) and accuracy (right) differences between trials following incorrect vs. correct responses. These effects suggest that participants become more cautious after committing an error, resulting in slower responses but higher accuracy. Error bars represent within-subject 95% confidence intervals.

### Experiment 2

Here, we report the results of Experiment 2 in more detail.

#### The number of congruent dimensions predicts task performance

As in Experiment 1, we found that the number of congruent dimensions predicted both RT and accuracy. In particular, we observed a negative linear effect of congruency on RTs (Mdn = -0.02, 95% CI = [-0.03, -0.02], $ER_{congruency<0}$ = Inf), and a positive linear effect of congruency on accuracy (Mdn = 0.44, 95% CI = [0.40, 0.48], $ER_{congruency>0}$ = Inf). Participants were fastest when the stimulus was fully congruent, and became slower with each decrease in the number of congruent dimensions ($M_0$ = 717 ms, CI = [699, 736]; $M_1$ = 705 ms, CI = [687, 723]; $M_2$ = 692 ms, CI = [674, 710]; $M_3$ = 680 ms, CI = [663, 698]; $ER_{0>1}$ = 5.1, $ER_{1>2}$ = 5.1, $ER_{2>3}$ = 5.1). Analogously, participants showed highest accuracy for fully congruent stimuli, which decreased with each increase in the number of congruent dimensions ($M_0$ = 91.9%, CI = [91, 93]; $M_1$ = 94.6%, CI = [94, 95]; $M_2$ = 96.5%, CI = [96, 97]; $M_3$ = 97.7%, CI = [97, 98]; $ER_{0>1}$ = Inf, $ER_{1>2}$ = Inf, $ER_{2>3}$ = Inf).

A negative Pearson's correlation between the random effects of congruency on RT and accuracy ($r$ = -0.42, $p$ < 0.001, 95% confidence intervals = [-0.57 -0.25]) confirmed a subject-level association between the two measures.

#### Each stimulus dimension produces interference

All non-cued dimensions produced interference for both RT (color: BF = $6.0\times10^{10}$, shape: BF = $1.7\times10^{12}$, edge: BF = $3.3\times10^{9}$, motion: $5.3\times10^{6}$) and accuracy (color: BF = $1.1\times10^{18}$, shape: BF = $2.5\times10^{21}$, edge: BF = $3.4\times10^{19}$, motion: BF = $1.0\times10^{13}$). In fact, we observed reliable interference from any non-cued dimension on any cued task (Figure 4BE) for accuracy (min BF = $1.8\times10^{3}$, max BF = $7.5\times10^{19}$). For RT, the interference of shape on motion was supported by moderate evidence (BF = 3.3), while there was strong evidence for any other effect (min BF = 10.1, max BF = $1.0\times10^{10}$).

On average between 3 and 4 non-cued dimensions induced a positive interference effect (RT: M = 3.15, SD = 0.82; accuracy: M = 3.55, SD = 0.71). If only one dimension provided interference per participant, we would expect 2.5 positive effects on average (1 from the single non-cued dimension that drove the effect, and 1.5 from 3 non-cued dimensions hovering around zero). Inconsistent with this hypothesis, the average number of positive interference effects was higher than this value for both RT ($BF_{(m>2.5\ /\ m<2.5)} = 2.6\times10^{25}$) and accuracy ($BF_{(m>2.5\ /\ m<2.5)} = 9.2\times10^{46}$).

#### Interference effects persist over time

To test the persistence of interference, we computed mean interference effects for each non-cued dimension as a function of the number of blocks that passed since it was the cued dimension (Supplementary Figure 15 & 16). Interference effects were largest on the first block after the switch but remained reliable until block four (i.e., ~16 trials) for both RT (range BF: min = 15.7, max = $6.7\times10^{22}$) and accuracy (range BF: min = $1.0\times10^{4}$, max = $7.1\times10^{28}$).

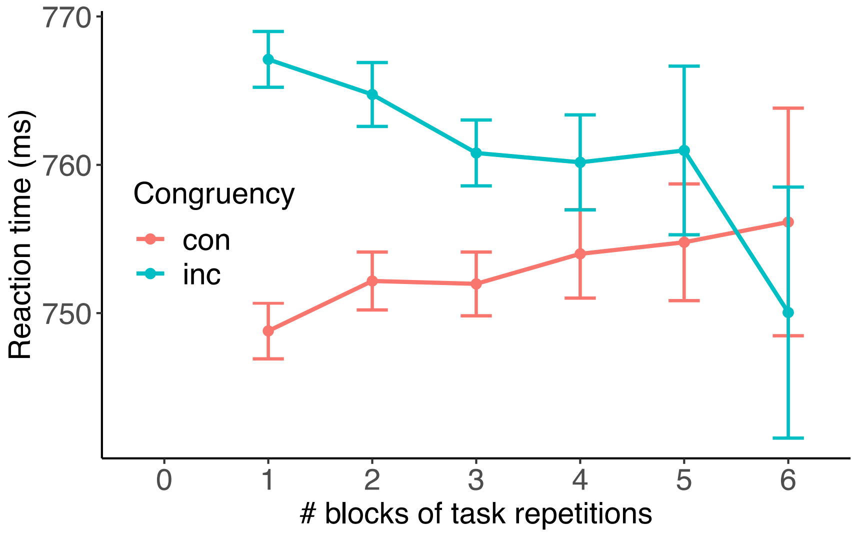

**Supplementary Figure 15.** In Experiment 2, the interference effect, measured in response time, of a non-cued dimension persists for four blocks after it was the cued dimension.

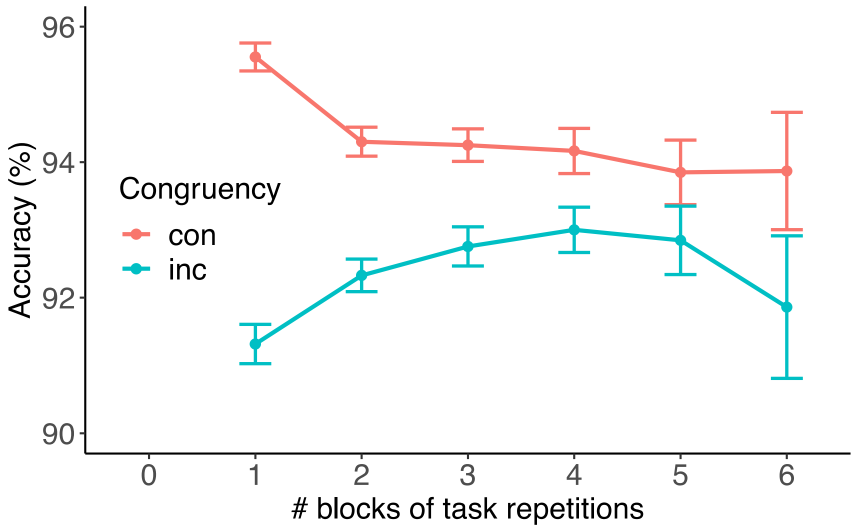

**Supplementary Figure 16.** In Experiment 2, the interference effect, measured in accuracy, of a non-cued dimension persists for four blocks after it was the cued dimension.

**Conflict adaptation is modulated by the degree of prior congruency**

The congruency sequence effects observed in Experiment 1 were replicated with the single-object version of the task.

For RTs, we observed moderate evidence in favor of a main effect of congruency (Mdn = 0.005, 95% CI = [0.00, 0.01], $ER_{cong.<0}$ = 0.03), but at difference with experiment one, it suggested a very modest *increase* of RTs with increasing congruency, conditional on previous congruency set to 0 ($M_{00}$ = 693 ms, CI = [675, 711]; $M_{03}$ = 701 ms, CI = [683, 720]). We observed strong evidence for a main effect of previous congruency (Mdn = 0.03, 95% CI = [0.03, 0.03], $ER_{prev\_cong.>0}$ = Inf), accounting for increasing RTs with increasing previous congruency, conditional on current congruency set to 0 ($M_{00}$ = 693 ms, CI = [675, 711]; $M_{30}$ = 742 ms, CI = [723, 763]). Importantly, we observed strong evidence in favor of an interaction effect between current and previous congruency (Mdn = -0.02, 95% CI = [-0.02, -0.02], $ER_{cong.:prev\_cong.<0}$ = Inf), accounting for larger effects of current congruency on RTs with increasing number of previous congruent dimensions.

As can be seen from the marginal means for the 16 levels of this interaction (Supplementary Figure 17), the effect of current congruency was strongest after fully congruent trials, and then parametrically declined as prior congruency decreased. Indeed, we observed evidence against a positive effect of congruency when the previous trial was fully incongruent (ER = 36.8, $M_{00}$ = 693 ms, $M_{03}$ = 701 ms). In contrast, there was very strong evidence for a positive effect of congruency after fully congruent trials, leading to faster responses (ER = Inf, mean $M_{30}$ = 742 ms, $M_{33}$ = 661 ms). In other words, compared to trials following fully incongruent trials, trials following fully congruent trials showed strong interference when current congruency was low (leading to longer RTs; $M_{30-00}$ = 49 ms, CI = [21, 75]) and strong facilitation when current congruency was high (leading to shorter RTs; $M_{33-03}$ = -40 ms, CI = [-64, -14]).

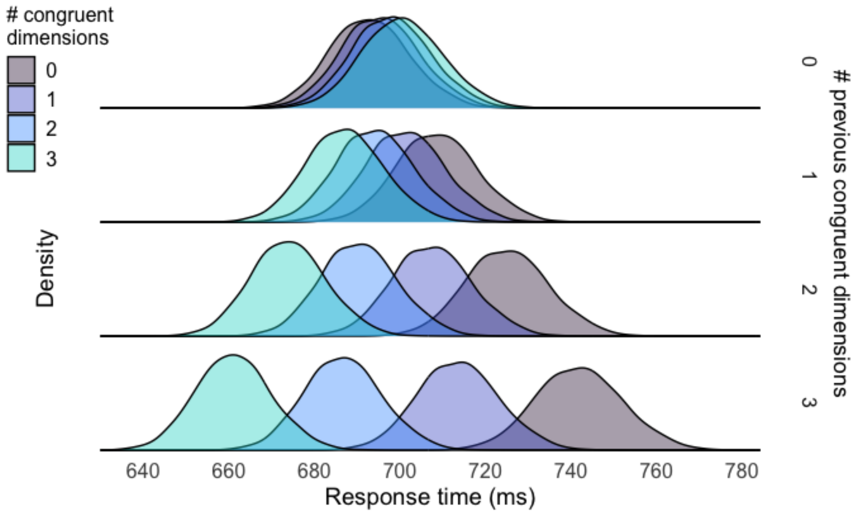

**Supplementary Figure 17.** Posterior distributions of the marginal means from a Bayesian linear model explaining response time as a function of both current (colors) and previous (rows) congruency in Experiment 2.

Analogous congruency sequence effects were observed for accuracy (Supplementary Figure 18). At difference with RTs, a moderate main effect of congruency (Mdn = 0.06, 95% CI = [0.01, 0.13], $ER_{cong.>0}$ = 17.4) indicated that accuracy increased with congruency even when holding previous congruency to level 0 ($M_{00}$ = 96.0%, CI = [95, 97]; $M_{03}$ = 96.7%, CI = [96, 97]). In line with the RTs results, we observed strong evidence for a main effect of previous congruency (Mdn = -0.46, 95% CI = [-0.51, -0.41], $ER_{prev\_cong.<0}$ = Inf), accounting for decreasing accuracy with increasing previous congruency, conditional on current congruency set to 0 ($M_{00}$ = 96.0%, CI = [95, 97]; $M_{30}$ = 85.9%, CI = [84, 87]). Importantly, there was strong evidence for an interaction between current and previous congruency (Mdn = 0.23, 95% CI = [0.20, 0.26], $ER_{cong.:prev\_cong.>0}$ = Inf) indicated by a larger effect of congruency with increasing numbers of previous congruent dimensions. Here, there was a moderate effect of congruency even when the previous trial was fully incongruent (ER = 13.9, $M_{00}$ = 96.0%, $M_{03}$ = 96.7%), but this effect was larger after fully congruent trials (ER = Inf, mean $M_{30}$ = 85.9%, $M_{33}$ = 98.3%). Similarly to RTs, we found interference and facilitation effects on accuracy ($M_{30-00}$ = -10.2%, CI = [-12, -8]; $M_{33-03}$ = 1.7%, CI = [1, 2]).

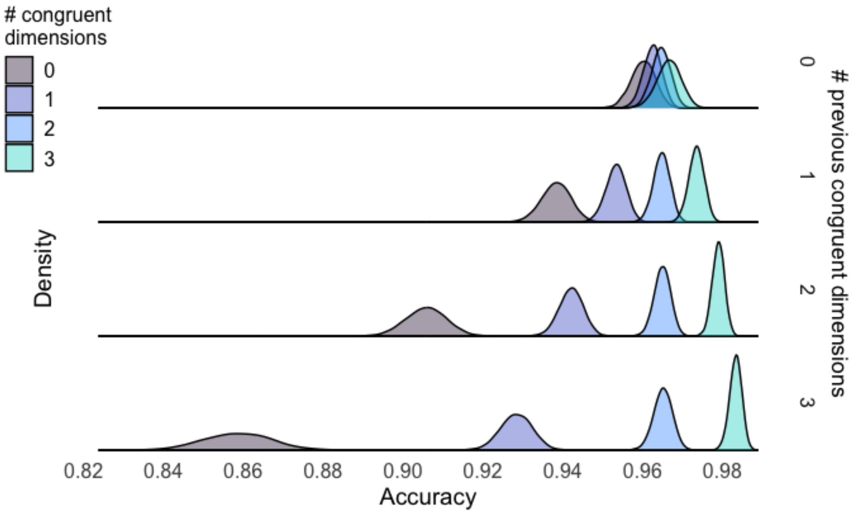

**Supplementary Figure 18.** Posterior distributions of the marginal means from a Bayesian linear model explaining accuracy as a function of both current (colors) and previous (rows) congruency in Experiment 2.

#### Conflict adaptation weakens over time and resets after a task switch

When we analyzed adaptation effects on switch trials alone (Figure 19AB), using the model in Equation 2, we observed strong evidence against an interaction between current and previous congruency for both RT (RTs: Mdn = $6.86\times10^{-4}$, 95% CI = [0.00, 0.00], $ER_{cong.:prev\_cong.=0}$ = 1870) and accuracy (Mdn = 0.01, 95% CI = [-0.04, 0.07], $ER_{cong.:prev\_cong.=0}$ = 98.3). This suggests that control history effects do not survive a task switch, with control settings 'resetting' upon encountering new task demands.

Last, we tested whether congruency sequence effects persist over multiple trials. We modeled RTs and accuracy adapting Equation 2, where *previous_congruency* now specifies the congruency level in the $2^{nd}$-to-last or $3^{rd}$-to-last trial. First, we tested whether the number of congruent non-cued dimensions experienced two trials ago modulated the congruency effect on the current trials (Figure 19CD). We observed strong evidence for such adaptation both in RT (Mdn = $-3.58\times10^{-3}$, 95% CI = [-0.01, 0.00], $ER_{cong.:prev\_2\_cong.<0}$ = 705) and accuracy (Mdn = 0.06, 95% CI = [0.03, 0.10], $ER_{cong.:prev\_2\_cong.>0}$ = 351). Next, we tested the CSE on the third-next trial (Figure 19EF). At difference with experiment one, congruency experienced three trials ago did not reliably modulated congruency effects on the current trial (RTs: Mdn = $1.67\times10^{-3}$, 95% CI = [0.00, 0.00], $ER_{cong.:prev\_3\_cong.<0}$ = 0.12; accuracy: Mdn = $2.10\times10^{-3}$, 95% CI = [-0.03, 0.04], $ER_{cong.:prev\_3\_cong.>0}$ = 1.19).

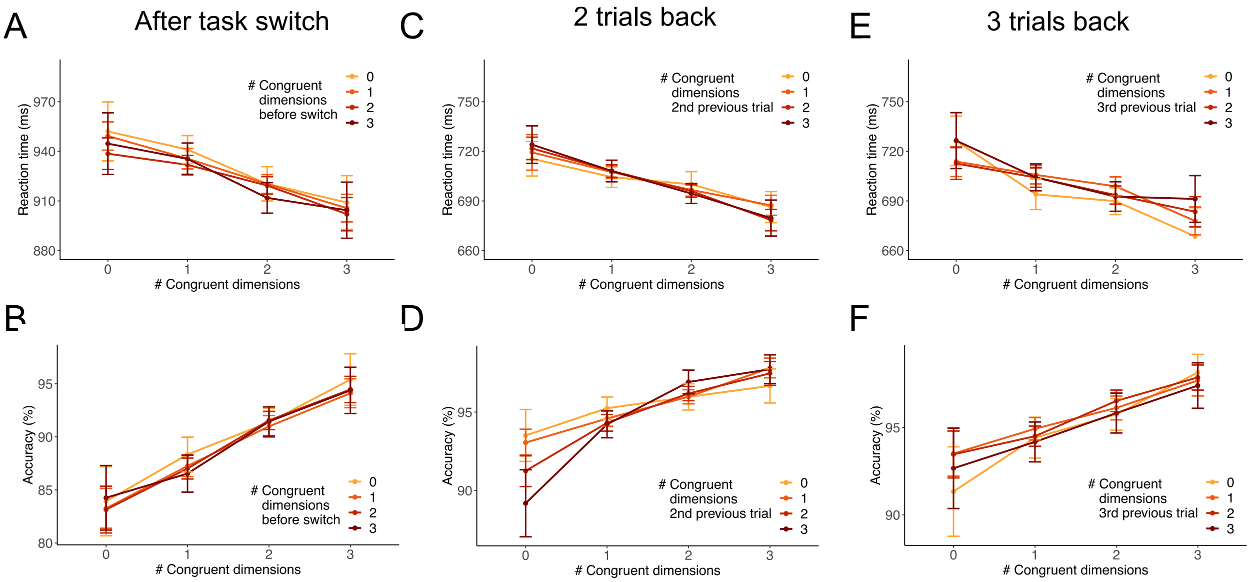

**Supplementary Figure 19.** Congruency sequence effects over time and after task switches in Experiment 2. We found that there was no conflict adaptation after task switches for (A) RT or (B) accuracy. However, we found that conflict from (CD) 2 trials back reliably induced adaptation for both measures. There was no evidence of adaptation based on conflict from 3 trials back (EF).

**Conflict adaptation is selectively driven by sequence effects within the same dimension.**

The main text reports the results of the trial-average approach, were we replicate that adaptation effects were driven by sequences within dimension (on-diagonal, Figure 4GI), and not across dimension (off-diagonal).

Next, we again modeled the adaptation effects at the single trial level, to test for cumulative effects of multiple non-cued dimensions acting in concert in any given trial. Congruency sequences within and across dimensions were modeled separately. The estimated marginal means from the two models are reported in Supplementary Figure 20. The cumulative effect of each numeric predictor is shown assuming a single CS instance (i.e., each numeric predictor is set to 1). When considering currently incongruent non-cued dimensions (red distributions), we observed slower RTs in trials following a congruent vs. incongruent non-cued dimension of the *same* type (CI==1 vs. II==1, both implying CC==2; within effect: $M_{CI-II}$ = 15.9 ms, 95% CI = [12.7, 19.1], $ER_{CI-II >0}$ = Inf.), while there was no difference when comparing across dimensions sequences (across effect: $M_{CI-II}$ = 0.3 ms, 95% CI = [-4.3, 4.7], $ER_{CI-II >0}$ = 1.2). We also observed slower RTs for currently congruent dimensions (blue distributions) following the *same* dimension being incongruent in the previous trial (IC==1 with CC==2), as compared to the intercept value (CC==3; within effect: $M_{IC-Int}$ = 13.5 ms, 95% CI = [9.7, 17.5], $ER_{IC-Int >0}$ = Inf.). No evidence for an effect was observed when considering across dimensions sequences (across effect: $M_{IC-Int}$ = -0.6 ms, 95% CI = [-5.4, 4.2], $ER_{IC-Int >0}$ = 0.6), suggesting that a facilitation effect of CC was only observed in within-dimension sequences. In other words, we observed adaptation effects (including facilitation and interference) selectively when considering within-dimension sequences.

| Contrast | Evidence ratio | Mean difference | HDI low | HDI high |
|---|---|---|---|---|
| CI-II>0 - within | Inf | 15.90 | 12.74 | 19.10 |
| CI-II>0 - across | 1.21 | 0.27 | -4.26 | 4.71 |
| IC-Int>0 - within | Inf | 13.47 | 9.70 | 17.49 |
| IC-Int>0 - across | 0.66 | -0.63 | -5.42 | 4.24 |

**Supplementary Table 3.** RT contrasts within and across dimensions.

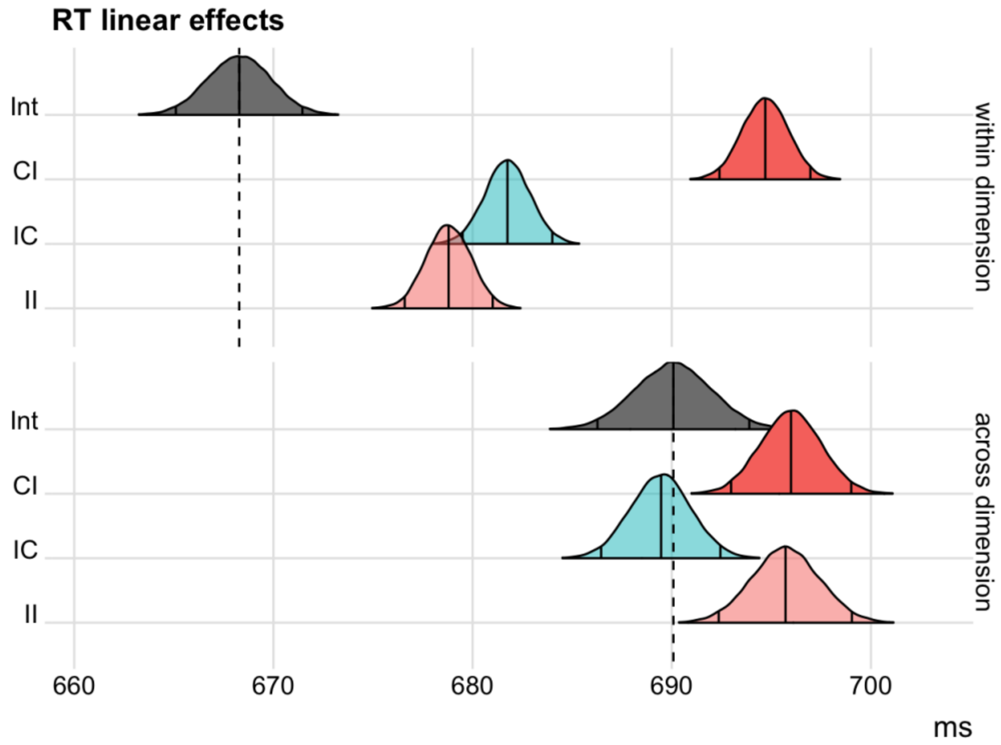

**Supplementary Figure 20.** Posterior distributions of the marginal means from two Bayesian linear models explaining RT adaptation effects in Experiment 2 as a function of either within-dimension conflict sequences (top row), or across-dimension conflict sequences (bottom row). The vertical solid lines represent the median and equal-tailed 95% credible intervals of the posterior samples. The intercept distributions (grey) correspond to the maximum number of CC sequences (within: 3; across: 6), and no instance of any other type. The other distributions represent the marginal means when introducing a single instance of the corresponding sequence type.

Analogous results were observed when predicting accuracy with the same approach (Supplementary Figure 21). Intriguingly, when turning our attention to across-dimension sequences, we observed a reliable main effect of previous congruency, with lower accuracy after trials including additional congruent dimensions ($M_{CI-II}$ = -0.30 %, 95% CI = [-0.8, 1.7], $ER_{CI-II<0}$ = 8.2; $M_{Int-IC}$ = -0.30 %, 95% CI = [-0.7, 0.1], $ER_{Int-IC<0}$ = 10.4). In other words, when considering across-dimension sequences, the main effect of previous congruency suggests that accuracy increased if there were more incongruent dimensions in the previous trial.

| Contrast | Evidence ratio | Mean difference | HDI low | HDI high |
|---|---|---|---|---|
| CI-II<0 within | Inf | -1.39 | -1.70 | -1.10 |
| IC-Int<0 within | Inf | -0.52 | -0.76 | -0.28 |
| CI-II<0 across | 8.27 | -0.30 | -0.78 | 0.17 |
| Int-IC<0 across | 10.45 | -0.30 | -0.73 | 0.14 |

**Supplementary Table 4.** Accuracy contrasts within and across dimensions.

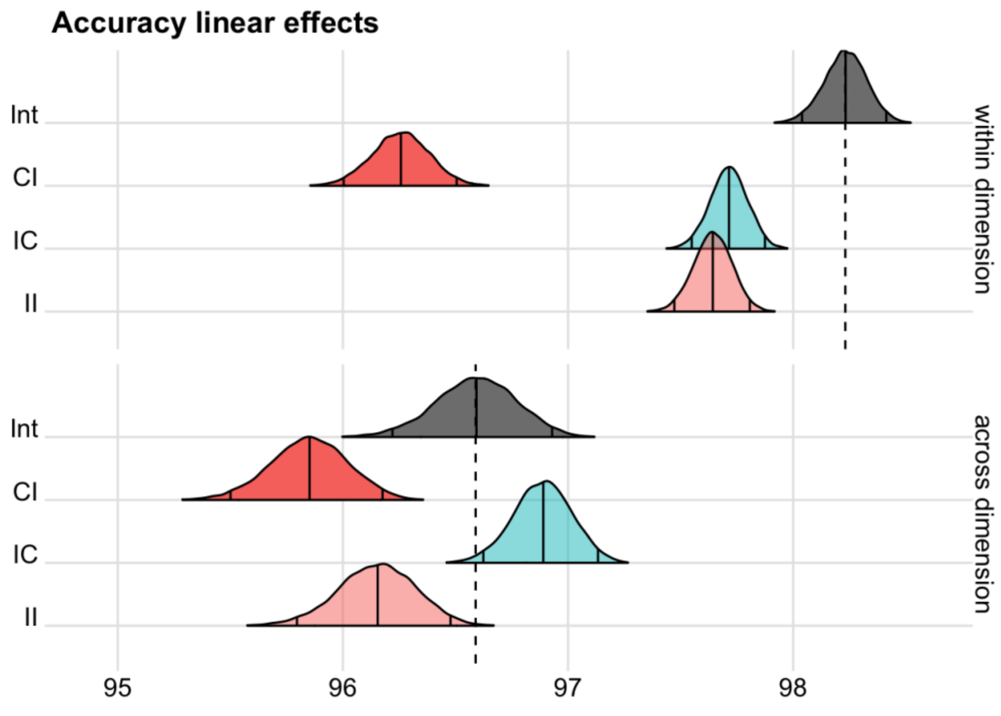

**Supplementary Figure 21.** Posterior distributions of the marginal means from two Bayesian linear models explaining accuracy adaptation effects in Experiment 2 as a function of either within-dimension conflict sequences (top row), or across-dimension conflict sequences (bottom row). The vertical solid lines represent the median and equal-tailed 95% credible intervals of the posterior samples. The intercept distributions (grey) correspond to the maximum number of CC sequences (within: 3; across: 6), and no instance of any other type. The other distributions represent the marginal means when introducing a single instance of the corresponding sequence type.

To rule out that each participant only applied within-dimension adaptation for one dimension, we computed participant's average number of positive within-dimension effects for both RT (RT: M = 3.31, SD = 0.82) and accuracy: (M = 3.34, SD = 0.74), and found strong evidence that these numbers were larger than the 2.5 positive adaptation effects one would expect under the alternative explanation (RT: $BF_{(m>2.5\ /\ m<2.5)} = 1.0 \times 10^{29}$, Accuracy: $BF_{(m>2.5\ /\ m<2.5)} = 1.3 \times 10^{38}$).

**Dimensions' processing pathways are independent**

To demonstrate that the stimulus processing pathways of the MULTI are independent, we sought to rule out the existence of interactive effects between non-cued dimensions' interference effects. For each cued task, we run a linear model predicting RTs including the main effects of congruency for each of the three non-cued dimension, and the interactions between them.

We found very strong evidence against interaction effects between every pair of non-cued dimensions, and across all of the four models run for each cued task (range $ER_{cong\_x:cong\_y=0}$ : min = 27.1, max = 285.3; median = 175.5). This result confirms that dimensions' interference effects do not interact. Rather, they show that RTs depend on the linear sum of each dimension's congruency effects.

### Distributional analyses

To investigate the temporal dynamics of dimensions' interference, here we report distributional analyses on accuracy and RT. In line with Experiment 1, the results of these analyses suggest that the time course of cumulative interference (and suppression thereof) results from the linear combination of the single dimensions' dynamics.

### Accuracy

We carried out distributional analyses of accuracy rates as a function of RT quantiles. First, we quantified the conditional accuracy function stratifying data by interfering (non-cued) dimension.

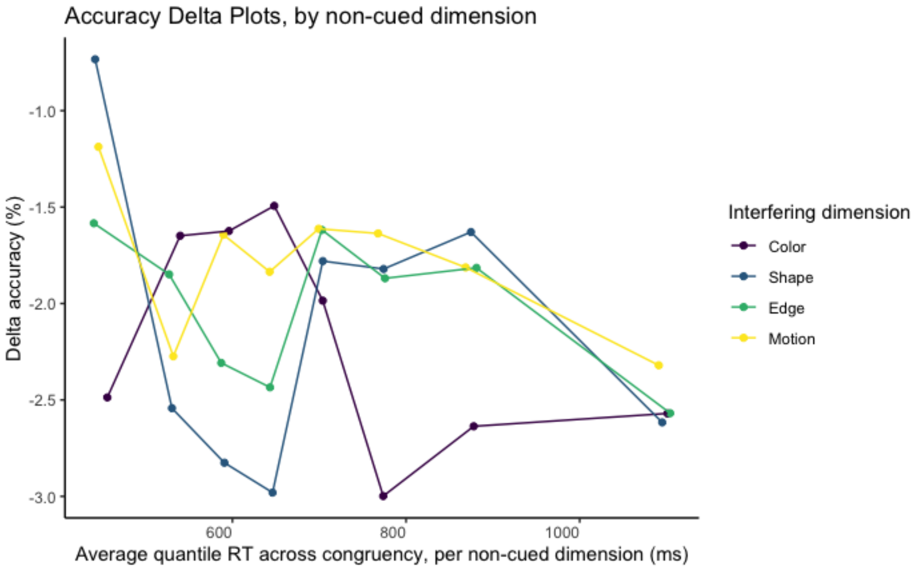

**Supplementary Figure 22.** Accuracy delta plots stratified by interfering dimension, obtained from contrasting incongruent vs. congruent trials, at each RT quantile. The RT quantile distributions were computed for each congruency level and interfering dimension. Data is plotted over the weighted RT mean across congruency levels, but separately for each interfering dimension. To note, each non-cued dimension exhibits a unique temporal profile. For example, the interfering dimension Color produced a strong initial attentional capture (high rate of fast errors for incongruent vs. congruent color). The idiosyncratic nature of the temporal dynamic of dimension interference is reminiscent of the different magnitudes of their congruency effects.

Next, we explored how these independent dynamics are combined across dimensions to produce the cumulative interference time course. To note, each dimension was cued (or non-cued) for an equal number of trials.

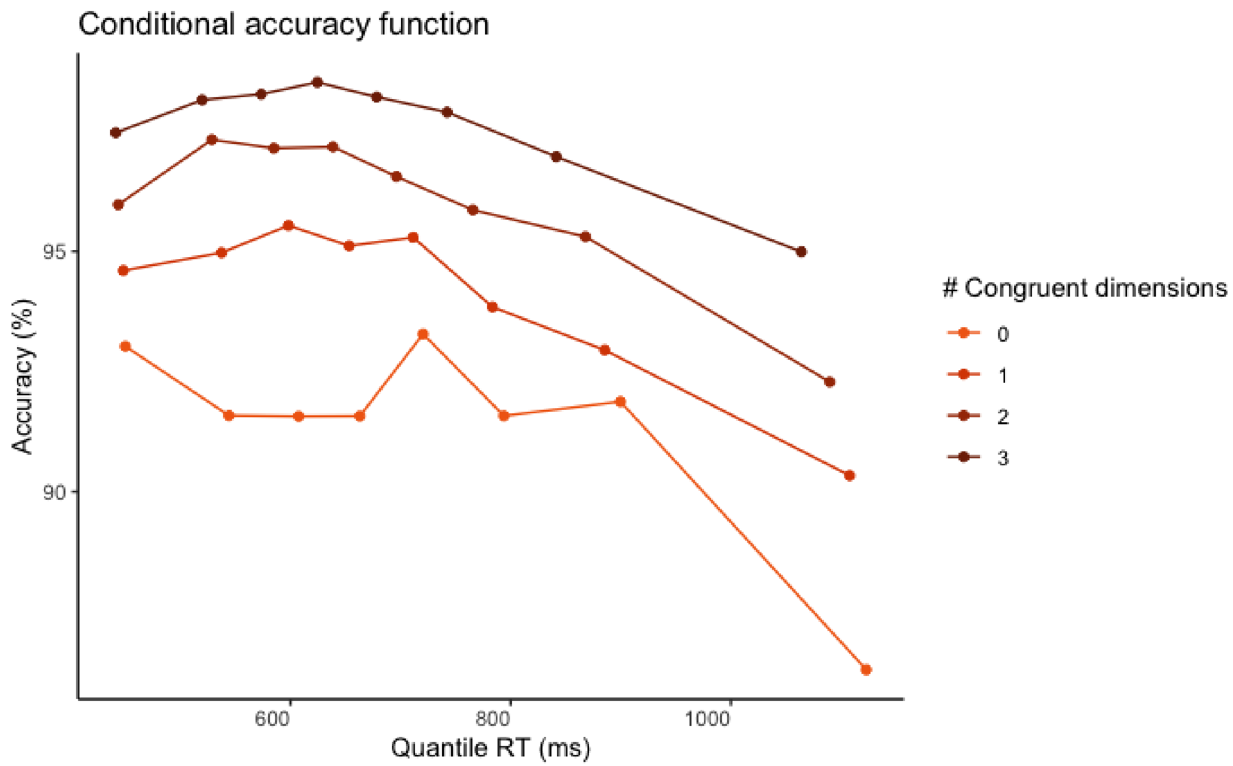

**Supplementary Figure 23.** Conditional accuracy function. Data is stratified by parametric congruency. The accuracy rate is computed separately for each RT quantile and congruency level.

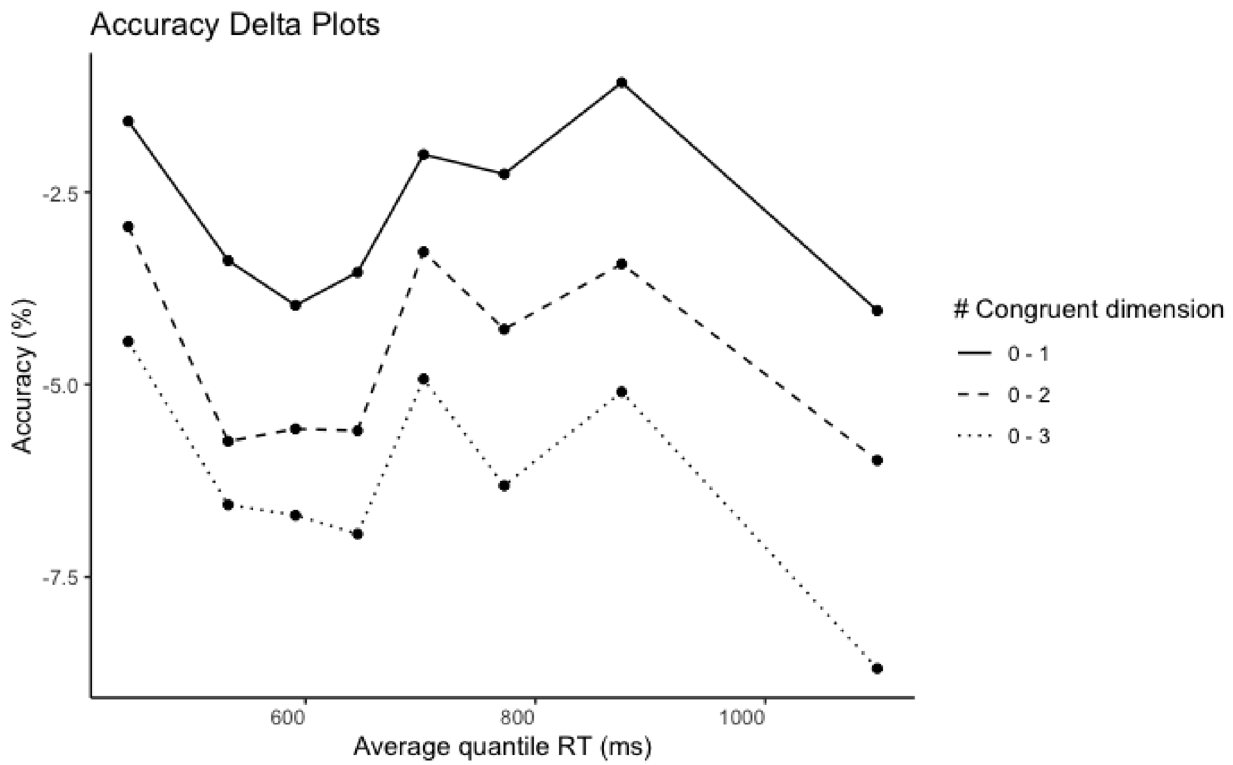

**Supplementary Figure 24.** Accuracy delta plots obtained contrasting congruency levels (zero vs. one, two, or three congruent dimensions) at each RT quantile.

The conditional accuracy function (the accuracy rate as a function of RT) suggests that most errors were either fast or slow responses. Importantly, only the intercept of the conditional accuracy function exhibited a linear increase as congruency levels rose. This is confirmed by overall flat accuracy delta plots when we contrasted congruency levels between each other. This pattern is compatible with a cumulative effect of interference (or facilitation) from independent non-cued dimensions.

### Reaction time

Next, we carried out the same distributional analyses on RT.

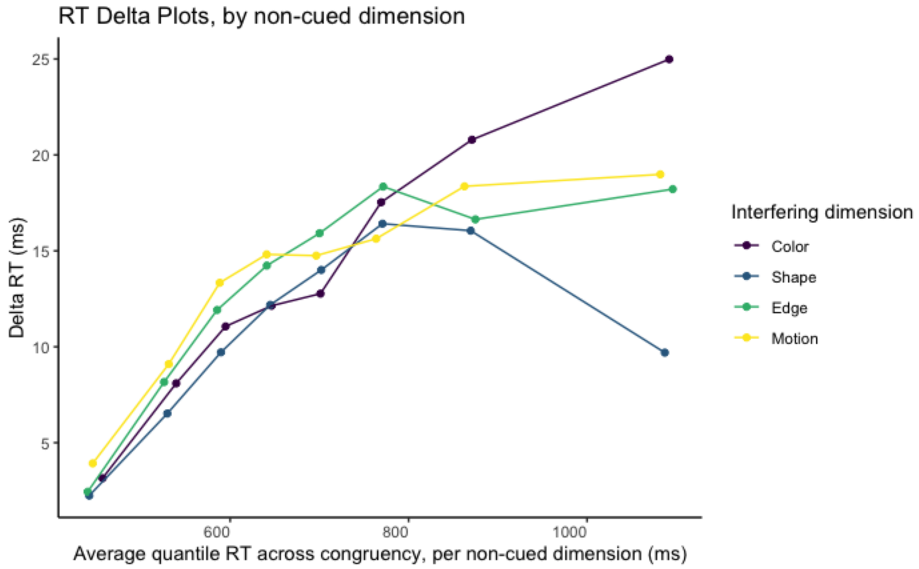

**Supplementary Figure 25.** Response time delta plots stratified by interfering dimension, obtained contrasting incongruent vs. congruent trials, at each RT quantile. RT quantile distribution were computed for each congruency level and interfering dimension. Data is plotted over the weighted RT mean across congruency level, but separately for each interfering dimension.

Similar to the accuracy delta plots, each non-cued dimension exhibited a unique temporal profile. The slopes during early responses suggested different strengths and timing of interference. Next, we explore how these independent dynamics are combined across dimensions to produce the cumulative interference time course.

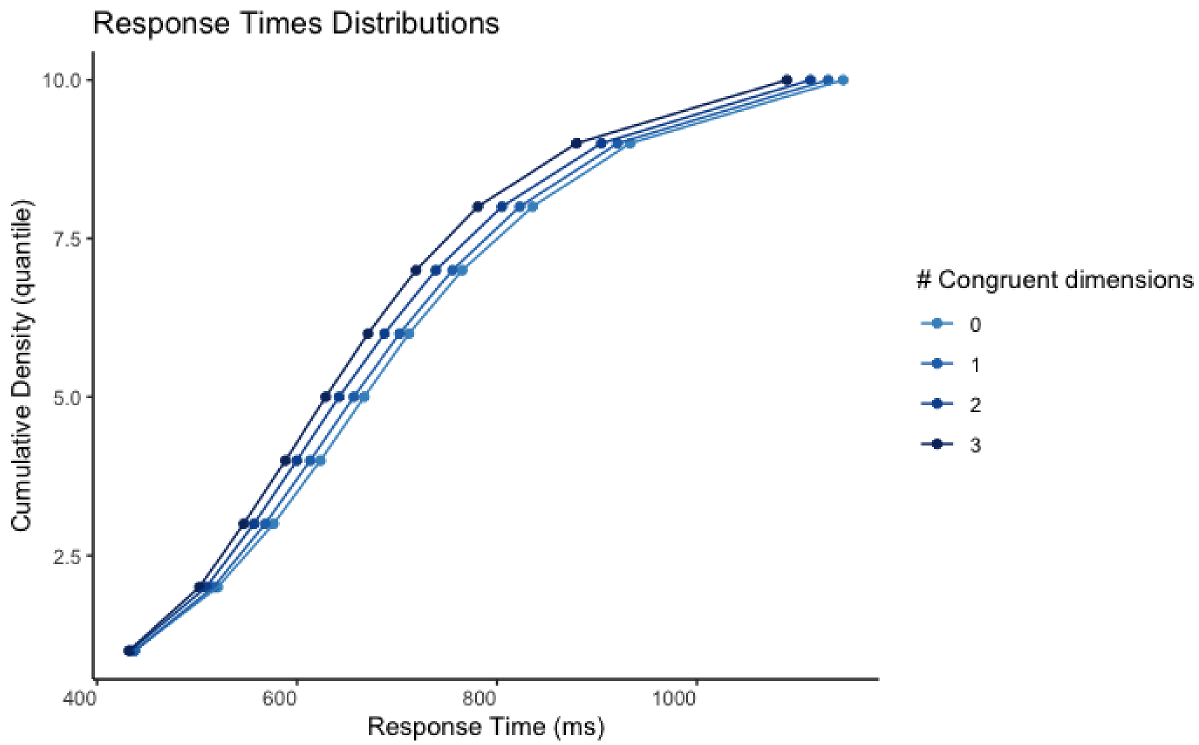

**Supplementary Figure 26.** Cumulative density distribution of response time. Data is stratified by parametric congruency.

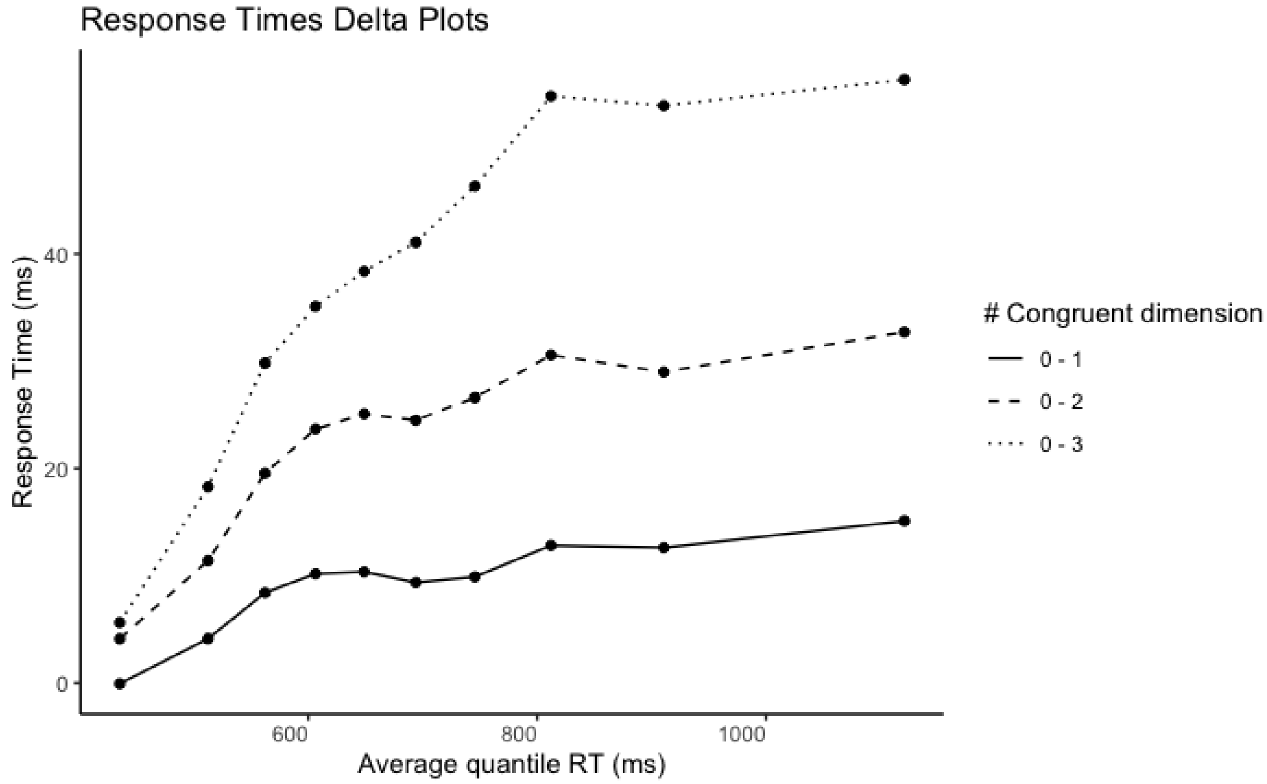

**Supplementary Figure 27.** Delta plots of interference effects obtained as RT difference between congruency levels (zero vs. one, two, or three congruent dimensions). Effects are shown over RT

quantiles, corresponding to the average RT of trials within each quantile and across congruency conditions, weighted by the corresponding trial count (note that the number of trials with congruency levels 1 and 2 is larger than with level 0 and 3).

In line with Experiment 1, congruency effects were the smallest for both early responses, but in this version of the task they were not suppressed at the end of the trial. The congruency effects increased till a plateau at around 800ms. Importantly, this plateau shifted linearly across congruency levels, again suggestive of a cumulative effect of interference (or facilitation) from independent non-cued dimensions.

**Responses are slower and less accurate after an error.**

We excluded the trials following incorrect responses from all analyses, to avoid contamination of conflict adaptation with other sources of adaptation, such as post-error slowing. Here we included these trials to highlight post-error slowing. Interestingly, at odds with Experiment 1, we observed decrements in accuracy after incorrect responses. This may suggest a global decrease of attention following errors.

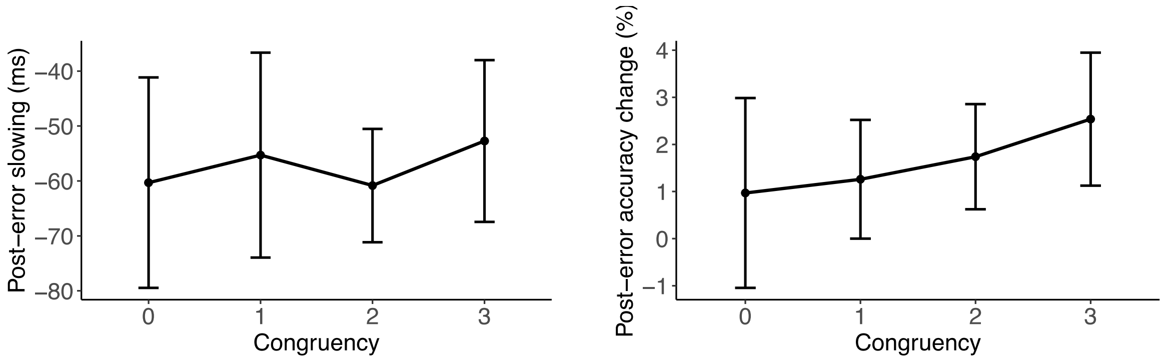

**Supplementary Figure 28.** RT (left) and accuracy (right) differences between trials following correct vs. incorrect responses. Error bars represent within-subject 95% confidence intervals.

### Experiment 3

Here, we report the results of Experiment 3 in more detail.

#### The number of congruent dimensions predicts task performance

As in Experiments 1 and 2, we found that the number of congruent dimensions predicted both RT and accuracy. In particular, we observed a negative linear effect of congruency on RTs (Mdn = -0.03, 95% CI = [-0.04, -0.03], $ER_{congruency<0}$ = Inf), and a positive linear effect of congruency on accuracy (Mdn = 0.66, 95% CI = [0.62, 0.69], $ER_{congruency>0}$ = Inf). Participants were fastest when the stimulus was fully congruent, and became slower with each decrease in the number of congruent dimensions ($M_0$ = 701 ms, CI = [682, 719]; $M_1$ = 683 ms, CI = [665, 701]; $M_2$ = 665 ms, CI = [648, 684]; $M_3$ = 648 ms, CI = [630, 666]; $ER_{0>1}$ = 11.4, $ER_{1>2}$ = 10.6, $ER_{2>3}$ = 10.0). Analogously, participants showed highest accuracy for fully congruent stimuli, which decreased with each increase in the number of congruent dimensions ($M_0$ = 87.1%, CI = [86, 89]; $M_1$ = 92.8%, CI = [92, 94]; $M_2$ = 96.2%, CI = [96, 97]; $M_3$ = 98.0%, CI = [98, 98]; $ER_{0>1}$ = Inf, $ER_{1>2}$ = Inf, $ER_{2>3}$ = Inf).

A negative Pearson's correlation between the random effects of congruency on RT and accuracy ($r$ = -0.39, $p$ < 0.001, 95% confidence intervals = [-0.54 -0.21]) confirmed a subject-level association between the two measures.

#### Each stimulus dimension produces interference

All non-cued dimensions produced interference for both RT (color: BF = $4.9\times10^{19}$, shape: BF = $7.7\times10^{15}$, edge: BF = $2.7\times10^{15}$, including the new dimension dot proportion: BF = $5.4\times10$), and accuracy (color: BF = $1.2\times10^{22}$, shape: BF = $3.4\times10^{22}$, edge: BF = $2.2\times10^{28}$, dot proportion: BF = $3.4\times10^{19}$). In fact, we observed reliable interference from any non-cued dimension on any cued task (Figure 7BE) for accuracy (min BF = $2.1\times10^{11}$, max BF = $6.2\times10^{25}$). For RT, the interference of dot proportion on edge was not supported (BF = 0.08), while there was strong evidence for every other effect (min BF = $1.5\times10$, max BF = $4.9\times10^{17}$).

On average between 3 and 4 non-cued dimensions induced positive interference effect (RT: M = 3.20, SD = 0.77; accuracy: M = 3.81, SD = 0.42). If only one dimension provided interference per participant, we would expect 2.5 positive effects on average (1 from the single non-cued dimension that drove the effect, and 1.5 from 3 non-cued dimensions hovering around zero). Inconsistent with this hypothesis, the average number of positive interference effects was higher than this value for both RT ($BF_{(m>2.5\ /\ m<2.5)} = 4.7\times10^{31}$) and accuracy ($BF_{(m>2.5\ /\ m<2.5)} = 1.8\times10^{86}$).

#### Interference effects persist over time

To test persistence of interference, we computed mean interference effects for each non-cued dimension as a function of the number of blocks that passed since it was the cued dimension (Supplementary Figure 29 & 30). Interference effects were largest on the first block after the switch, but remained reliable until block five (i.e., ~20 trials) for both RT (range BF: min = $4.8\times10$, max = $1.1\times10^{29}$) and accuracy (range BF: min = $1.6\times10^{5}$, max = $1.0\times10^{32}$).

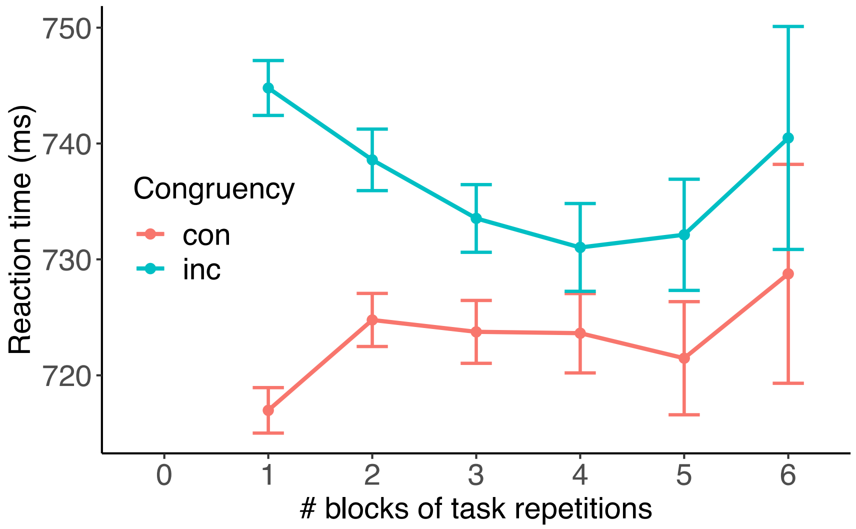

**Supplementary Figure 29.** In Experiment 3, the interference effect, measured in response time, of a non-cued dimension persists for six blocks after it was the cued dimension.

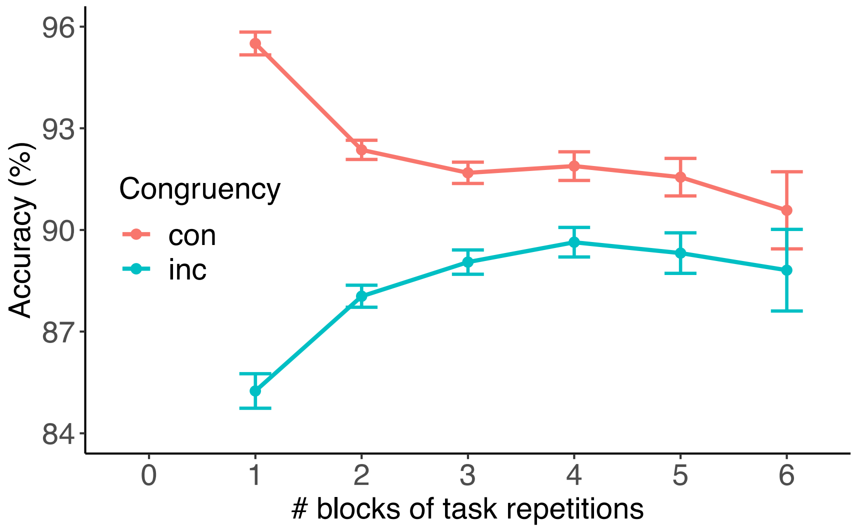

**Supplementary Figure 30.** In Experiment 3, the interference effect, measured in accuracy, of a non-cued dimension persists for six blocks after it was the cued dimension.

**Conflict adaptation is modulated by the degree of prior congruency**

The congruency sequence effects observed in Experiments 1 and 2 were replicated with the continuous-objects version of the task.

For RTs, we observed strong evidence in favor of a main effect of congruency (Mdn = -0.02, 95% CI = [-0.02, -0.01], $ER_{cong.<0}$ = Inf.), accounting for a very robust decrease of RTs with increasing congruency, conditional on previous congruency set to 0 ($M_{00}$ = 689 ms, CI = [670, 707]; $M_{03}$ =

662 ms, CI = [643, 680]). We observed strong evidence for a main effect of previous congruency (Mdn = 0.02, 95% CI = [0.01, 0.02], $ER_{prev\_cong.>0}$ = Inf.), accounting for increasing RTs with increasing previous congruency, conditional on current congruency set to 0 ($M_{00}$ = 689 ms, CI = [670, 707]; $M_{30}$ = 716 ms, CI = [696, 736]). Importantly, we observed strong evidence in favor of an interaction effect between current and previous congruency (Mdn = -0.01, 95% CI = [-0.01, -0.01], $ER_{cong.:prev\_cong.<0}$ = Inf), accounting for larger effects of current congruency on RTs with increasing number of previous congruent dimensions.

As can be seen from the marginal means for the 16 levels of this interaction (Supplementary Figure 31), the effect of current congruency was strongest after fully congruent trials, and then parametrically declined as prior congruency decreased. Indeed, even though we observed strong evidence in favor of an effect of congruency when the previous trial was fully incongruent (ER = Inf, $M_{00}$ = 689 ms, $M_{03}$ = 662 ms), the effect of congruency after fully congruent trials was even larger (ER = Inf, mean $M_{30}$ = 716 ms, $M_{33}$ = 637 ms). In other words, compared to trials following fully incongruent trials, trials following fully congruent trials showed strong interference when current congruency was low (leading to longer RTs; $M_{30-00}$ = 27 ms, CI = [0, 54]) and strong facilitation when current congruency was high (leading to shorter RTs; $M_{33-03}$ = -25 ms, CI = [-52, 0]). Compared to experiments one and two, in experiment three the interference effect was relatively smaller than the facilitation effect.

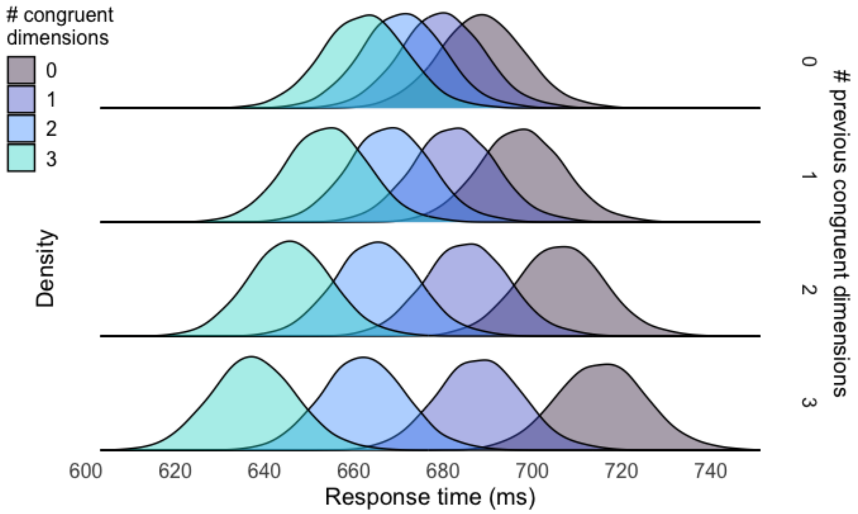

**Supplementary Figure 31.** Posterior distributions of the marginal means from a Bayesian linear model explaining response time as a function of both current (colors) and previous (rows) congruency in Experiment 3.

Analogous congruency sequence effects were observed for accuracy. At difference with RTs, a strong main effect of congruency (Mdn = 0.47, 95% CI = [0.40, 0.54], $ER_{cong.>0}$ = Inf.) indicated that accuracy increased with congruency even when holding previous congruency to level 0 ($M_{00}$ = 92.5%, CI = [91, 94]; $M_{03}$ = 98.0%, CI = [98, 99]). In line with RTs, we observed strong evidence for a main effect of previous congruency (Mdn = -0.37, 95% CI = [-0.41, -0.32], $ER_{prev\_cong.<0}$ = Inf), accounting for decreasing accuracy with increasing previous congruency, conditional on current congruency set to 0 ($M_{00}$ = 92.5%, CI = [91, 94]; $M_{30}$ = 80.4%, CI = [78, 83]). Importantly, there was strong evidence for an interaction between current and previous congruency (Mdn = 0.11, 95% CI = [0.08, 0.14], $ER_{cong.:prev\_cong.>0}$ = Inf) indicated by a larger effect of congruency with increasing numbers of previous congruent dimensions. Also here, there was a strong effect of congruency even when the previous trial was fully incongruent (ER = Inf, $M_{00}$ = 92.5%, $M_{03}$ = 98.0%), but this effect was larger after fully congruent trials (ER = Inf, mean $M_{30}$ = 80.4%, $M_{33}$ = 97.9%). We observed strong interference effects on accuracy ($M_{30-00}$ = -12.1%, CI = [-15, -10]) but, at difference with RTs, no facilitation effects ($M_{33-03}$ = -0.16%, CI = [-0, 0]), the latter likely due to a ceiling accuracy for fully congruent trials (Supplementary Figure 32, teal distributions).

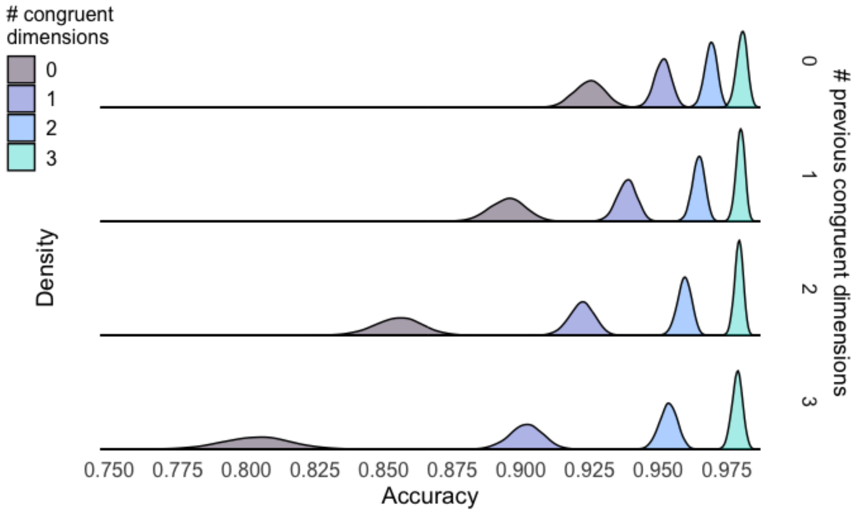

**Supplementary Figure 32.** Posterior distributions of the marginal means from a Bayesian linear model explaining accuracy as a function of both current (colors) and previous (rows) congruency in Experiment 3.

#### Conflict adaptation weakens over time and resets after a task switch

When we analyzed adaptation effects on switch trials alone (Supplementary Figure 33AB), using the model in Equation 2, we observed strong evidence against an interaction between current and previous congruency for both RT (Mdn = $-8.83\times10^{-4}$, 95% CI = [-0.00, 0.00], $ER_{cong.:prev\_cong.=0}$ = 1389) and accuracy (Mdn = $-5.85\times10^{-3}$, 95% CI = [-0.05, 0.04], $ER_{cong.:prev\_cong.=0}$ = 124). This suggests that control history effects do not survive a task switch, with control settings 'resetting' upon encountering new task demands.

Last, we tested whether congruency sequence effects persist over multiple trials. We modeled RTs and accuracy adapting Equation 2, where *previous_congruency* now specifies the congruency level in the 2nd-to-last or 3rd-to-last trial. First, we tested whether the number of congruent non-cued dimensions experienced two trials ago modulated the congruency effect on the current trials (Supplementary Figure 33CD). We observed strong evidence for such adaptation in RT (Mdn = $-5.90\times10^{-3}$, 95% CI = [-0.01, 0.00], $ER_{cong.:prev\_2\_cong.<0}$ = $1.19\times10^{4}$) and moderate evidence for accuracy (Mdn = 0.02, 95% CI = [-0.01, 0.05], $ER_{cong.:prev\_2\_cong.>0}$ = 6). Next, we tested the CSE on the third-next trial (Supplementary Figure 33EF). At difference with experiment one, and in line with experiment two, congruency experienced three trials ago did not reliably modulated congruency effects on the current trial (RTs: Mdn = $-4.75\times10^{-4}$, 95% CI = [0.00, 0.00],

$ER_{cong.:prev\_3\_cong.<0}$ = 1.6; accuracy: Mdn = $-4.93\times10^{-3}$, 95% CI = [-0.04, 0.03], $ER_{cong.:prev\_3\_cong.>0}$ = 0.6).

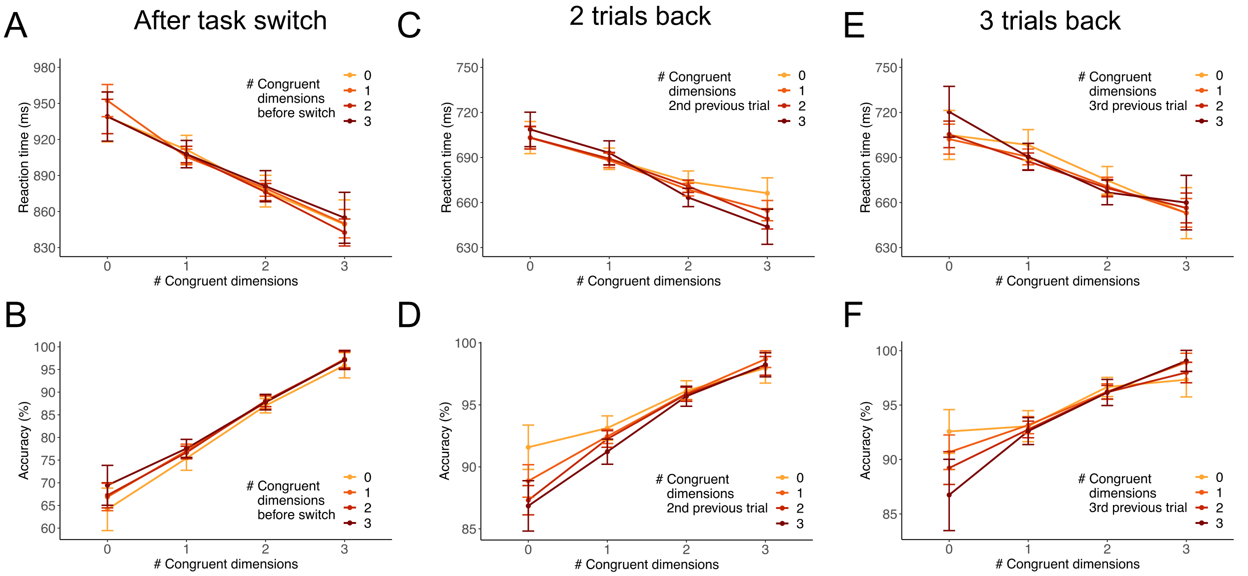

**Supplementary Figure 33.** Congruency sequence effects over time and after task switches in Experiment 3. We found that there was no conflict adaptation after task switches for (A) RT or (B) accuracy. However, we found that conflict from (CD) 2 trials back reliably induced adaptation for both measures. There was no evidence of adaptation based on conflict from 3 trials back (EF).

**Conflict adaptation is selectively driven by sequence effects within the same dimension**

The main text reports the results of the trial-average approach, where we replicate that adaptation effects were driven by sequences within dimension (on-diagonal, Figure 7GI), and not across dimension (off-diagonal).

We then modeled the adaptation effects at the single trial level, to test for cumulative effects of multiple non-cued dimensions acting in concert in any given trial. Congruency sequences within and across dimensions were modeled separately. The estimated marginal means from the two models are reported in Supplementary Figure 34. The cumulative effect of each numeric predictor is shown assuming a single CS instance (i.e., each numeric predictor is set to 1). When considering currently incongruent non-cued dimensions (red distributions), we observed slower RTs in trials following a congruent vs. incongruent non-cued dimension of the *same* type (CI==1 vs. II==1, both implying CC==2; within effect: $M_{CI-II}$ = 8 ms, 95% CI = [5, 12], $ER_{CI-II>0}$ = Inf.), while there was no effect when comparing across dimensions sequences (across effect: $M_{CI-II}$ = -0 ms, 95% CI = [-5, 4], $ER_{CI-II>0}$ = 0.8). We also observed slower RTs for currently congruent dimensions (blue distributions) following the *same* dimension being incongruent in the previous trial (IC==1 with CC==2), as compared to the intercept value (CC==3; within effect: $M_{IC-Int}$ = 8 ms, 95% CI = [4, 12], $ER_{IC-Int>0}$ = $3.1\times10^{34}$), suggesting a facilitation effect driven by previous congruency in the same dimension (CC sequences). No evidence for this effect was observed also when considering across dimensions sequences (across effect: $M_{IC-Int}$ = 0 ms, 95% CI = [-5, 5], $ER_{IC-Int>0}$ = 1.1). Therefore, the overall pattern of results was compatible with experiments one and two.

In other words, we observed adaptation effects (including facilitation and interference) selectively when considering within-dimension sequences.

| Contrast | Evidence ratio | Mean difference | HDI low | HDI high |
|---|---|---|---|---|
| CI-II>0 - within | Inf | 8.45 | 5.03 | 11.76 |
| CI-II>0 - across | 0.87 | -0.19 | -4.84 | 4.28 |
| IC-Int>0 - within | 31999.00 | 8.45 | 4.29 | 12.39 |
| IC-Int>0 - across | 1.09 | 0.15 | -4.74 | 5.03 |

**Supplementary Table 5.** RT contrasts within and across dimensions.

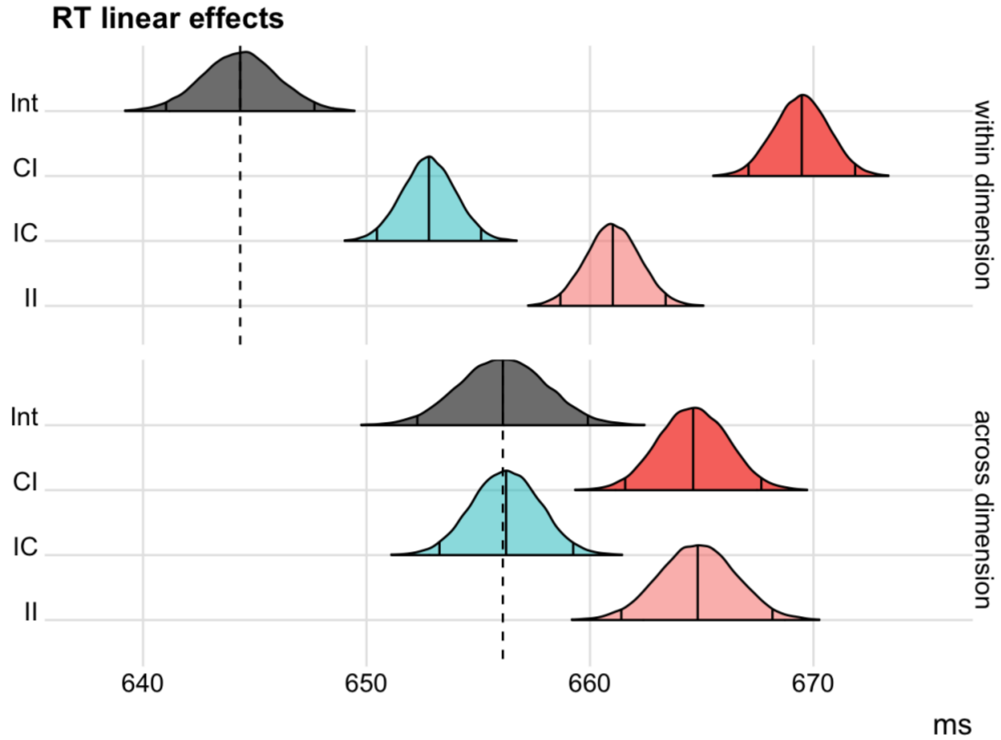

**Supplementary Figure 34.** Posterior distributions of the marginal means from two Bayesian linear models explaining RT adaptation effects in Experiment 3 as a function of either within-dimension conflict sequences (top row), or across-dimension conflict sequences (bottom row). The vertical solid lines represent the median and equal-tailed 95% credible intervals of the posterior samples. The intercept distributions (grey) correspond to the maximum number of CC sequences (within: 3; across: 6), and no instance of any other type. The other distributions represent the marginal means when introducing a single instance of the corresponding sequence type.

Analogous results were observed when predicting accuracy with the same approach (Supplementary Figure 35). When turning our attention to across-dimensions sequences, we observed a reliable main effect of previous congruency, with lower accuracy after trials including additional congruent dimensions ($M_{CI-II}$ = -0.37 %, 95% CI = [-0.89, 0.15], $ER_{CI-II<0}$ = 11.3; $M_{Int-IC}$ = -0.63 %, 95% CI = [-1.07, -0.20], $ER_{Int-IC<0}$ = 749). In other words, when considering across-dimension sequences, the main effect of previous congruency suggests that accuracy increased if there were more incongruent dimensions in the previous trial.

| Contrast | Evidence ratio | Mean difference | HDI low | HDI high |
|---|---|---|---|---|
| CI-II<0 within | Inf | -1.70 | -2.05 | -1.36 |
| IC-Int<0 within | 17.93 | -0.20 | -0.44 | 0.04 |
| CI-II<0 across | 11.37 | -0.37 | -0.89 | 0.15 |
| Int-IC<0 across | 749.00 | -0.63 | -1.07 | -0.20 |

**Supplementary Table 6.** Accuracy contrasts within and across dimensions.

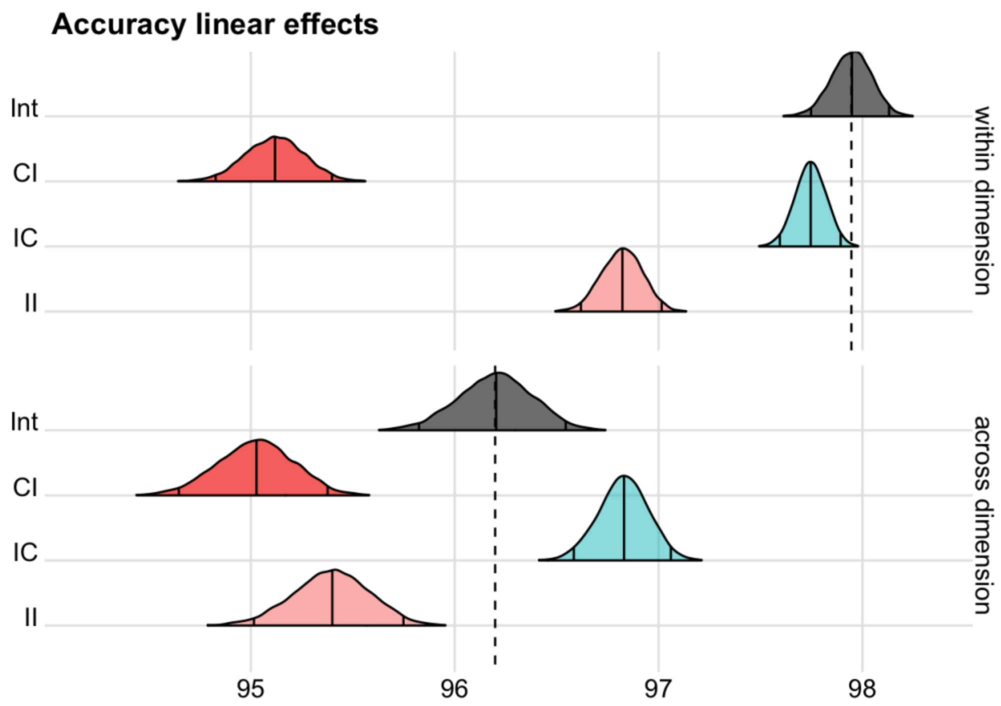

**Supplementary Figure 35.** Posterior distributions of the marginal means from two Bayesian linear models explaining accuracy adaptation effects in Experiment 3 as a function of either within-dimension conflict sequences (top row), or across-dimension conflict sequences (bottom row). The vertical solid lines represent the median and equal-tailed 95% credible intervals of the posterior samples. The intercept distributions (grey) correspond to the maximum number of CC sequences (within: 3; across: 6), and no instance of any other type. The other distributions represent the marginal means when introducing a single instance of the corresponding sequence type.

To rule out that each participant only applied within-dimension adaptation for one dimension, we computed participant's average number of positive within-dimension effects for both RT (RT: M = 2.87, SD = 0.90) and accuracy: (M = 3.18, SD = 0.87). Compared to experiments one and two, a lower number of positive adaptation effects was due to the weaker interference exerted by the new dimension 'dot proportion'. Nevertheless, we found strong evidence that these numbers were larger than the 2.5 positive adaptation effects one would expect under the alternative explanation (RT: $BF_{(m>2.5\ /\ m<2.5)} = 4.4\times10^{10}$, Accuracy: $BF_{(m>2.5\ /\ m<2.5)} = 1.4\times10^{21}$).

**Dimensions' processing pathways are independent**

To demonstrate that the stimulus processing pathways of the MULTI are independent, we sought to rule out the existence of interactive effects between non-cued dimensions' interference effects. First, we subset the data based on which dimension was cued. For each subset, we ran a linear model predicting RTs including the main effects of congruency for each of the three non-cued dimension, and the interactions between them:

RT(cued_dim) ~ 1+ congruency_1 * congruency_2 * congruency_3 + (1 + … | subject) where 1 refers to the intercept, and congruency_x is a categorical predictor of congruency for a non-cued dimension. All main effects and interactions were also added as random effects.

Since the main effects of congruency of each non-cued dimension have been already reported, here we only tested their interaction effects against the point-null hypothesis. We found very strong evidence against interaction effects between every pair of non-cued dimensions, and across all of the four models run for each cued task (range $ER_{cong\_x:cong\_y=0}$: min = 30, max = 218, median = 135). This result confirms that dimensions' interference effects do not interact. Rather, they show that RTs depend on the linear sum of each dimension's congruency effects.

#### Distributional analyses

To investigate the temporal dynamics of dimensions' interference, here we report distributional analyses on accuracy and RT. In line with Experiment 1, the results of these analyses suggest that the time course of cumulative interference (and suppression thereof) results from the linear combination of the single dimensions' dynamics.

#### Accuracy

We carried out distributional analyses of accuracy rates as a function of RT quantiles. First, we quantified the conditional accuracy function stratifying data by interfering (non-cued) dimension.

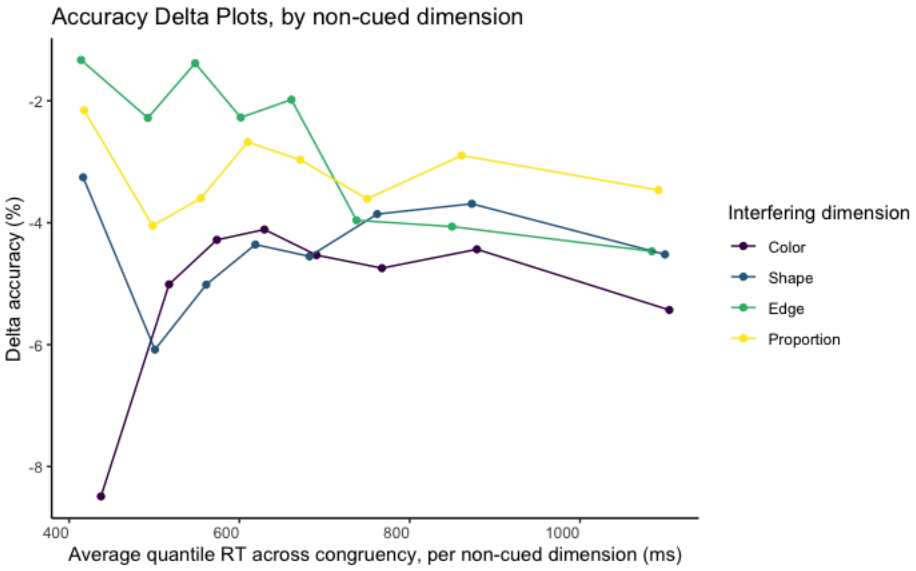

**Supplementary Figure 36.** Accuracy delta plots stratified by interfering dimension, obtained from contrasting incongruent vs. congruent trials, at each RT quantile. The RT quantile distributions were computed for each congruency level and interfering dimension. Data is plotted over the weighted RT mean across congruency levels, but separately for each interfering dimension. To note, each non-cued dimension exhibits a unique temporal profile. For example, the interfering dimension color is the only one that produced a strong initial attentional capture (high rate of fast errors for incongruent vs. congruent color), followed by a stabilization of such interference. The interference of the other dimensions shows a gradual temporal increase, with varying levels of late suppression.

The idiosyncratic nature of the temporal dynamic of dimension interference is reminiscent of the different magnitudes of their congruency effects. Next, we explore how these independent dynamics are combined across dimensions to produce the cumulative interference time course. To note, each dimension was cued (or non-cued) for an equal number of trials.

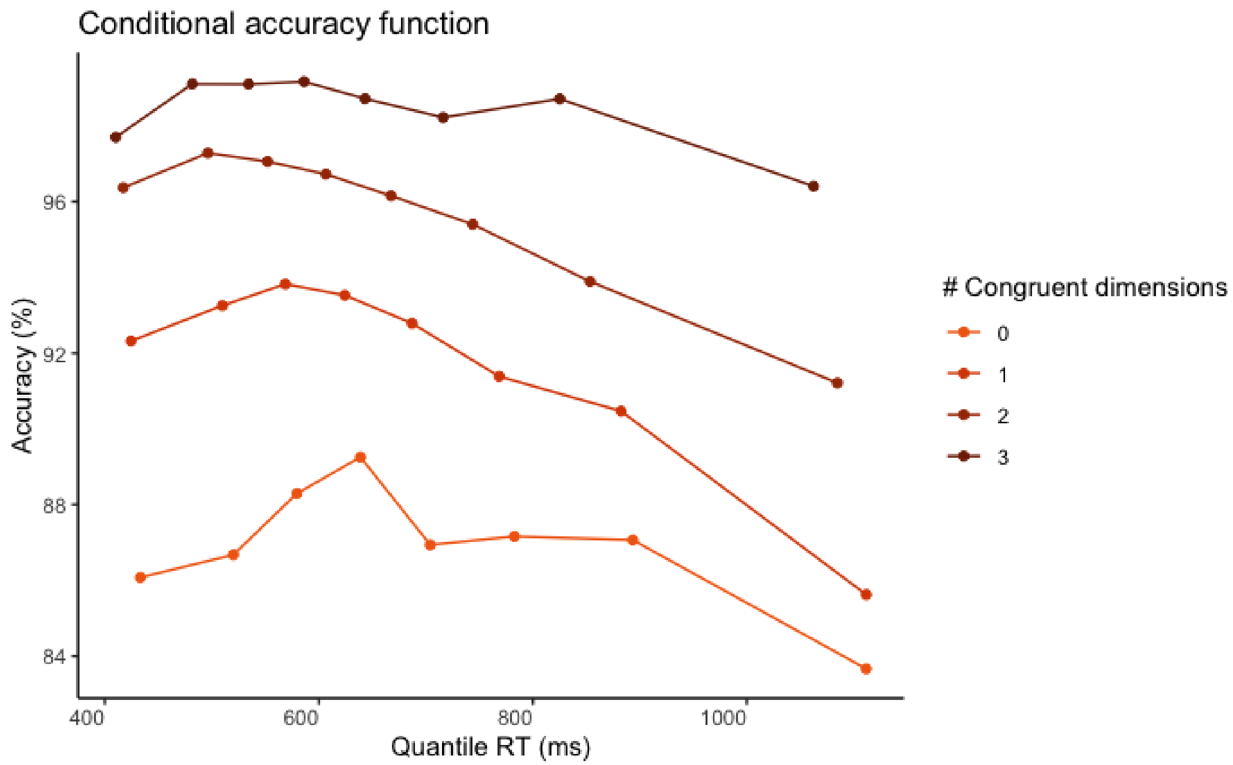

**Supplementary Figure 37.** Conditional accuracy function. Data is stratified by parametric congruency. The accuracy rate is computed separately for each RT quantile and congruency level.

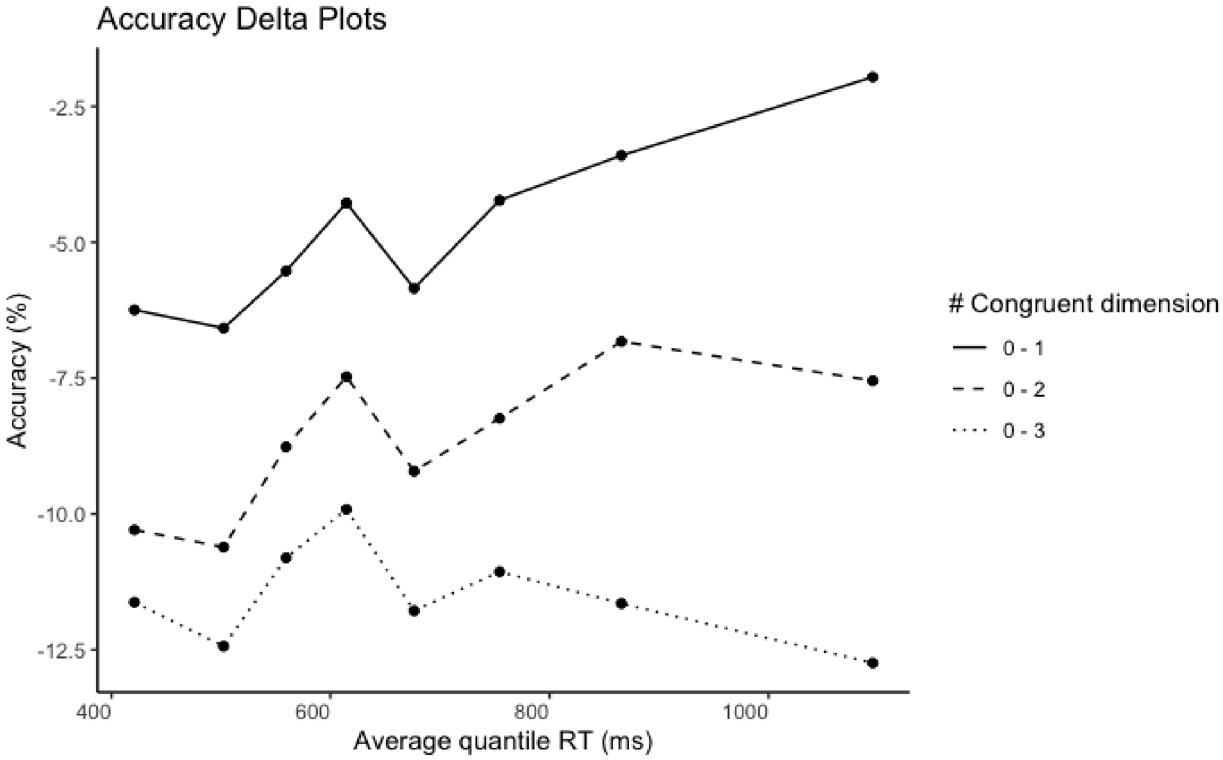

**Supplementary Figure 38.** Accuracy delta plots obtained contrasting congruency levels (zero vs. one, two, or three congruent dimensions) at each RT quantile.

The conditional accuracy function (the accuracy rate as a function of RT) suggests that most errors were either fast or slow responses. Importantly, only the intercept of the conditional accuracy function exhibited a linear increase as congruency levels rose. This is confirmed by overall flat accuracy delta plots when we contrasted congruency levels between each other. This pattern is compatible with a cumulative effect of interference (or facilitation) from independent non-cued dimensions.

**Reaction time**

Next, we carried out the same distributional analyses on RT.

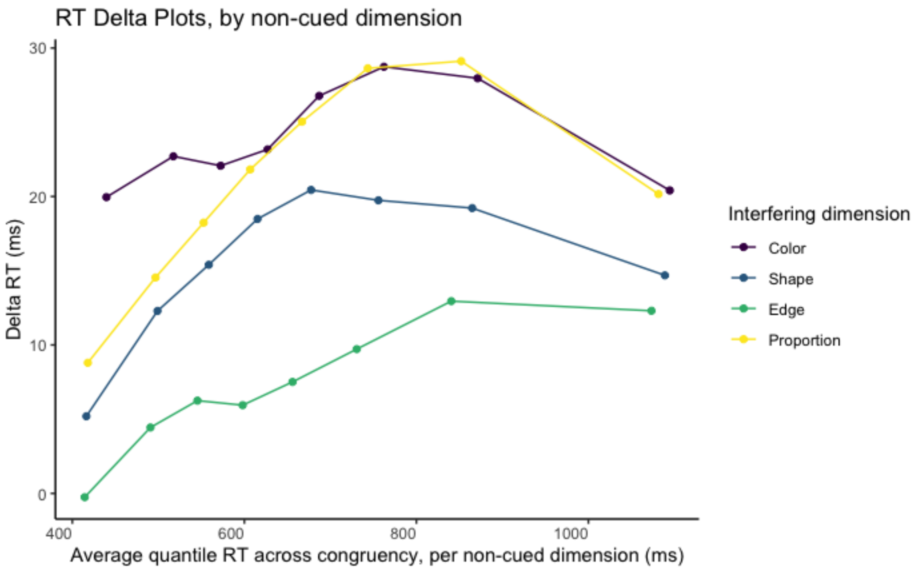

**Supplementary Figure 39.** Response time delta plots stratified by interfering dimension, obtained contrasting incongruent vs. congruent trials, at each RT quantile. RT quantile distribution were computed for each congruency level and interfering dimension. Data is plotted over the weighted RT mean across congruency level, but separately for each interfering dimension.

Similar to accuracy delta plots, each non-cued dimension exhibited a unique temporal profile. The slopes during early responses suggested different strengths and timing of interference. Next, we explore how these independent dynamics are combined across dimensions to produce the cumulative interference time course.

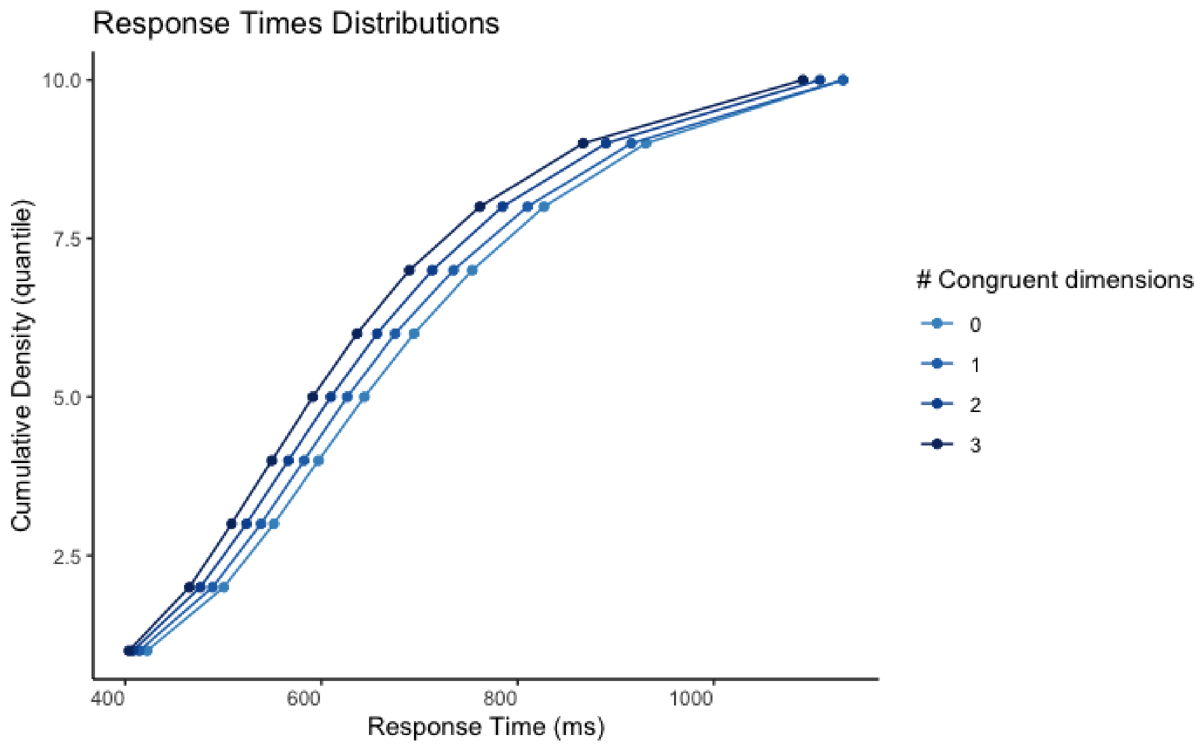

**Supplementary Figure 40.** Cumulative density distribution of response time. Data is stratified by parametric congruency.

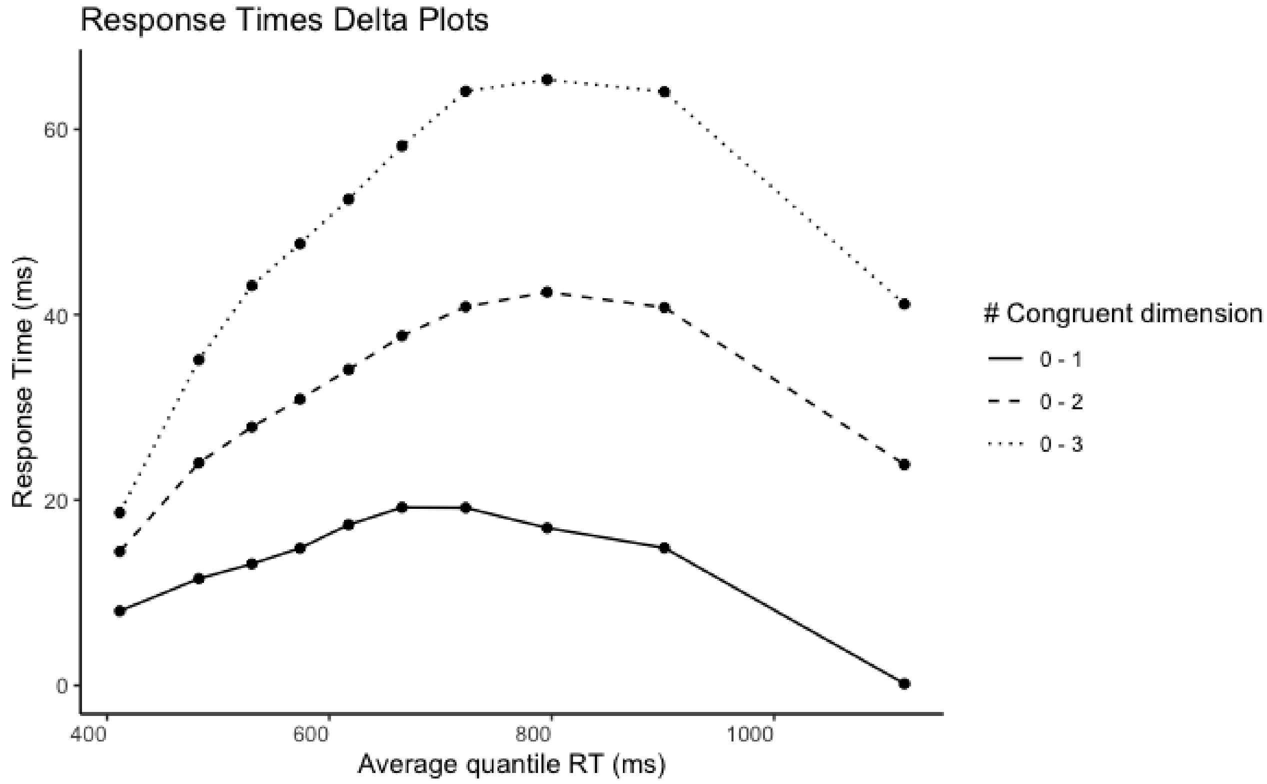

**Supplementary Figure 41.** Delta plots of interference effects obtained as RT difference between congruency levels (zero vs. one, two, or three congruent dimensions). Effects are shown over RT

quantiles, corresponding to the average RT of trials within each quantile and across congruency conditions, weighted by the corresponding trial count (note that the number of trials with congruency levels 1 and 2 is larger than with level 0 and 3).

Similarly to Experiment 1, congruency effects were the smallest for both early and late responses. The congruency effects increased and decreased within the RT range, with a peak time of around 800ms. Importantly, this peak increased linearly across congruency levels, again suggestive of a cumulative effect of interference (or facilitation) from independent non-cued dimensions.

**Responses are slower and less accurate after an error.**

We excluded the trials following incorrect responses from all analyses, to avoid contamination of conflict adaptation with other sources of adaptation, such as post-error slowing. Here we included these trials to highlight post-error slowing. In line with Experiment 2, we also observed lower accuracy following errors, suggesting global decreases in attention.

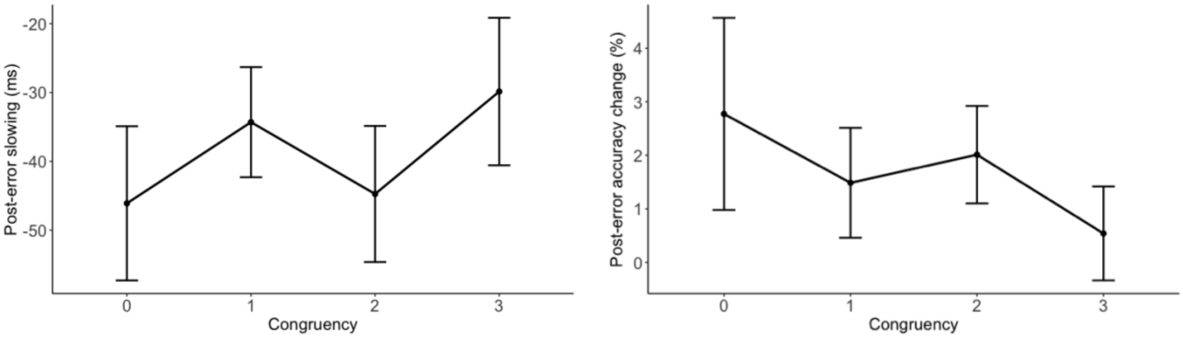

**Supplementary Figure 42.** RT (left) and accuracy (right) differences between trials following incorrect vs. correct responses. Error bars represent within-subject 95% confidence intervals.

### Target-signal sequential effects

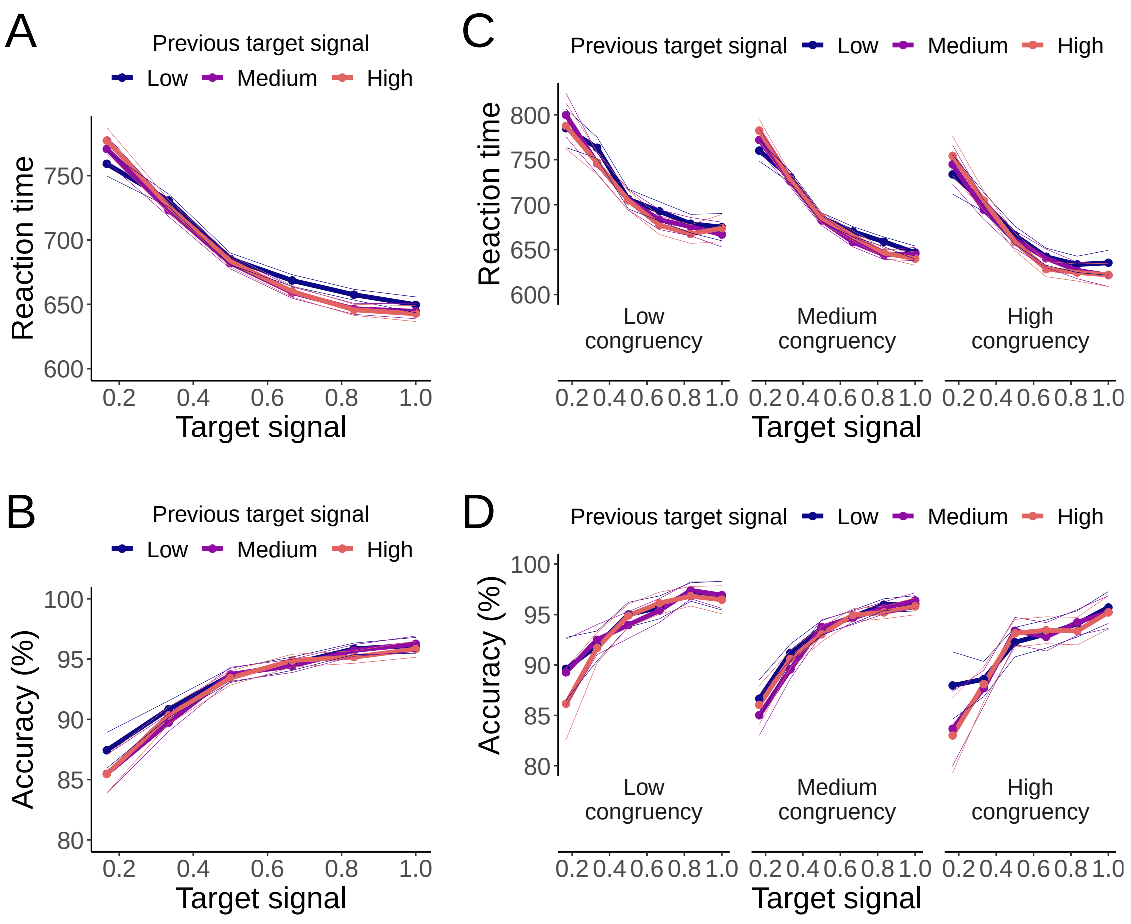

**Supplementary Figure 43. Experiment 3 (N = 103).** Signal detection analyses on RT (top row) and accuracy (bottom row). A low target signal in the previous trial was associated with slower RTs (a) and higher accuracy (b). This pattern of results reflects speed/accuracy trade-off and suggests that low discriminability of the cued dimension can elicit response caution. These effects were independent of the congruency of non-cued dimensions (c,d).

#### A note on post-conflict caution

When we predicted performance on the single-trial level, and classified trials based on the presence and number of each sequence type, we confirmed that conflict adaptation was selectively driven by within-dimension sequences. This second approach recapitulated varying magnitudes of facilitation vs. inhibition effects across the three experiments, which were already observed in terms of congruency sequence effects (see Fig. 2-3).

However, when we turned our attention to across-dimension behavioral effects, we observed clear main effects of congruency, with more accurate and faster responses for current congruency, but intriguingly, we also noticed main effects of previous congruency, with performance following a trial with one more incongruent dimension being more accurate (and also slower, in Experiment 3) than when following a trial with one more congruent dimension. This pattern of behavior, observed only for across-dimension sequences (which were not affected by conflict adaptation effects), suggests some subtle but reliable effect of target adaptation, in combination with a general post-conflict increase in response caution, which has been previously hypothesized (Verguts et al., 2011) but rarely observed in classic control paradigms. Specifically, heightened response thresholds are needed to explain increased accuracy in absence of corresponding faster RTs.

### Supplementary Simulations

#### Model simulations of two-dimensional interference paradigms

To demonstrate continuity across tasks and experimental effects, we simulate the standard Gratton effect by reducing the processing pathways of the model from four to two (i.e., a target and a distractor dimension, as in typical interference tasks). The code used to generate these simulations allows to define any arbitrary number of stimulus dimensions. Here, the distractor-specific adaptation algorithm was used. Comparable results can be obtained with a global distractor adaptation, but not with target adaptation (see Results and Discussion).

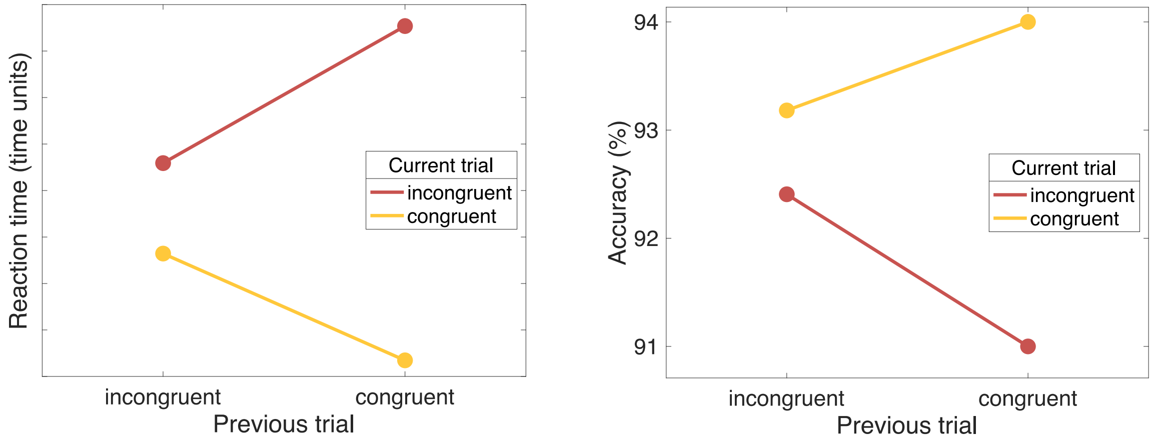

**Supplementary Figure 44.** Simulation results obtained with a classic two-dimensional model architecture. To model information processing of typical two-dimensional stimuli used in classic interference paradigms, corresponding to a task-relevant and a task-irrelevant dimension, only two feed-forward pathways were used (cf. Figure 5A). The parametric congruency sequence effects observed in our experiments (Figure 7CF) reduce to classic adaptation effects: the main effect of current trial congruency is modulated by congruency on the previous trial.

#### The model reproduces essential behavior across a broad parameter space

To demonstrate the robustness of the model to in a broad parameter space, we performed grid-search across the hand-tuned parameters. These include the negative bias to the hidden units (ranging -10:1:0) and task-to-hidden weights (ranging 0:1:10). For each tile of the parameter space, we repeated the simulation of 1000 synthetic participants, each of which ran through the same number of trials as in our experiments (1600 trials).

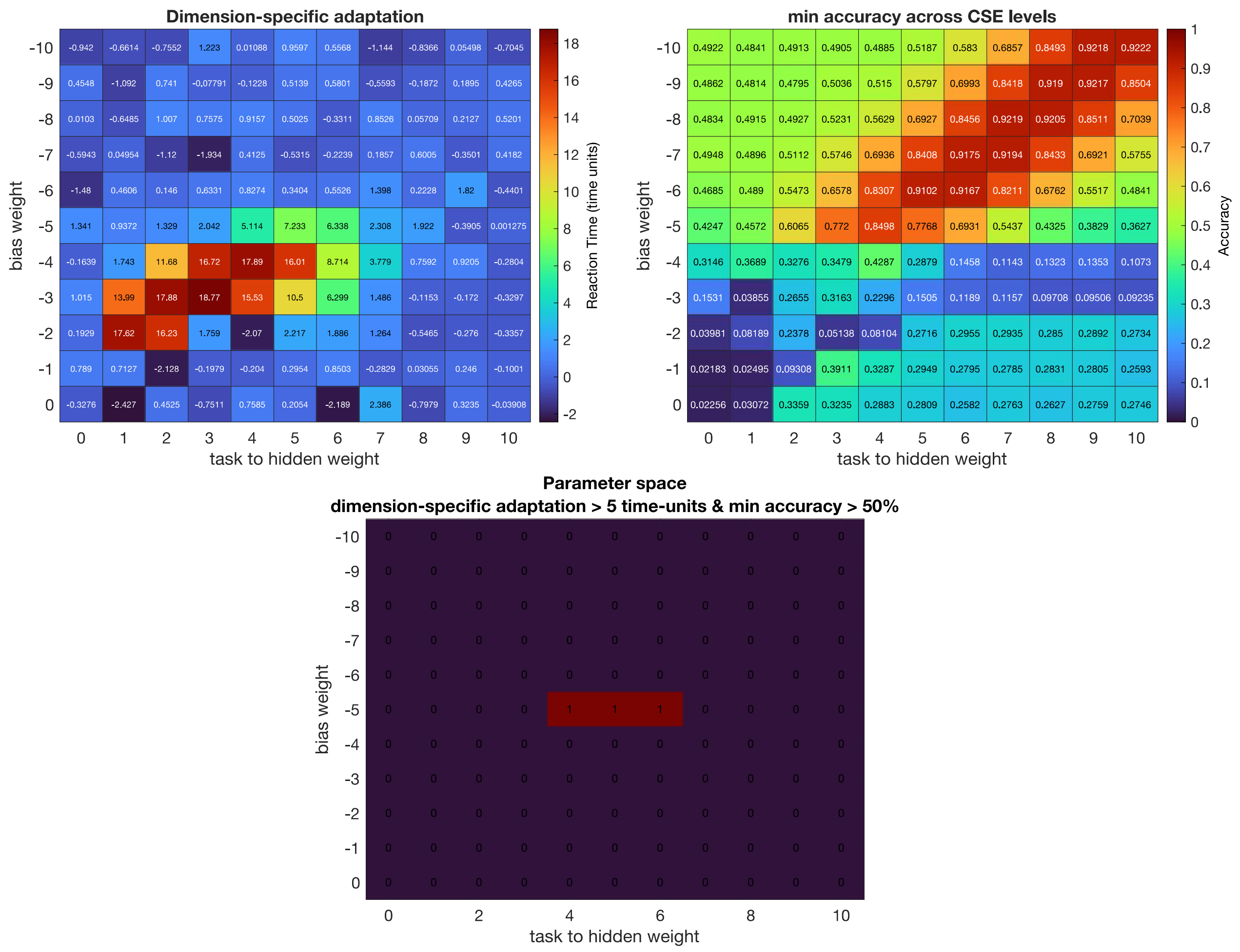

**Supplementary Figure 45.** Grid search results obtained using the dimension-specific adaptation algorithm. **a,** Specificity of adaptation effects obtained subtracting across-dimension adaptation effects from within-dimension effects. **b,** Lowest accuracy value across the 16 levels of the parametric congruency sequence analysis (cf. Figures 2CF, 4CF, 7CF). **c,** Conjunction of grids from (a) and (b), obtained by imposing specificity effect larger than 5 time units, and lowest accuracy above 50%. Red tiles indicate well-calibrated weights.

The grid-search results indicate that the distractor-specific model captures dimension-specific adaptation effects in a relatively large swath of parameter space (panel A, warm colors). To ensure that simulations qualitatively reproduce essential behavior, the selection of parameters is further constrained by observed accuracy levels (panel B), which was never lower than 50% in any of the congruency sequence conditions (cf. Figures 2CF, 4CF, 7CF). In comparison, simulations using global distractor adaptation and target adaptation did not show any dimension-specific adaptation effects across the full parameter space.

### R packages

We used the following R packages: Brms (Bürkner, 2017), Ggplot2 (Wickham, 2010), Gridges (Wilke, 2018), and Tidyverse (Wickham, 2017).
